# Survival and Spread of Engineered *Mycobacterium smegmatis* and Associated Mycobacteriophage in Soil Microcosms

**DOI:** 10.1101/2025.01.27.635130

**Authors:** Megan S. Fleeharty, Kate B.R. Carline, Bilalay V. Tchadi, Bjorn B. Shockey, Emma C. Holley, Margaret S. Saha

## Abstract

The inoculation of microbes into soil environments has numerous applications for improving soil quality and crop health; however, the ability of exogenous and engineered microbes to survive and spread in soil remains uncertain. To address this challenge, we assayed the survival and spread of *Mycobacterium smegmatis*, engineered with either plasmid transformation or genome integration, as well as its mycobacteriophage Kampy, in both sterilized and non-sterilized soil microcosms over a period of 49 days. While engineered *M. smegmatis* and Kampy persisted in all soil microcosms, there was minimal evidence of spread to 5 cm away from the inoculation site. There was a higher prevalence of Kampy observed in sterilized soil than non-sterilized soil, suggesting a detrimental effect of the native soil biotic and viral community on the ability of this phage to proliferate in the soil microcosm. Additionally, higher abundance of the genome-integrated bacteria relative to the plasmid-carrying bacteria as well as evidence for loss of plasmid over the duration of the experiment suggest a burden associated with bacteria harboring plasmids, although plasmids were still retained across 49 days. To our knowledge, this is the first study to simultaneously measure the persistence and spread of bacteria and their associated phage in both sterilized and non-sterilized soil microcosms, employing bacteria with plasmid-based and genome-integrated engineered circuits. As such, this study provides a novel understanding of challenges associated with the deployment of bioengineered microbes into soil environments.

**Importance:** Healthy soil is essential to sustain life, as it provides habitable land, enables food production, promotes biodiversity, sequesters and cycles nutrients, and filters water. Given the prevalence of soil degradation, treatment of soil with microbes that promote soil and crop health could improve global soil sustainability; furthermore, the application of bioengineering and synthetic biology to these microbes allows fine-tunable and robust control of gene-of-interest expression. These solutions require the successful deployment of bacteria into the soil, an environment in which abundant competition and often limited nutrients can result in bacterial death or dormancy. This study employs *Mycobacterium smegmatis* as a chassis alongside its bacteriophage Kampy in soil microcosms to assess the ability of non-native microbes to survive and spread in soil. Insights from this experiment highlight important challenges which must be overcome for successful deployment of engineered microbes in the field.

## Introduction

Soil health is at risk globally (1, 2) threatening biodiversity (3), nutrient cycling (4), and food production (5). Scientists have recently issued an urgent call to action for the utilization of microbes to address escalating climate change (6) due to their constantly evolving capabilities to catabolize pollutants (7, 8), sequester greenhouse gasses (9, 10), restore degraded soils and prevent erosion (11, 12), and promote crop growth (13, 14). Bioengineering enables the finely-tunable expression of these genes in organisms adapted for growth in diverse environments (15); these genetic circuits have already demonstrated potential to remediate soil environments (16, 17). However, these engineered microbes are often studied exclusively in model species grown under optimal laboratory conditions, using nutrient-rich media and with protection from adversaries—conditions that promote exponential growth and bountiful protein production (18, 19) and differ greatly from those of the soil in which the microbes must perform once deployed. In soil environments, unlike under typical laboratory conditions, bacteria can face variable temperature (20, 21) and pH (22, 23), scarce nutrients (24, 25), as well as competitors, predators, and symbionts (26–28). These harsh conditions typically promote dormancy; it is estimated that only 2% of bacteria in the soil are non-dormant (29). Additionally, engineered cells may be at a disadvantage, as genetic circuits can impose an additional burden on their chassis (30–32).

Furthermore, in natural environments, continual interactions with bacteriophage (phage) drive alterations in bacterial population dynamics, underlying the importance of examining both populations concurrently in soil. Bacterial relative fitness in communities with inoculated phage has been found to be significantly lower in natural environments, including soil microcosms, compared to high-nutrient broths (33, 34). Additionally, phage presence in soil has been correlated with changes in bacterial diversity (35) as well as nutrient contents key for healthy soils (35–38). Due to the profound effect of the soil virome on soil health and the soil microbiome, several recent reviews and opinion articles have suggested that phages can be utilized to improve soil and crop health (39–42).

While some studies have evaluated microbial survival and growth dynamics in soil, the results from these studies have been inconsistent, likely due to differences in experimental design such as bacteria type, soil conditions, and presence of inoculated phage. For example, in the absence of added phage, bacteria inoculated into sterilized soil have been shown both to increase overall in abundance (43, 44) and spread (43, 45) as well as to decrease overall in abundance (43, 44, 46). However, when bacteria is co-inoculated with phage in sterilized soil, bacteria populations tend to decrease in abundance on average (33, 47, 48). Alternatively, bacteria inoculated into non-sterilized soil have generally been shown to decrease in abundance overall, whether in the presence of co-inoculated phage (33, 48–51) or not (44, 46, 51–56). Despite this observed decline of exogenous bacterial abundance in non-sterilized soil, bacteria have still been measured spreading vertically through non-sterilized soil columns (51, 57, 58), with phage presence impeding migration (51). There is a clear need for research that systematically compares multiple key variables in order to provide a more unified perspective on microbial persistence and spread in soil environments.

To our knowledge, no single study to date has simultaneously analyzed the population dynamics of both engineered bacteria and their associated phage in soil microcosms while also comparing two major conditions which reveal essential information about the feasibility of soil synthetic biology: soil type (sterilized versus non-sterilized) and the method of genetically engineering the bacteria (plasmid-transformed versus genome-integrated). This study addresses this gap in the literature by assessing the persistence and spread of Kampy phage and its host *Mycobacterium smegmatis* — engineered to produce red fluorescence through either plasmid transformation or genome integration — when co-inoculated into soil microcosms. The findings offer critical insights into the challenges and limitations of deploying engineered bacteria for soil remediation, particularly in understanding microbial persistence, spread, and interactions with native biotic communities, and will inform future applications in sustainable agriculture and environmental restoration.

## Results

### Experimental overview

Our study tested the persistence and spread to 5 cm of *M. smegmatis* engineered with mCherry and its mycobacteriophage Kampy (“Kampy”) in soil microcosms over 49 days and across two comparisons: sterilized versus non-sterilized soil and plasmid-based (*Ms-*pC3) versus genome-integrated (*Ms-*pML(int)) *M. smegmatis* engineered strains (Fig. 1). In brief, triplicate soil microcosms were inoculated with Kampy and either *Ms*-pC3 or *Ms*-pML(int); half of the microcosms were sterilized prior to inoculation while the rest were left non-sterilized. Additional sterilized and non-sterilized soil microcosms with only PBS added were used as controls. Each was sampled at the inoculation site and at 5 cm away from the inoculation site for six time points across 49 days. Soil samples were plated for bacteria and phage to assay *M. smegmatis* and Kampy absolute abundance, respectively, and also used to identify the percentage of pink *Ms*-pC3 colonies over time to provide insight into plasmid retention.

**Fig. 1.**
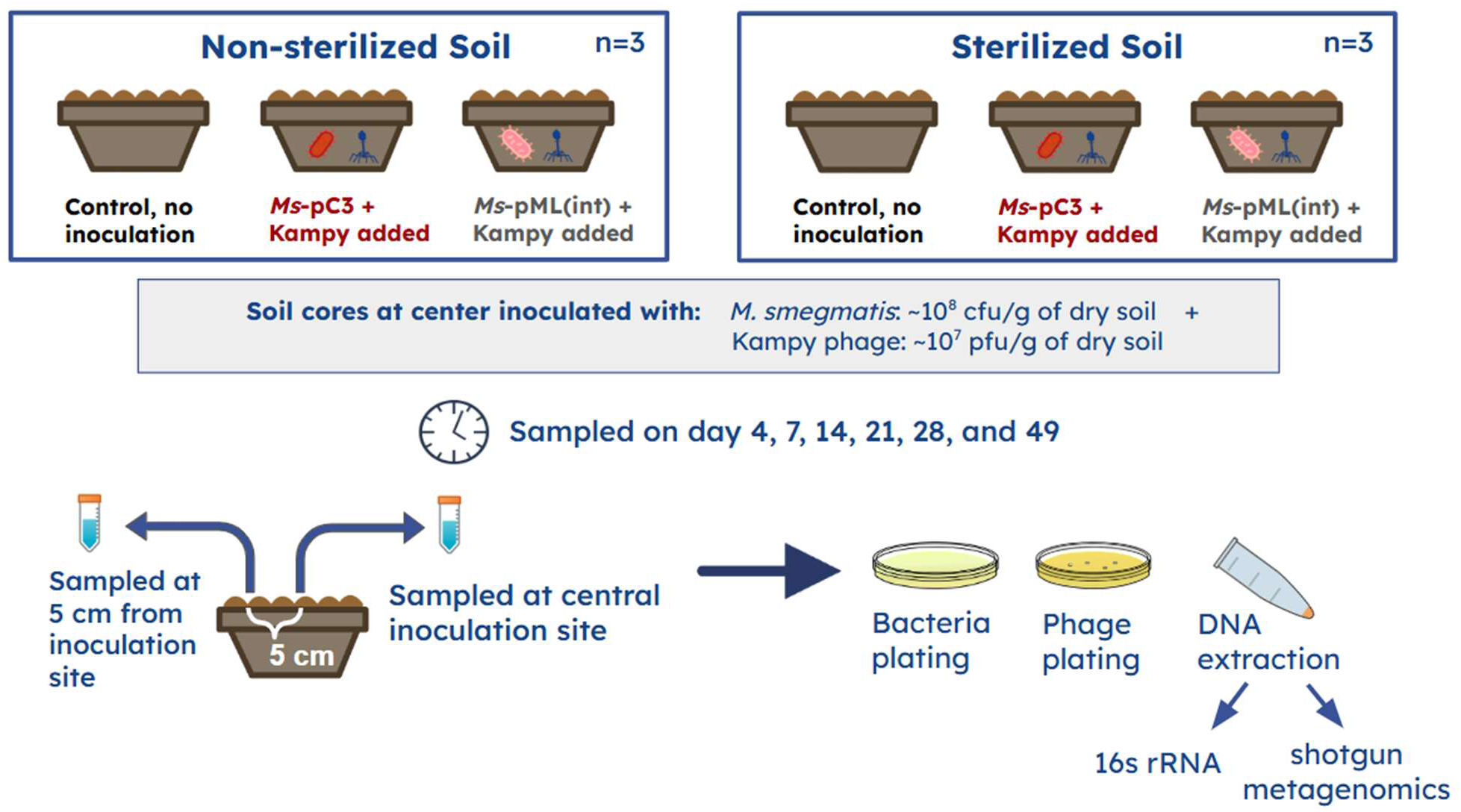
Experimental design for microcosms. A total of 18 microcosms were constructed: nine sterilized and nine non-sterilized. For each sterility type, there were three replicate control microcosms with only PBS added, 3 microcosms with *Ms-*pC3 and Kampy added, and 3 microcosms with *Ms*-pML(int) and Kampy added. Microcosms were sampled six times over a 49 day period, each time at the inoculation site and 5 cm away from the inoculation site. Each sample was plated for bacteria and phage, and had DNA extracted for 16s rRNA and shotgun metagenomic sequencing.

Additionally, DNA was extracted from soil samples for 16S rRNA and shotgun metagenomic sequencing to assay *M. smegmatis* relative abundance, as well as to perform a principal component analysis (PCA) on broader microbial community structure over time after inoculation (Fig 1). *P*-Values were computed through two-way and three-way repeated measures ANOVA, with *P* < 0.05 indicating significance and *P* < 0.10 indicating a trend.

### Engineered *M. smegmatis* persisted in both sterilized and non-sterilized soil microcosms over seven weeks

In order to assess the persistence of engineered *M. smegmatis* co-inoculated with its associated mycobacteriophage Kampy in soil over 49 days, *Ms-*pC3 and *Ms*-pML(int) absolute abundances at the inoculation site of sterilized and non-sterilized soil microcosms were measured using colony counts, while their relative abundances in non-sterilized microcosms were assayed using 16S rRNA analysis and shotgun metagenomic analysis. *M. smegmatis* abundances in these microcosms were measured at six timepoints over seven weeks and across two comparisons: soil sterilization (sterilized versus non-sterilized) and *M. smegmatis* engineering strain (plasmid-transformed versus genome-integrated).

Overall, all three assays (colony counts, 16S rRNA, and shotgun metagenomic analysis) suggest *M. smegmatis* abundance decreases over time, yet remains detectable in soil at levels significantly different from control microcosms (with no inoculation) even after 49 days. The colony counting assay showed that, across both soil types and engineering strains, *M. smegmatis* absolute abundance changed significantly over time (P = 0.007), decreasing 3.6-fold in sterilized soil and 38-fold in non-sterilized soil on average by day 49 (relative to day 4) (Fig. 2A, Table S1). This corresponded to an average final absolute abundance of approximately 10^7^ CFU/g of sterilized soil and 10^6^ CFU/g of non-sterilized soil. 16S rRNA analysis similarly showed significant changes in *M. smegmatis* relative abundance in non-sterilized microcosms over time (*P* = 0.033), with the relative abundance decreasing 1.6-fold by day 49 (relative to day 4) to 0.578% on average (Fig. 2B, Table S2). Likewise, shotgun metagenomic analysis showed an apparent decrease in the relative abundance of both *M. smegmatis* engineering strains between days 7 and 49, although this change was not statistically significant (*P* = 0.16).

**Fig. 2.**
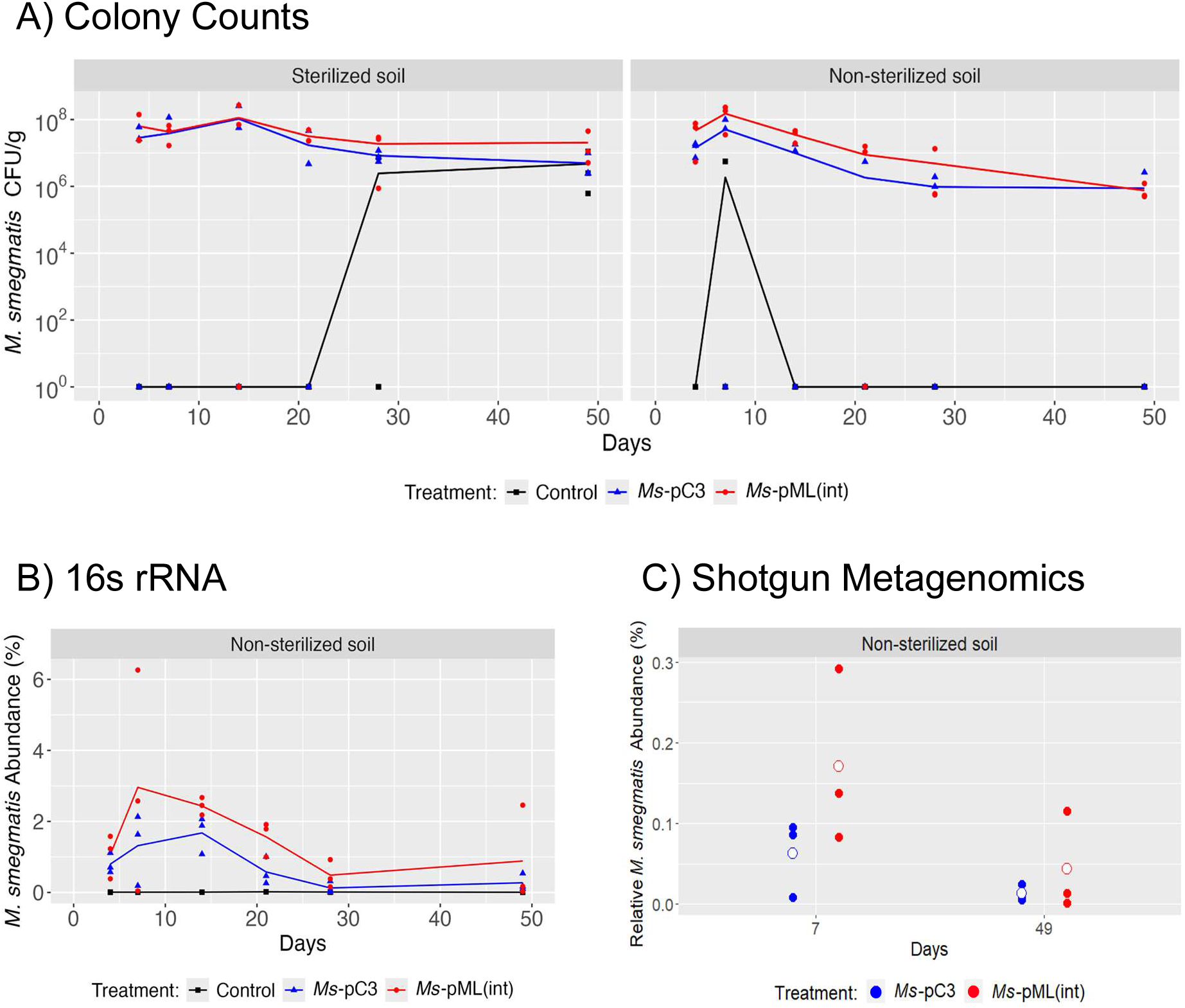
*M. smegmatis* persistence at the inoculation site over 49 days. **(A)** *M. smegmatis* CFU/g dry soil determined by colony counting in both sterilized (left) and non-sterilized (right) soil. *M. smegmatis* counts in control microcosms are shown in black, *Ms-*pC3 in blue, and *Ms-*pML(int) in red. Lines depict the average of 3 biological replicates, with each replicate displayed as an individual point. **(B)** *M. smegmatis* 16S rRNA relative abundance (as a percentage) in non-sterilized soil. *M. smegmatis* relative abundance in control microcosms are shown in black, *Ms-*pC3 in blue, and *Ms-*pML(int) in red. Lines depict the average of 3 biological replicates, with each replicate displayed as an individual point. **(C)** *M. smegmatis* shotgun metagenomic relative abundance (as a percentage) in non-sterilized soil on days 7 and 49. White circles show the mean of 3 biological replicates, with each replicate displayed as a blue (*Ms*-pC3) or red (*Ms*-pML(int)) point.

Despite the observed differences in *M. smegmatis*’s final absolute abundance between sterilized and non-sterilized soil, there was limited evidence for a statistically significant effect of soil sterilization on *M. smegmatis* abundances over time. The colony count assay revealed there were no statistically significant differences in absolute abundances at the inoculation site between sterilized versus non-sterilized soil microcosms (*P* = 0.26). However, when performing separate two-way repeated measures ANOVAs on the sterilized and non-sterilized count data, only the non-sterilized soil microcosms saw a statistically significant change in absolute abundance over time (*P* < 0.001). This significance in non-sterilized soil is in line with the aforementioned 16s rRNA results.

Additionally, the three assays suggest an effect of engineering strain on *M. smegmatis* abundance. For the colony count assay, there was a trend of colony counts differing between microcosms inoculated with *Ms*-pC3 and *Ms*-pML(int) (*P =* 0.089), with *Ms*-pML(int) absolute abundance typically being higher. Separate two-way repeated measures ANOVAs on absolute abundances measured from microcosms inoculated with either *Ms*-pC3 or *Ms*-pML(int) revealed that *Ms*-pML(int) counts had a trend of changing over time (*P =* 0.059) while *Ms*-pC3 counts did not significantly change over time *(P =* 0.27). Furthermore, when comparing both engineering strains but considering only counts from non-sterilized microcosms, there was a statistically significant difference between *Ms*-pC3 counts and *Ms-*pML(int) counts (*P* = 0.025), with *Ms*-pML(int) populations generally more abundant. We performed a permutation analysis on this data as another means of testing statistical significance: when *Ms*-pC3 absolute abundances from one replicate of the non-sterilized soil colony count data were swapped with those of a replicate from *Ms*-pML(int), eight of the nine possible permutations were not statistically significant (Table S8), consistent with the result of a difference between plasmid-transformed and genome-integrated bacterial absolute abundances. In support of colony count data, the 16S rRNA results demonstrated a significant effect of engineering strain on *M. smegmatis* relative abundance in non-sterilized soil samples (*P =* 0.043), with *Ms*-pML(int) relative abundances higher on average. Permutational analysis found that none of the nine possible permutations demonstrated statistical significance (Table S8), supporting the finding that the *Ms-*pC3 engineering strain affects the abundance of *M. smegmatis* in non-sterilized soil. Conversely, shotgun metagenomic analysis, which considered only days 7 and 49, showed no significant effect of engineering strain on *M. smegmatis* relative abundance in non-sterilized microcosms (*P* = 0.28).

### Limited evidence for horizontal spread of inoculated *M. smegmatis* to 5 cm over seven weeks in soil microcosms

Spread of engineered *M. smegmatis* to 5 cm away from the inoculation site in sterilized and non-sterilized soil microcosms over 49 days was assessed by measuring *Ms-*pC3 and *Ms-* pML(int) absolute and relative abundances using colony counts and 16S rRNA analysis, respectively. All microcosms were co-inoculated with the mycobacteriophage Kampy. Colony counts showed no significant differences in *M. smegmatis* absolute abundance at 5 cm away from the inoculation site between microcosms inoculated with *M. smegmatis* and the control microcosms with no inoculation for both sterilized (*P* = 0.12) and non-sterilized (*P* = 0.72) soil (Fig. 3A, Table S3). This indicates no apparent increase in *M. smegmatis* colony counts at 5 cm over time as a result of the spread of our inoculated bacteria. However, 16S rRNA analysis did show statistically significant differences in *M. smegmatis* relative abundance between non-sterilized treatments and controls at 5 cm from the inoculation site (*P =* 0.015), with significant changes in *M. smegmatis* relative abundance over time (*P* = 0.041) (Fig. 3B, Table S4). This suggests that inoculation of *M. smegmatis* and Kampy into soil alters the relative abundance of *M. smegmatis* at 5 cm from the inoculation site over time, a possible indicator of spread. Additionally, no significant effect of engineering strain on *M. smegmatis* relative abundance 5 cm from the inoculation site was observed (*P* = 0.54).

**Fig. 3.**
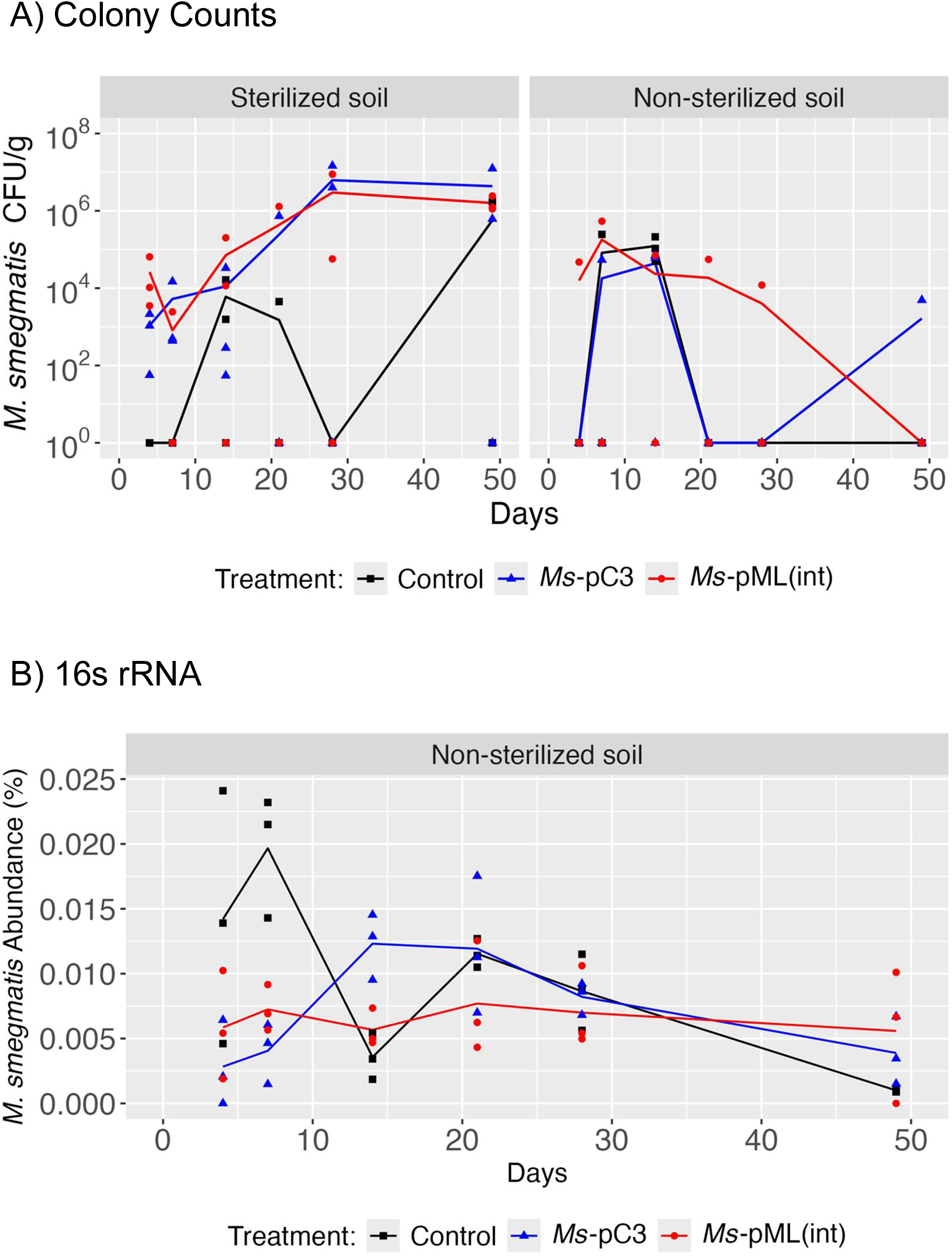
*M. smegmatis* abundance at 5 cm away from the inoculation site. **(A)** *M. smegmatis* CFU/g dry soil at 5 cm from the inoculation site as measured by colony counting. *M. smegmatis* counts in control microcosms are shown in black, *Ms-*pC3 in blue, and *Ms-*pML(int) in red. Lines depict the average of 3 biological replicates, with each replicate displayed as an individual point. **(B)** *M. smegmatis* 16S rRNA relative abundances (as a percentage) in non-sterilized soil microcosms at 5 cm from the inoculation site. *M. smegmatis* relative abundances in control microcosms are shown in black, *Ms-*pC3 in blue, and *Ms-* pML(int) in red. Lines depict the average of 3 biological replicates, with each replicate displayed as an individual point.

### Inoculated mycobacteriophage Kampy persisted in soil microcosms over seven weeks, with levels 10^5^ fold higher in sterilized soil than non-sterilized soil

In the same sterilized and non-sterilized soil microcosms inoculated with the mycobacteriophage Kampy and either *Ms-*pC3 or *Ms*-pML(int) as detailed above, the persistence of Kampy in soil was assessed over 49 days. Kampy abundances at the inoculation site were measured using plaque counts and were compared over time and across two comparisons: soil sterilization (sterilized versus non-sterilized) and *M. smegmatis* engineering strain (plasmid-transformed versus genome-integrated). Kampy, regardless of soil sterilization or whether *Ms*-pC3 or *Ms-*pML(int) was co-inoculated, remained detectable at levels statistically significantly different from control microcosms (with no inoculation) throughout the experiment.

There was a strong effect of soil sterilization on Kampy populations over time. Plaque count assays reveal statistically significant changes in Kampy absolute abundances at the inoculation site over time only in sterilized soil microcosms (*P* < 0.001), where Kampy initially increased in abundance at the inoculation site for the first 28 days before stabilizing at approximately 10^9^ PFU/g of soil, corresponding to a 11900-fold increase in abundance from day 4 to day 49. Comparatively, there was no statistically significant change in abundance over time for non-sterilized soil microcosms (*P =* 0.51) in which Kampy populations remained consistent at approximately 10^4^ PFU/g across 49 days (Fig. 4A, Table S5). Accordingly, there were statistically significant differences in Kampy absolute abundances between sterilized soil and non-sterilized soil microcosms (*P* < 0.001). Permutation analysis of Kampy absolute abundances between sterilized and non-sterilized soil microcosms indicated that none of the nine possible permutations demonstrated significance (Table S8), supporting the result that there is an effect of soil sterilization on Kampy abundance.

**Fig. 4.**
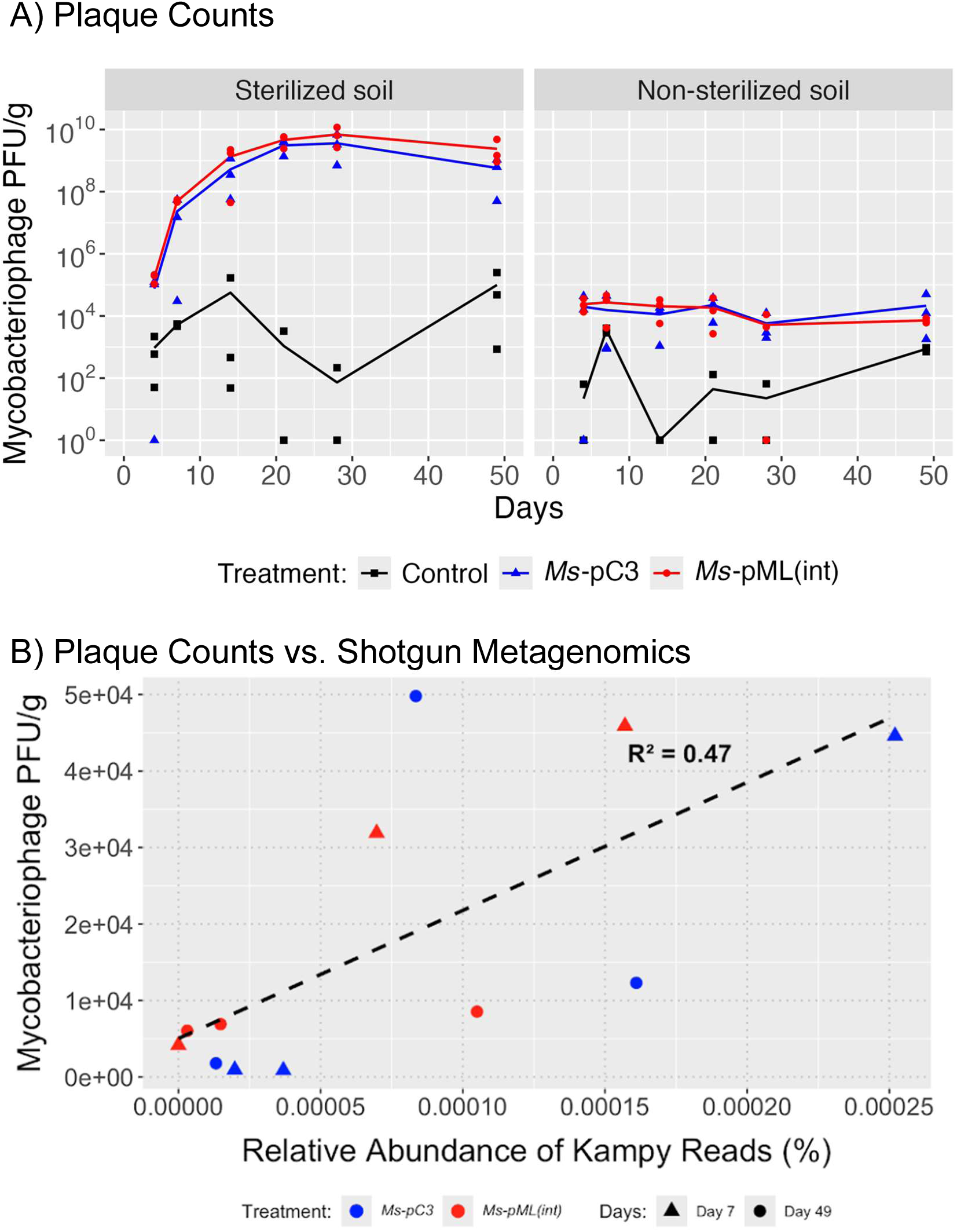
Kampy persistence at the inoculation site. **(A)** Kampy PFU/g dry soil at the inoculation site over 49 days. Kampy population density was measured by plaque counting on plates with *M. smegmatis* lawns. Kampy counts in control microcosms are shown in black, Kampy counts in *Ms-*pC3 microcosms in blue, and Kampy counts in *Ms-*pML(int) microcosms in red. Lines depict the average of 3 biological replicates, with each replicate displayed as an individual point. **(B)** The correlation between the relative abundance (as a percentage) of Kampy metagenomic shotgun sequencing reads and phage plaque counts at the inoculation site in non-sterilized soil on day 4 and 49. Soil microcosm replicates are represented by individual points, with the dotted line representing the linear correlation between the plaque assay measurement and the relative abundance of Kampy within each sample.

Additionally, there was a limited effect of *M. smegmatis* engineering strain on Kampy abundances over time. Plaque counting revealed that there was a trend of differential Kampy absolute abundance between *Ms*-pC3 microcosms and *Ms*-pML(int) microcosms (*P* = 0.052). However, when comparing absolute abundances between *M. smegmatis* engineering strains but under only one soil sterility condition at a time, this trend was only reflected in sterilized soil microcosms (*P* = 0.058), but not non-sterilized soil microcosms (*P* = 0.92).

To determine whether the plaque count assay was a precise measure of Kampy persistence, metagenomic shotgun sequencing reads derived from DNA extracted from non-sterilized microcosms on days 7 and 49 were analyzed using BLASTn against the Kampy genome. While anywhere up to 0.00025% of reads within each sample showed significant identity to Kampy ( >99% identity, >99% query length), the length and count of these reads were too ambiguous and too small to effectively quantify the presence of Kampy. This difficulty reflects ongoing challenges in the field of metagenomics, where phages are notoriously hard to detect due to understudied factors such as short read lengths, sequencing errors, assembly quality, and the complexities of phage taxonomy (59). However, PFU/g dry soil and shotgun metagenomic relative abundance of suspected Kampy reads showed a weakly positive correlation (R^2^ = 0.47), suggesting that the plaque assay correlates with suspected Kampy DNA in the soil microcosms (Fig. 4B, Table S6). A perfectly linear correlation is not expected as absolute abundance and relative abundance are not always correlated in soil environments (60).

### Inoculated mycobacteriophage Kampy spread horizontally to 5 cm over seven weeks when co-inoculated with plasmid-transformed *M. smegmatis* in sterilized soil

To examine the spread of Kampy over 49 days alongside two *M. smegmatis* engineered strains and in sterilized versus non-sterilized soil, soil was sampled from the aforementioned microcosms at 5 cm away from the inoculation site and Kampy’s absolute abundance was quantified using a direct plating plaque assay (Table S7). There was only a significant difference in Kampy abundances at 5 cm between control microcosms and inoculated microcosms in sterilized soil microcosms inoculated with *Ms*-pC3 (*P* = 0.043), but not for sterilized soil microcosms with *Ms*-pML(int) or non-sterilized microcosms with *M. smegmatis* of either engineering strain (Fig. 5). For this reason, statistical testing was performed only on counts from sterilized soil microcosms with *Ms*-pC3; this test showed a significant change over time in Kampy abundances at 5 cm from the inoculation site, suggesting that spread occurred specifically in *Ms*-pC3 sterilized microcosms (*P* = 0.025).

**Fig. 5.**
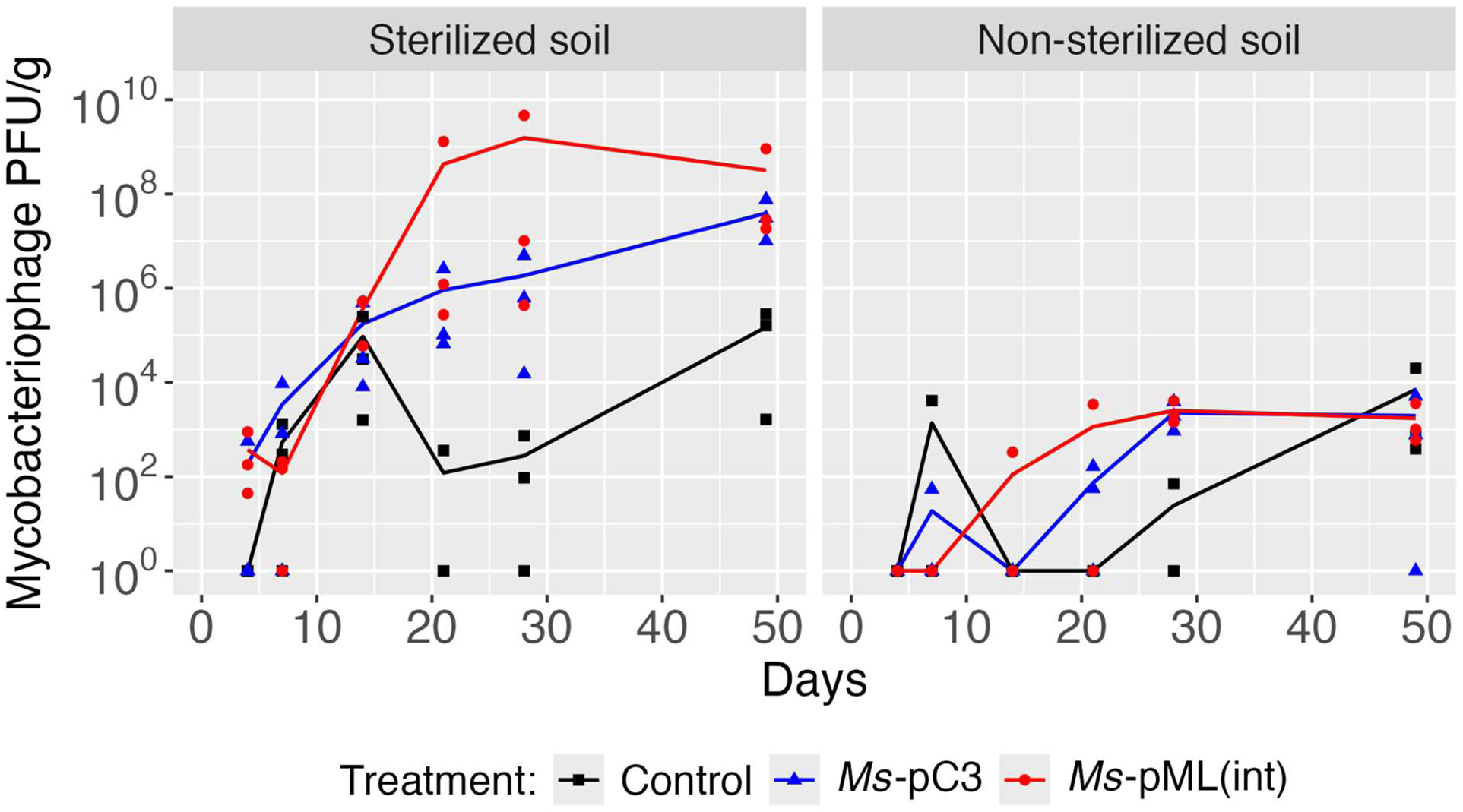
Kampy PFU/g dry soil at 5 cm away from the inoculation site. Kampy density was measured by plaque counting on plates with *M. smegmatis* lawns. Kampy counts in control microcosms are shown in black, Kampy counts in *Ms-*pC3 microcosms in blue, and Kampy counts in *Ms-*pML(int) microcosms in red. Lines depict the average of 3 biological replicates, with each replicate displayed as an individual point.

### Mycobacteriophage Kampy populations at the inoculation site experienced a greater fold-change in abundance than *M. smegmatis* populations

Patterns in bacteria and phage relationships over time were assessed by comparing the absolute abundance of *M. smegmatis* and Kampy within the same microcosms (Fig. S1). At the inoculation site, the ratio of the Kampy abundance fold-change from day 4 to day 49 relative to the *M. smegmatis* abundance fold-change from day 4 to day 49 is 43,100 in sterilized soil and 24.3 in non-sterilized soil. This corresponds to Kampy abundance increasing while *M. smegmatis* abundance decreased in sterilized soil and Kampy abundance decreasing less than *M. smegmatis* abundance decreased in non-sterilized soil, with this difference occurring to a much greater degree in sterilized soil.

### Although still detectable in both sterilized and non-sterilized soil, pCherry3 plasmid was progressively lost over seven weeks

Because *Ms-*pC3 plates showed both pink and non-pink *M. smegmatis* colony colors, the percentage of pink *M. smegmatis* colonies over time was calculated to qualitatively investigate loss of plasmid. Pink colonies were assumed to have a higher copy number of pCherry3, which was corroborated by band density PCR amplifying DNA from the pCherry3 plasmid, performed on *Ms*-pC3 colonies from sterilized soil microcosms (Fig. 6A). The mean percentage of pink colonies decreased from day 4 to day 49 in both sterilized (63% decrease) and non-sterilized (45% decrease) microcosms, suggesting the loss of plasmid over time (Fig. 6B, Table S9).

**Fig. 6.**
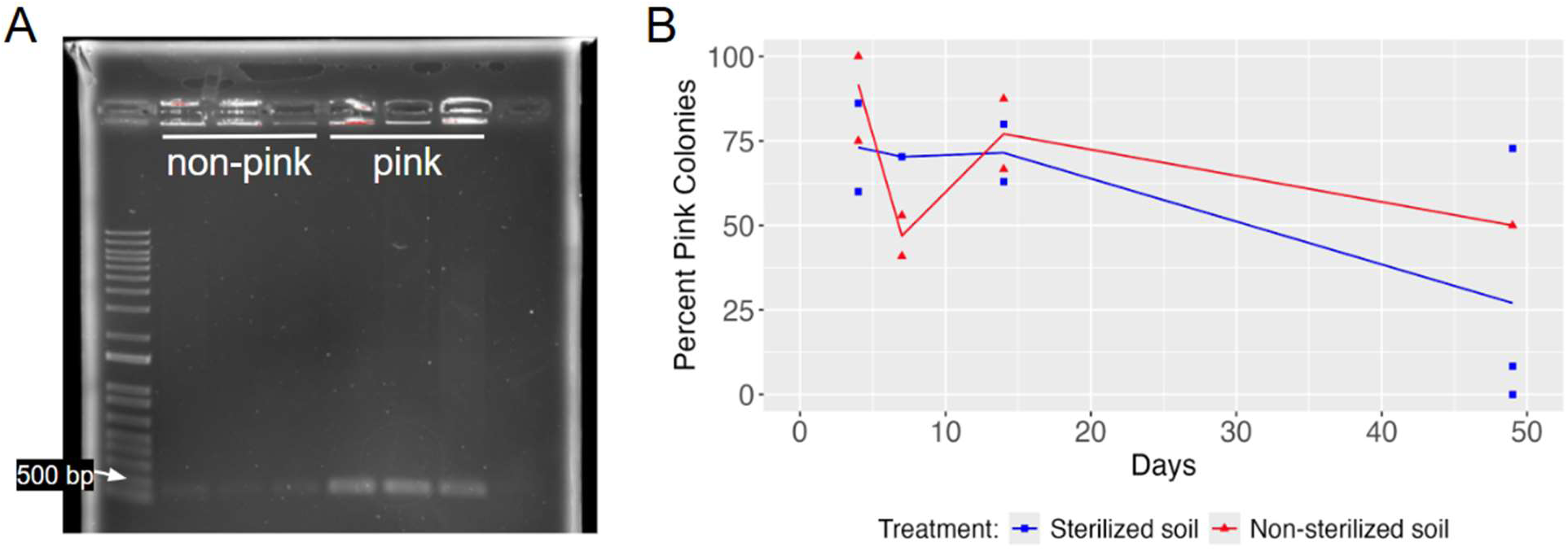
Qualitative analysis of plasmid retention in *Ms*-pC3**. (A)** Gel electrophoresis on PCR amplicons from both non-pink and pink *Ms*-pC3 colonies, with PCR targeting the pCherry3 plasmid. The last lane is a no-template control. **(B)** Percentage of pink colonies over time for sterilized and non-sterilized microcosms inoculated with *Ms*-pC3 (plasmid-transformed). The percentage of pink colonies in sterilized soil is shown in blue and the percentage of pink colonies in non-sterilized soil is shown in red. Lines depict the average of up to 3 biological replicates (replicates with no colonies were not considered), with each replicate displayed as an individual point.

### Inoculation of exogenous *M. smegmatis* and mycobacteriophage Kampy affected soil community structure

To determine if there was an effect of inoculating engineered *M. smegmatis* and Kampy on soil microbial community composition in non-sterilized soil microcosms, principal component analyses (PCAs) were performed on 16S rRNA relative abundances (all time points) and shotgun metagenomic relative abundances (days 7 and 49). PCAs on both assays revealed grouping of samples by time. For the 16S rRNA relative abundance PCA, the first time point (day 4) and the last time point (day 49) clustered separately from the rest of the data (Fig. 7A). Moreover, day 4 samples of microcosms inoculated with *M. smegmatis* and Kampy clustered separately from control microcosms with only PBS added, suggesting inoculation creates an initial disruption to bacterial community structure within the first few days. Similarly, the shotgun metagenomic PCA showed grouping of samples by time, differentiating day 7 and day 49 (Fig. 7B). This further suggests an effect of *M. smegmatis* and Kampy inoculation on the relative abundances of species members within the broader microbial community.

**Fig. 7.**
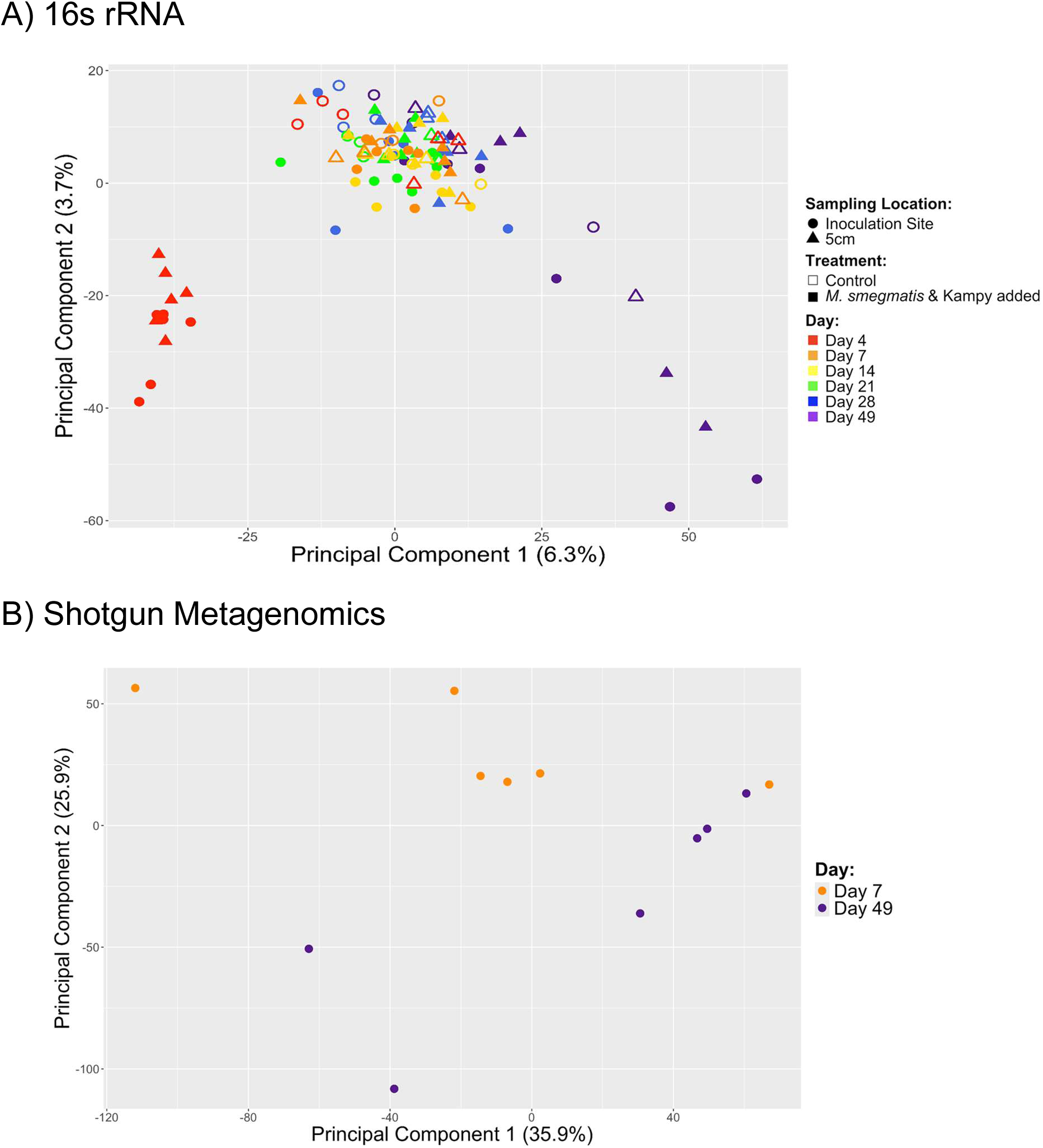
Microbial community structure after inoculation of engineered *M. smegmatis* and Kampy phage. Principal Component Analysis performed in R. **(A)** PCA on 16S rRNA sequencing reads from non-sterilized soil microcosms across all timepoints. **(B)** PCA on shotgun metagenomic sequencing reads from the inoculation site for non-sterilized soil microcosms on days 7 and 49.

## Discussion

### Overview

This study evaluates the persistence and spread of genetically engineered *M. smegmatis* and its mycobacteriophage Kampy in sterilized and non-sterilized soil microcosms, providing crucial insights into the challenges of using exogenous microbes to address environmental issues within soil ecosystems. Four major findings emerge from our study. First, inoculated *M. smegmatis* and Kampy phage persist in soil microcosms over seven weeks regardless of the method used to engineer *M. smegmatis* (plasmid transformation or genome integration) or whether the soil was sterilized. Second, Kampy population densities were several orders of magnitude lower in non-sterilized soil compared to sterilized soil, suggesting that non-sterilized soil conditions may limit the persistence of this phage. Third, while the higher abundance of *Ms*-pML(int) than *Ms*-pC3 in non-sterilized soil and the plasmid loss observed throughout the experiment are indicative of plasmid burden, some plasmids were nonetheless retained over the seven week period. Fourth, minimal observation of the spread of *M. smegmatis* or Kampy populations to 5 cm away from the inoculation site over 49 days indicates potential challenges for future deployment of exogenous bacteria into soil environments.

### Engineered *M. smegmatis* and mycobacteriophage Kampy persisted in microcosms regardless of soil sterilization or bacterial engineering method

If exogenous bacteria are to be deployed into the field, persistence of inoculated constructs is essential. While *M. smegmatis* remained detectable on average for all assays across all inoculated microcosms over the 49 day experiment, it decreased significantly in abundance in non-sterilized soil over time according to colony counting and 16S rRNA analysis – although this was not supported by shotgun metagenomic analysis. This declining prevalence of bacteria over several weeks when co-inoculated with phage into non-sterilized soil is consistent with other reports in the literature (33, 48–51). While the decreased abundance of *M. smegmatis* over time in non-sterilized soil microcosms of the present study could suggest a continuing decline beyond the duration of the experiment, its survival over 49 days is nonetheless promising for in-field applications.

While statistical tests on our colony count data indicate that *M. smegmatis* population densities at the inoculation site are not significantly altered by soil sterilization, important differences in the behavior of *M. smegmatis* between sterilized and non-sterilized soil were still observed. Notably, *M. smegmatis* abundance only changed significantly over time in non-sterilized soil, and the effect of engineering method on *M. smegmatis* abundance was only significant in non-sterilized soil. Previous soil microcosm studies have shown significant differences in inoculated bacterial abundance between sterilized and non-sterilized soil; for example, *P. fluorescens* populations were significantly greater in sterilized than non-sterilized soil, by a factor of about 4, when co-inoculated with its lytic phage SBW25ø2 after 20 days (33). Additionally, *E. coli* population densities have been found to be approximately 1000 times greater in sterilized than non-sterilized soil 21 days after the temperate transducing P1 phage and its *E. coli* host were co-inoculated into soil microcosms (48). These results suggest that there are species-specific differences in the persistence of bacteria inoculated into soil and affirm that soil sterilization does impact bacterial colonization patterns.

The microbial community present in our non-sterilized soil samples was assessed through principal component analysis (PCA) of 16S rRNA and shotgun metagenomic relative abundance data. Grouping of samples by day observed in both PCA plots suggests that microbial community compositions change following inoculation with *M. smegmatis* and Kampy, remain steady for the next several weeks, and then change again after a longer period of incubation. These results support previous literature which has characterized an effect of bacterial inoculation on the clustering of bacterial reads from soil microcosms (61, 62). The impact of inoculated *M. smegmatis* and Kampy on the broader biotic community emphasizes the importance of microcosm studies to ensure future field-deployment does not adversely affect ecosystems.

### Mycobacteriophage Kampy persisted at significantly lower abundance in non-sterilized soil than sterilized soil

Understanding the persistence of phage inoculated into soil is imperative both in consideration of phage as a predator for added engineered bacteria or even as a possible vector for in-situ editing in soil (63). As measured by plaque counts, the absolute abundance of Kampy on day 4 at the site of inoculation was less than an order of magnitude greater in sterilized than non-sterilized soil, but by day 49 was about five orders of magnitude greater in sterilized soil. This corresponds to a statistically significant change in plaque counts over time in sterilized soil, where Kampy abundance increased on average over time, but not in non-sterilized soil microcosms. The significant difference in Kampy counts between sterilized and non-sterilized microcosms may be explained by the abiotic and biotic aspects of non-sterilized soil dampening Kampy phage populations. Concentrations of nitrate, ammonia, phosphate, potassium, and organic matter have all been shown to change by less than 3-fold after 4 hours of autoclaving soil at 150°C (64), while concentrations of dissolved organic carbon (DOC) can increase 20–37 fold (65, 66). Past studies have demonstrated positive correlations between viral abundance and organic carbon, nitrate, potassium, and calcium (36, 67), but the effect of DOC on phage abundance still appears to be unknown. pH changes from autoclaving soil are unlikely to explain our results since autoclaving soil typically induces only small decreases in soil pH (64–66, 68–70).

Autoclaving soil also significantly reduces the bacterial and fungal species richness and diversity (71); these changes may be beneficial to phage persistence. Many soil bacteria enter a dormant state due to carbon limitation resulting from the intense competition of natural soil environments (25, 55). Mimicking this state by limiting carbon in *E. coli* growth media has been shown to halt replication of T4, λ, and P1 phages; this could similarly explain the low Kampy populations in our non-sterilized soil microcosms as a result of bacterial competition and dormancy (72, 73). Furthermore, native bacteria can release outer membrane vesicles that capture free phage virions, thereby blocking infection (74). In alignment with our results, phage populations have been observed to be substantially more abundant in sterilized than non-sterilized soil, with phage densities ranging from 10^4^–10^6^ PFU/g in sterilized soil versus nearly zero in non-sterilized soil (48). A later study reaffirmed this trend (33), but with much smaller magnitudes of difference likely due to variation in bacterial and phage species used and soil type. In particular, inoculation of either temperate or lytic phage can result in significantly different population abundances in soil; measurement of phage population density in sterilized soil microcosms inoculated with *B. subtilis* and temperate phage THø versus lytic phage SP10C demonstrated that the lytic phage population density leveled out at about 100 times greater than that of the temperate phage over 40 days (47). These results may be attributable to the higher replication rate of lytic phage or may suggest the presence of a hidden population of lysogenic phages integrated within their hosts and thereby undercounted by standard plate counts. While this study extended these conclusions by investigating the behavior of temperate phage in non-sterilized soil microcosms, further investigation into the role of lysogeny or lytic phages in non-sterilized soil systems is necessary.

### The mCherry plasmid is detrimental to the survival of *M. smegmatis* in non-sterilized soil

A particularly novel feature of this study examines the effect of the method utilized to engineer *M. smegmatis* (plasmid transformation versus genome integration) on the persistence and spread of *M. smegmatis* and its associated mycobacteriophage Kampy co-inoculated into soil microcosms. *M. smegmatis* population densities at the inoculation site, as determined by colony counts, non-significantly trend higher in microcosms inoculated with *M. smegmatis* modified with genome integration (*Ms*-pML(int)) than in those with plasmid transformation (*Ms*-pC3). In non-sterilized soil specifically, *M. smegmatis* population densities were significantly different in microcosms inoculated with *Ms*-pML(int) than with *Ms*-pC3; this amounts to approximately a 3-fold greater abundance for Ms-pML(int) on average. Furthermore, the decline in the percentage of pink colonies from microcosms inoculated with *Ms*-pC3 indicates that over 49 days, on average, only about 27% of CFUs in sterilized soil and 50% of CFUs in non-sterilized soil retained the mCherry plasmid. 16S rRNA analysis also showed significantly different *M. smegmatis* relative abundances in non-sterilized microcosms inoculated with *Ms*-pML(int) compared to *Ms*-pC3, with Ms-pML(int) generally more abundant. Although this statistically significant difference in Ms-pC3 and Ms-pML(int) relative abundances was not supported by shotgun metagenomic analysis, the evidence from 16S rRNA, colony count, and colony color assays indicate a small but measurable survival advantage of the genome-integrated construct in non-sterilized soil. This affirms concerns about the negative impact of plasmid burden on host growth (30–32). However, the continued detection of pink colonies, the presence of *Ms*-pC3 colonies at approximately 10^6^ CFU/g dry soil, and the non-zero relative abundances for all 16S rRNA samples even after 49 days suggest plasmid-transformed constructs would still serve as an effective tool for deployment into non-sterilized soil.

### Limited horizontal spread of *M. smegmatis* and mycobacteriophage Kampy under dry soil conditions

Successful deployment of engineered microbes in the field also requires spread of these microbes through the inoculated environment. While previous literature has demonstrated bacterial spread over time both horizontally over smaller distances and vertically (43, 45, 51, 57, 58), many of these experiments included regular watering (45, 51, 57, 58) and motile bacteria (43, 45, 51, 57, 58). Our experiment, which deployed a non-motile bacteria and watered microcosms only twice over seven weeks, did not provide determinative support for statistically significant spread of *M. smegmatis* in either sterilized or non-sterilized soil to 5 cm from the inoculation site over 49 days. Specifically, *M. smegmatis* population densities at 5 cm from the inoculation site were only significantly different than the abundances from control microcosms with no inoculation as assayed by 16S rRNA sequencing but not as measured by colony counts. In the plaque counting assay, spread was only suggested for phage in sterilized soil microcosms inoculated with *Ms*-pC3, indicating that spread to 5 cm over 49 days may be possible for phage under certain conditions but not all. Going forward, while research has shown an inverse relationship between soil microbial diversity and the survival of invading bacteria likely due to competition over resources (75), this must be extended to examining the impact of community structure on bacterial spread.

Furthermore, microcosm experiments cannot fully replicate the complexity of natural soil environments, including effectors of spread such as rainfall, the churning of soil by animals and plant roots, and nonhomogeneity across soil types. Moreover, as autoclaving does not fully sterilize the soil (71, 76), abiotic conditions cannot be truly simulated. The limited research on bacterial spread, particularly investigating bacteria and phage in non-sterilized soil, highlights a critical gap in understanding the real-world applicability of exogenous microbes in soil environments.

### Conclusions

This microcosm study was the first of its kind, to our knowledge, to measure the persistence and spread of co-inoculated phage and bacteria, engineered with plasmid transformation or genome integration, in sterilized and non-sterilized soil. The lack of conclusive spread observed for *M. smegmatis* and Kampy and the significantly lower Kampy abundance in non-sterilized soil compared to sterilized soil highlights a critical limitation for bioremediation applications requiring field deployment of bacteria or phages. However, the persistence of *M. smegmatis* and Kampy in both sterilized and non-sterilized soil over 49 days poses a promising avenue for future synthetic biology environmental applications. The minor differences in persistence between plasmid-transformed and genome-integrated *M. smegmatis* constructs indicate that, while it has some effect, plasmid burden may not be the most significant determinant of bacterial survival in soil conditions; however, loss of plasmid does occur without selection. Moving forward, this work underscores the necessity of developing approaches that enhance both persistence and dispersal of microbes in non-sterilized soils, providing a clear direction for future research aimed at optimizing microbial deployment for environmental applications.

## Materials and Methods

### M. smegmatis engineering

*M. smegmatis*, a non-pathogenic, fast-growing, gram-positive microbe, was chosen for this experiment because of its ubiquity across soil ecosystems and potential for bioengineering applications (77). Two different engineered strains of *M. smegmatis* mc^2^155 (ATCC #700084) were used in the microcosm experiments: one with mCherry, a red fluorescence protein gene, transformed in a plasmid (called *Ms*-pC3) and one with mCherry integrated into the genome (called *Ms-*pML(int)). In brief, electrocompetent *M. smegmatis* cells were prepared by chilling cells (OD600 0.8–1.0) on ice, centrifuging (ThermoScientific Sorvall Legend Micro 17R Centrifuge) at approximately 2500 × *g* for 10 minutes, washing cells three times in 10% glycerol with centrifugation after each wash, and then resuspending cells in 10% glycerol and storing at - 80°C until use (78, 79).

To engineer *Ms-*pC3, as adapted from several protocols (80, 81, Eppendorf Protocol No. 4308 915.522), 1 µg of pCherry3 (82) (AddGene #24659) DNA was added to 60 µl of electrocompetent *M. smegmatis*, mixed by pipetting, and then incubated on ice for 10 minutes. This mixture was transferred to a 1 mm gap electroporation cuvette and inserted into an Eppendorf Eporator. One pulse (5.0 msec time constant, 12 kV/cm field strength) was delivered. Electroporated cells were incubated on ice in the cuvette for 5 minutes, and then resuspended in 1 ml of Middlebrook 7H9 Media with 10% AD Supplement and transferred to a microcentrifuge tube. Cells were then incubated at 37°C with constant shaking (250 rpm) for 2 hours before being plated (∼100 µl per plate) on 7H9 agar plates with hygromycin (200 µg/ml) for selection. Plates were incubated at 37°C for 48 hours. Single colonies were selected and incubated in 7H9 media with hygromycin (200 µg/ml) for 72 hours at 37°C and 250 rpm. Fluorescence of the construct was confirmed on a BioTek Synergy H1 Microplate Reader (600 nm absorbance, 488 nm excitation wavelength, 530 nm emission wavelength).

To engineer *Ms-*pML(int), the construct ‘pMLcherry’ was created using pCherry3 (82) (AddGene #24659) and the integration vector pML1357 (83) (AddGene #32378). To replace the green fluorescence protein gene in pML1357 with mCherry, a restriction digest was performed on both pML1357 and pCherry3 using XbaI (cut site: TCTAGA) and HindIII (cut site: AAGCTT). The mCherry insert of pCherry3 and the backbone of pML1357 were gel extracted and purified using a Qiagen MinElute Gel Extraction Kit (#28604) following the manufacturer’s protocol and ligated overnight using a T4 ligase. The resulting 6408 bp ‘pMLcherry’ plasmid (Fig. S2) was transformed, as adapted from several protocols (84, 85) into *Escherichia coli* DH5-alpha by mixing ∼100 ng of DNA with 50 µl of competent cells (Invitrogen DH5a #18258012) and incubating on ice for 30 minutes. Then, the microcentrifuge tube containing the cells and DNA was held in 42°C water for 45 seconds and then incubated on ice for 2 minutes. Cells were afterwards suspended in 950 µl SOC Medium (NEB #B9020S) and incubated at 37°C with constant shaking (250 rpm) for 1 hour. Cells (∼100 µl per plate) were then plated on LB agar with hygromycin (200 µg/ml) and incubated overnight at 37°C. The plasmid was isolated with a New England BioLabs Monarch Plasmid MiniPrep Kit (#T1010S) according to the manufacturer and electroporated into *M. smegmatis* as previously described. This *M. smegmatis* strain was grown for approximately 72 hours at 37°C on 7H9 agar plates with hygromycin (200 µg/ml). Fluorescence of this strain was confirmed via plate reader as previously described. Integration of the plasmid into the genome was confirmed by diagnostic PCR performed on a Thermocycler with NEB Q5 High-Fidelity Master Mix at an annealing temperature of 55°C with one primer (5’ - GGACGAGCTGTACAAGTGAAA) targeting pMLcherry and the other (5’ - TTCTTCTCGACACCGTTGAAG) targeting the *M. smegmatis* genome near the attB site. Confirmed colonies were then inoculated into 7H9 media without antibiotics to encourage the loss of unintegrated plasmid. Accordingly, *Ms*-pML(int) media displayed 3–5-fold lower red fluorescence levels compared to *Ms*-pC3 media (results not shown).

### Soil collection and characterization

Soil was collected from a home garden in Williamsburg, Virginia (Latitude: 37.283115 / N 37° 16’ 59.215’’, Longitude: -76.664902 / W 76° 39’ 53.646’’) and homogenized by passage through a 1 mm sieve. A portion of the soil was sterilized by autoclaving on a G60 cycle three times and then baking at 150°C in an oven for 24 hours. The dry weight of the soil was calculated by baking 60 g wet weight of both the sterilized and non-sterilized soil for 16 hours in an oven at 150°C. The soil moisture content was calculated using the difference in weight before and after baking divided by the dry weight.

The composition of the sterilized and non-sterilized soil was assessed (Table 1). In brief, organic carbon (as a percentage of weight) was measured by weight loss on ignition. Total carbon and nitrogen (as a percentage of weight) were measured using a Perkin-Elmer 2400 Elemental Analyzer. Total phosphorus (as a percentage of weight) was measured using an ashing/acid hydrolysis method (86).

**Table 1.** Organic material, carbon, nitrogen, and phosphorus concentrations (as a percentage of weight), as well as soil moisture content, of sterilized and non-sterilized soil.

|  | % Organic material | % Carbon | % Nitrogen | % Phosphorous | Dry basis moisture content (g water/g dry soil) |
| --- | --- | --- | --- | --- | --- |
| Non-sterilized Soil | 5.95 | 3.00 | 0.23 | 0.03 | 0.417 |
| Sterilized Soil | 6.05 | 3.16 | 0.24 | 0.04 | 0.286 |

### Preparation and sampling of soil microcosms

Microcosms were constructed in 1.9 L plastic containers, approximately 20 x 20 x 7 cm in dimensions with 16 evenly spaced drainage holes drilled into the bottom. There were a total of 18 microcosms, each containing approximately 500 g (dry weight) of soil. To bring the microcosms to approximately similar moisture contents, 25 ml of autoclaved (L60) water was added to each non-sterilized microcosm and 130 ml was added to each sterilized microcosm.

Sterilized and non-sterilized soil microcosms were inoculated either with *Ms-*pC3 and Kampy or with *Ms-*pML(int) and Kampy (n = 3) (Fig. 7). There were additionally three sterilized and three non-sterilized control microcosms with no inoculation. Kampy, an A4 temperate mycobacteriophage, was selected due to existing transcriptomic characterization and its soil origins (87).

To add the *M. smegmatis* and Kampy, 13 g cores of soil were taken from the center of each microcosm. *M. smegmatis* was pelleted (4100 × *g* for 5 min), suspended in 1 ml of Phosphate Buffered Saline (PBS) to a concentration of approximately 10^8^ CFU/g of dry soil, combined with approximately 10^7^ PFU/g of dry soil of Kampy Phage, and added to the 13 g soil core. This soil core was thoroughly mixed and then placed back into the center of the microcosms. For control microcosms, 1 mL of uninoculated PBS was added to the soil core, thoroughly mixed, and placed back into the center.

Sterilized microcosms were kept in a sterile hood throughout the duration of the experiment while non-sterilized microcosms were kept on a benchtop, both receiving ambient room light. All microcosms were covered loosely with a plastic lid. Microcosms were watered on days 6 and 42 by sprinkling 15 and 50 ml, respectively, of autoclaved (L60) water over the top of each microcosm, resulting in relatively dry soil conditions.

Soil was sampled with a sterilized scoopula at the center of each microcosm (the inoculation site) and approximately 5 cm away from the inoculation site on days 4, 7, 14, 21, 28, and 49. Approximately 1 g of soil was removed each time soil was sampled. DNA was extracted from 0.25 g of this soil using a Qiagen DNeasy PowerSoil Pro Kit (#47014), according to the manufacturer’s instructions, and eluted into 50 μl of Elution Buffer. Approximately 0.5 g of the remaining sampled soil was used for standard bacterial plate counting and direct phage plaque assays. In brief, the ∼0.5 g soil sample was suspended in 2 ml of phage buffer (10 mM Tris [pH 7.5], 10 mM MgSO4, 68 mM NaCl, 1 mM CaCl2 dissolved in ddH2O), vortexed, and diluted in PBS as necessary. A relatively high dilution of the soil suspension (10^-5^–10^-4^) was used for bacterial plating of samples taken from the inoculation site to prevent non-*M. smegmatis* colonies from obscuring *M. smegmatis* colonies on the plate. For bacterial counting, 100 µl of the diluted suspension was spread on 7H9 agar plates. Then, the remaining solution was centrifuged (2000 × *g* for 10 minutes) and the supernatant was filtered through a 0.22 µm filter and diluted in phage buffer as necessary. For direct plaque counting, 100 µl of this filtered sample was added to 1 ml of untransformed *M. smegmatis* and 900 µl of phage buffer, incubated for 20 minutes, then plated on LB bottom agar plates with 2X 7H9 top agar (without antibiotics). Bacterial plates were typically counted after 72 hours and phage plates were typically counted after 48 hours. For bacterial plates, only colonies with morphology similar to *M. smegmatis* (identified as small, yellow-tinted, and wrinkled) were counted (88). Plates with no colonies or plaques were recorded as having an abundance of zero. Phage plates for which webbed lysis occurred were recorded as having 5000 plaques and plates that were completely cleared were recorded as having 10000 plaques. CFU and PFU counts were normalized to the dry weight of the soil sample (Fig. 8).

**Fig. 8.**
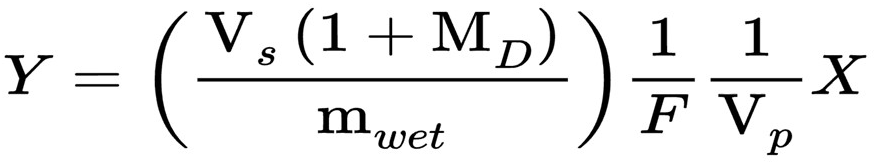
Normalization of CFU or PFU/g dry soil. Y = CFU or PFU/g dry soil, Vs = volume of liquid for soil suspension (2 ml in all cases here), MD = dry basis moisture content of soil (Table 1), mwet = mass of wet soil sampled, F = dilution factor (negative exponent), Vp = volume of dilution plated, X = CFU or PFU counted on a given plate.

### PCR confirmation of colony and plaque identities

To confirm the identities of colonies and plaques being counted, random samples were selected from plates, underwent DNA amplification using PCR, and visualized via agarose gel electrophoresis. A 30-cycle PCR was performed on a Thermocycler at 55°C annealing for colony PCR and 67°C annealing for plaque PCR, with 98°C denaturation and 72°C extension steps for both. For colony PCR, 12 distinct colonies counted as *M. smegmatis* from 4 unique plates were selected using sterilized toothpicks. PCR was performed as previously described with 5 μl of NEB Q5 Hot-Start Hi-Fi Master Mix, 3 μl nuclease-free water, 0.5 μl each of *M. smegmatis* forward (5’ - ATGGAACCTGTCGACGGGGC) and reverse (5’ - ACTGCTCGTCGCGTGTCTGG) primers, and 1 μl of colony suspension. Products were run on a 1% agarose gel. For plaque PCR, 5 plugs from 5 unique plates, including one from a microcosm without *M. smegmatis* or Kampy added to serve as a negative control, were selected. Plaque plugs were heated to 85°C for 15 minutes to promote phage DNA solvation, then PCR was performed as previously described with 5 μl of NEB Q5 Hot-Start Hi-Fi Master Mix, 3 μl nuclease-free water, 0.5 μl each of Kampy forward (5’-CGTTCCTCACACCCATTCC) and reverse (5’-ACGCTGTGATCGAGACTGAC) primers, and 1 μl of plug suspension. Products were run on a 1% agarose gel with two Kampy positive controls. Gel electrophoresis with both *M. smegmatis* and Kampy PCR products showed amplicons of the expected length for every colony and plaque tested, suggesting that counted colonies and plaques were accurately identified as *M. smegmatis* and Kampy, respectively.

### Color analysis of plasmid retention in *Ms*-pC3

To qualitatively investigate loss of plasmid in *Ms-*pC3, the amount of both pink and non-pink *M. smegmatis* colonies were counted for all *Ms*-pC3 plates sampled from the inoculation site on days 4, 7, 14, and 49 by two independent researchers. These counts were used to calculate the percentage of pink colonies on each plate. On plates with no colonies, no percentage was calculated. The average percentage of pink colonies for each group of at-most three replicates calculated between the two researchers were always within 15% of each other, and these two values were averaged when there was disagreement.

To assess whether this colorimetric analysis was an appropriate measure of plasmid retention, PCR targeting plasmid DNA was performed on 3 non-pink and 3 pink colonies from sterilized soil microcosms on Day 28 as previously described with 5 μl of NEB Q5 Hot-Start Hi-Fi Master Mix, 3 μl nuclease-free water, 0.5 μl each of *M. smegmatis* forward (5’ - CTACAGCGTGAGCTATGAGAAA) and reverse (5’ - CGAAACCCGACAGGACTATAAA) primers, and 1 μl of colony suspension. Products were run on a 1% agarose gel to confirm that pink colonies resulted in a visually denser band than non-pink colonies.

### Bacterial 16S rRNA sequencing

Using DNA extractions from microcosm samples detailed above, 16S rRNA sequencing was performed by Illumina on all non-sterilized soil extractions. In brief, Omega Bioservices amplified 16S rRNA gene V3-V4 regions using a forward primer (5’-TCGTCGGCAGCGTCAGATGTGTATAAGAGACAGCCTACGGGNGGCWGCAG) and reverse primer (5’-GTCTCGTGGGCTCGGAGATGTGTATAAGAGACAGGACTACHVGGGTATCTAATCC). A 25-cycle PCR reaction using KAPA HiFi HotStart ReadyMix (Kapa Biosystems, Wilmington, MA) was performed with initial denaturation at 95°C, 55°C annealing, 72°C extension, and 72°C elongation steps. Mag-Bind RxnPure Plus magnetic beads (Omega Bio-tex, Norcross, GA) were used to clean up the PCR product. Another index PCR amplification and elongation step was then performed. Agilent 2200 TapeStation and QuantiFluor dsDNA System (Promega, Madison, WI) were used, respectively, to check and quantify the ∼600 bp library. Nextseq (Illumina, San Diego, CA) then normalized, pooled, and sequenced the library on a 2x 300 bp end-read setting.

To independently calculate *M. smegmatis*’s relative abundance in the microcosms, query sequences were created using the first and last 300 bp of *M. smegmatis*’s V3-V4 16S region (NCBI genome #CP000480.1). Both query sequences were reduced to the internal 280 bp to account for discrepancies at the edges of the 16S amplicons caused by PCR. The forward read files were queried with their respective *M. smegmatis* 16S region query sequence, and only 100% identity hits were used. The number of hits, relative to the total number of reads provided by Illumina in that sample, was considered the relative abundance of *M. smegmatis* in that microcosm (Table S2, Table S4).

### Shotgun metagenomic sequencing

DNA extractions from non-sterilized microcosm samples on days 7 and 49 were sent for shotgun metagenomic sequencing by the Oklahoma Medical Research Foundation NGS Core using a NovaSeq X Plus (Illumina, San Diego, CA) with paired-end settings of 2 x 150 bp.

Whole DNA genome libraries were prepared by the provider using the IDT xGen DNA EZ prep kit and with Kapa qPCR and Agilent TapeStation for quality control. Twelve paired-end shotgun metagenomic sequencing samples were first assessed for quality and adapter content utilizing FastQC v0.12.1 (89). Adapters and low-quality base pairs were trimmed with default parameters utilizing trim_galore v.0.6.10 and evaluated with FASTQC once more to ensure adapters were removed. Once the raw sequences were processed for quality, two main classifications of specific microbial presence were utilized.

First, taxonomic classification of trimmed reads was performed utilizing Kraken2 v.2.1.3 (90) across RefSeq Bacterial (91) and Viral (92) libraries using default parameters. Confidence score thresholds of 0.2 and 0.5 were tested across all twelve samples, yielding a lower classification rate ranging from 3.63–11.43% and 0.85–8.61% reads classified respectively. A default confidence score of 0.0 yielded the highest percentage of reads classified, ranging from 22.45–29.04%. Kraken2 classification results were then run through Bracken v2.9 with default parameters to estimate relative abundance.

Second, suspected Kampy phage reads were identified by first converting trimmed reads from FASTQ to FASTA format utilizing seqtk v.1.4-r122. A reference database was then constructed from Kampy’s genome derived from PhagesDB (93) utilizing BLAST v2.15.0+ (94). Each FASTA sequencing read file was then run using the BLASTn command utilizing the new Kampy reference database (Table S6). All software were installed and managed on Anaconda3 conda environments and run using the Hima and Gust subclusters of the William & Mary High-Performance Computing cluster, which offered up to 128 cores per node for run parallelization and optimized computing time.

### Statistical analysis and data visualization

All statistical analyses were performed in R v4.4.1 (95). The count data was analyzed with three-way repeated measures analysis of variance (ANOVA) tests, with differences among *M. smegmatis* and Kampy counts assessed on the sterility of microcosms, the engineered bacterial strain, and time. We additionally performed two-way repeated measures ANOVAs on *M. smegmatis* and Kampy counts to further investigate a specific sterility or engineered strain treatment, when applicable. Two-way repeated measures ANOVA tests were employed for 16S rRNA forward sequence relative abundances to compare differences across engineered bacterial strain and time. Two-way repeated measures ANOVA tests were also performed for shotgun metagenomic relative abundances found using Bracken v2.9 default settings to analyze engineered bacterial strain and time differences. Significance was set for all statistical tests at *P* < 0.05. As an additional test, permutational analysis was performed on all ANOVA tests for each replicate combination. To do this, replicates were manually switched one pair at a time across a comparison (sterilized versus non-sterilized soil or *Ms*-pC3 versus *Ms*-pML(int)) to assess whether results became nonsignificant.

Visualization of data was performed with the *ggplot2* package (96). The *tidyr* package was used to organize dataset nesting (97).

### Principal Component Analysis

Principal Component Analysis (PCA) plots were generated in R v4.4.1 (95) on 16S rRNA and shotgun metagenomic relative abundances. Species for which every microcosm had the same relative abundance were removed from the dataset since they contributed no information about variance (98). The PCA was performed and visualized using *factoextra* (99), *ggplot2* (96), and *cowplot* (100) packages.

### Data Availability

Relative abundance data from 16s rRNA sequencing is available on the SRA database under accession number PRJNA1189583 (https://www.ncbi.nlm.nih.gov/sra/?term=PRJNA1189583). Relative abundance data from shotgun metagenomic sequencing is available on the SRA database under accession number PRJNA1189582 (https://www.ncbi.nlm.nih.gov/sra/PRJNA1189582). All other data is included in the manuscript and supplementary materials or is available from the corresponding author upon request.

## Acknowledgements

This research was supported by the National Institute of Health through grant No. 1R15HD114135-01. We sincerely thank the Vice Provost for Research Dennis Manos for his financial and scientific support as well as the Dean of Arts and Science and Charles Center for student financial support. William & Mary’s High Performance Computing aided with computational resources. We also thank the 2023 William and Mary iGEM team, in particular Sofia Najjar and Lin Fang, for their assistance with the project.

## Supplemental Materials

**Table S1:**
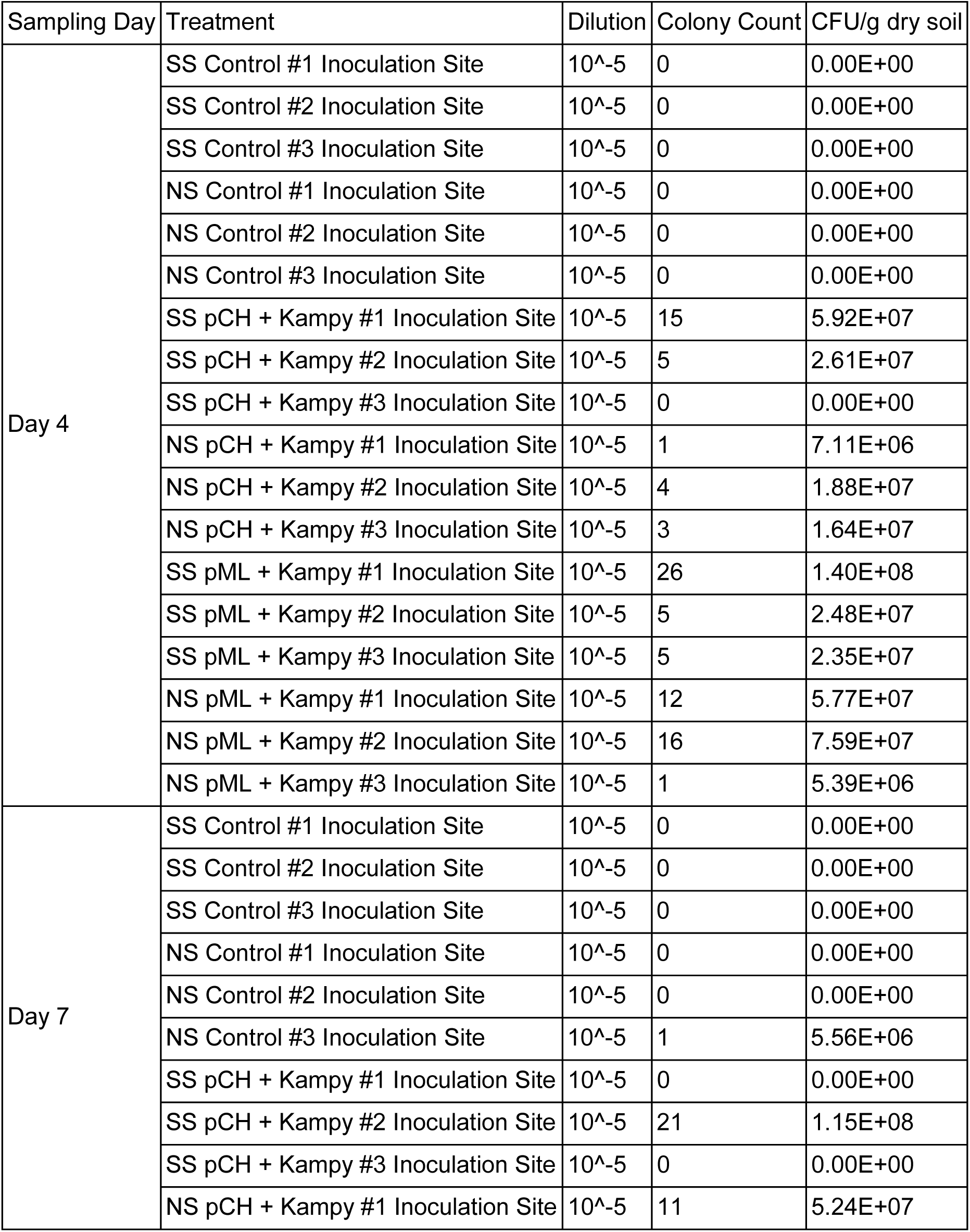
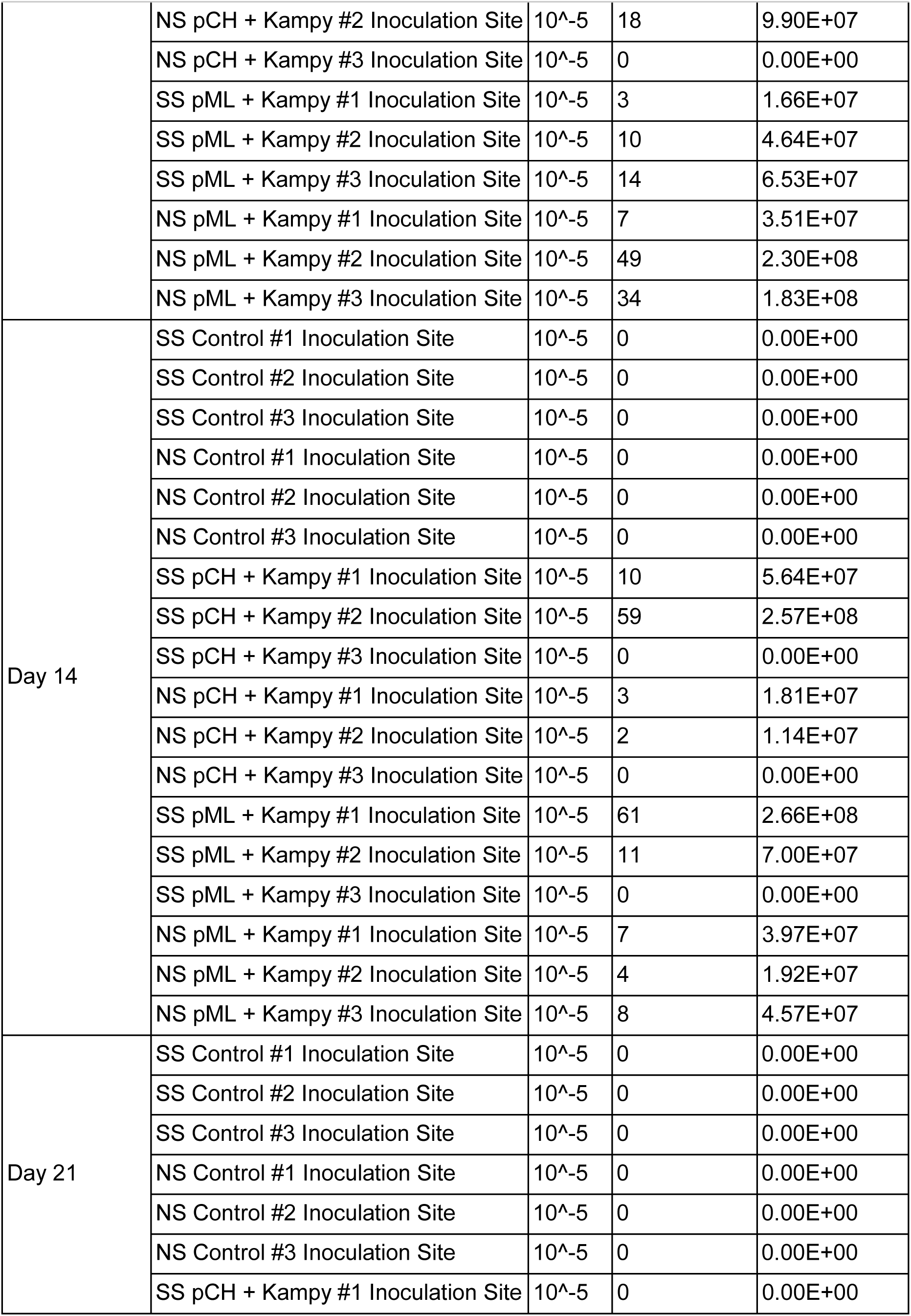
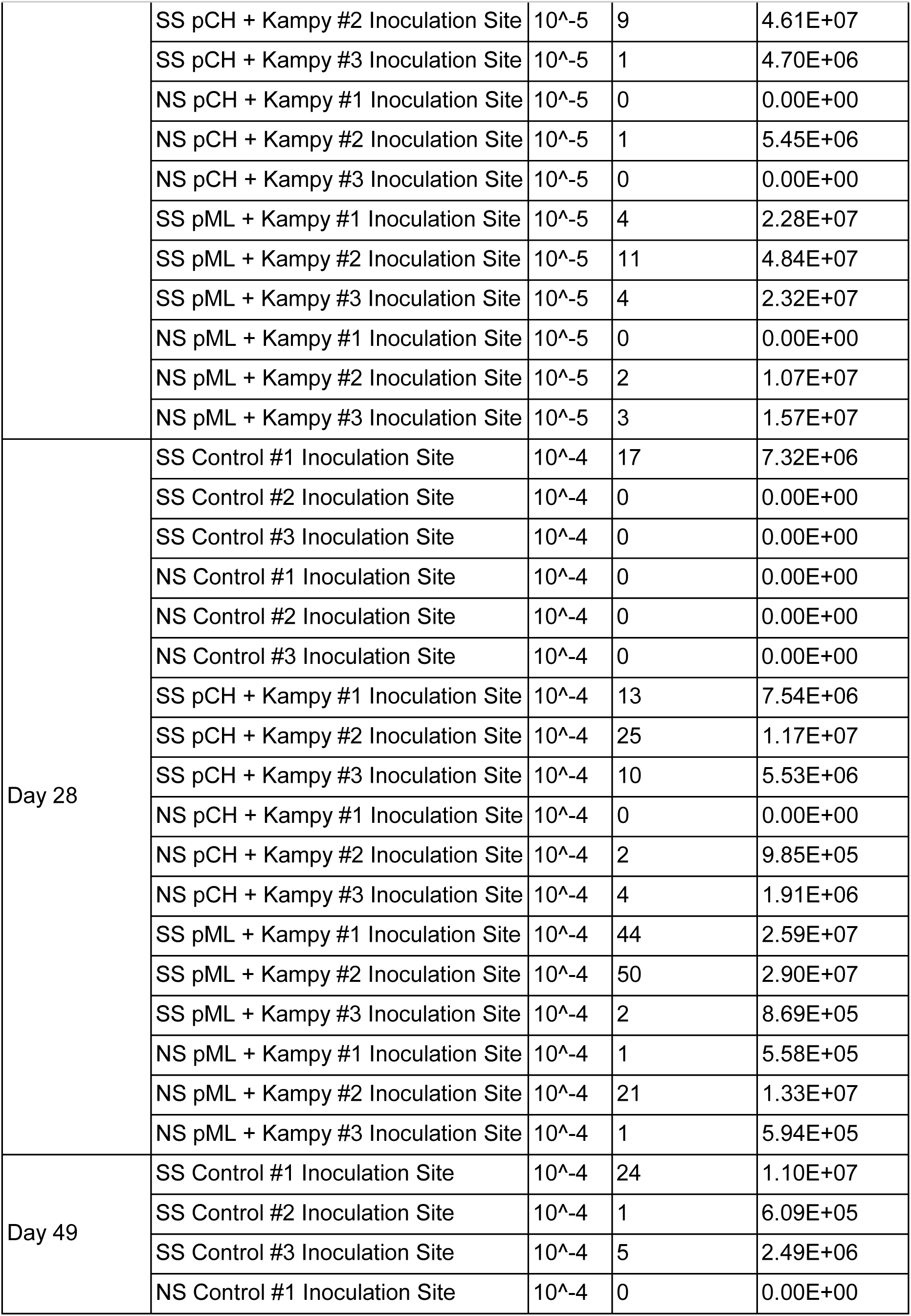
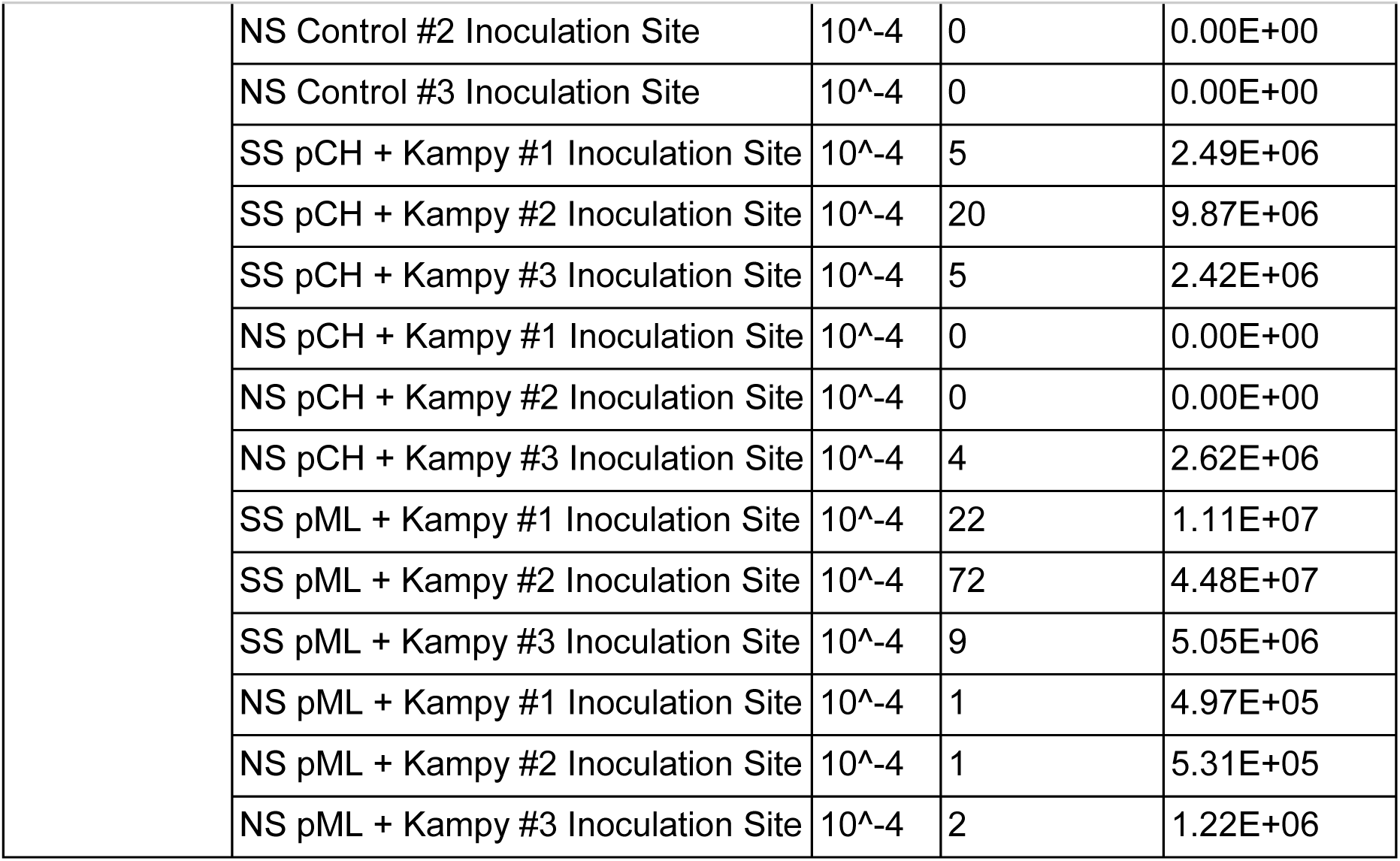
*M. smegmatis* colony counts and calculated CFU/g of dry soil at the inoculation site.

| Sampling Day | Treatment | Dilution | Colony Count | CFU/g dry soil |
| --- | --- | --- | --- | --- |
| Day 4 | SS Control #1 Inoculation Site | 10<sup>-5</sup> | 0 | 0.00E+00 |
|  | SS Control #2 Inoculation Site | 10<sup>-5</sup> | 0 | 0.00E+00 |
|  | SS Control #3 Inoculation Site | 10<sup>-5</sup> | 0 | 0.00E+00 |
|  | NS Control #1 Inoculation Site | 10<sup>-5</sup> | 0 | 0.00E+00 |
|  | NS Control #2 Inoculation Site | 10<sup>-5</sup> | 0 | 0.00E+00 |
|  | NS Control #3 Inoculation Site | 10<sup>-5</sup> | 0 | 0.00E+00 |
|  | SS pCH + Kampy #1 Inoculation Site | 10<sup>-5</sup> | 15 | 5.92E+07 |
|  | SS pCH + Kampy #2 Inoculation Site | 10<sup>-5</sup> | 5 | 2.61E+07 |
|  | SS pCH + Kampy #3 Inoculation Site | 10<sup>-5</sup> | 0 | 0.00E+00 |
|  | NS pCH + Kampy #1 Inoculation Site | 10<sup>-5</sup> | 1 | 7.11E+06 |
|  | NS pCH + Kampy #2 Inoculation Site | 10<sup>-5</sup> | 4 | 1.88E+07 |
|  | NS pCH + Kampy #3 Inoculation Site | 10<sup>-5</sup> | 3 | 1.64E+07 |
|  | SS pML + Kampy #1 Inoculation Site | 10<sup>-5</sup> | 26 | 1.40E+08 |
|  | SS pML + Kampy #2 Inoculation Site | 10<sup>-5</sup> | 5 | 2.48E+07 |
|  | SS pML + Kampy #3 Inoculation Site | 10<sup>-5</sup> | 5 | 2.35E+07 |
|  | NS pML + Kampy #1 Inoculation Site | 10<sup>-5</sup> | 12 | 5.77E+07 |
|  | NS pML + Kampy #2 Inoculation Site | 10<sup>-5</sup> | 16 | 7.59E+07 |
|  | NS pML + Kampy #3 Inoculation Site | 10<sup>-5</sup> | 1 | 5.39E+06 |
| Day 7 | SS Control #1 Inoculation Site | 10<sup>-5</sup> | 0 | 0.00E+00 |
|  | SS Control #2 Inoculation Site | 10<sup>-5</sup> | 0 | 0.00E+00 |
|  | SS Control #3 Inoculation Site | 10<sup>-5</sup> | 0 | 0.00E+00 |
|  | NS Control #1 Inoculation Site | 10<sup>-5</sup> | 0 | 0.00E+00 |
|  | NS Control #2 Inoculation Site | 10<sup>-5</sup> | 0 | 0.00E+00 |
|  | NS Control #3 Inoculation Site | 10<sup>-5</sup> | 1 | 5.56E+06 |
|  | SS pCH + Kampy #1 Inoculation Site | 10<sup>-5</sup> | 0 | 0.00E+00 |
|  | SS pCH + Kampy #2 Inoculation Site | 10<sup>-5</sup> | 21 | 1.15E+08 |
|  | SS pCH + Kampy #3 Inoculation Site | 10<sup>-5</sup> | 0 | 0.00E+00 |
|  | NS pCH + Kampy #1 Inoculation Site | 10<sup>-5</sup> | 11 | 5.24E+07 |
|  | NS pCH + Kampy #2 Inoculation Site | 10<sup>-5</sup> | 18 | 9.90E+07 |
|  | NS pCH + Kampy #3 Inoculation Site | 10<sup>-5</sup> | 0 | 0.00E+00 |
|  | SS pML + Kampy #1 Inoculation Site | 10<sup>-5</sup> | 3 | 1.66E+07 |
|  | SS pML + Kampy #2 Inoculation Site | 10<sup>-5</sup> | 10 | 4.64E+07 |
|  | SS pML + Kampy #3 Inoculation Site | 10<sup>-5</sup> | 14 | 6.53E+07 |
|  | NS pML + Kampy #1 Inoculation Site | 10<sup>-5</sup> | 7 | 3.51E+07 |
|  | NS pML + Kampy #2 Inoculation Site | 10<sup>-5</sup> | 49 | 2.30E+08 |
|  | NS pML + Kampy #3 Inoculation Site | 10<sup>-5</sup> | 34 | 1.83E+08 |
| Day 14 | SS Control #1 Inoculation Site | 10<sup>-5</sup> | 0 | 0.00E+00 |
|  | SS Control #2 Inoculation Site | 10<sup>-5</sup> | 0 | 0.00E+00 |
|  | SS Control #3 Inoculation Site | 10<sup>-5</sup> | 0 | 0.00E+00 |
|  | NS Control #1 Inoculation Site | 10<sup>-5</sup> | 0 | 0.00E+00 |
|  | NS Control #2 Inoculation Site | 10<sup>-5</sup> | 0 | 0.00E+00 |
|  | NS Control #3 Inoculation Site | 10<sup>-5</sup> | 0 | 0.00E+00 |
|  | SS pCH + Kampy #1 Inoculation Site | 10<sup>-5</sup> | 10 | 5.64E+07 |
|  | SS pCH + Kampy #2 Inoculation Site | 10<sup>-5</sup> | 59 | 2.57E+08 |
|  | SS pCH + Kampy #3 Inoculation Site | 10<sup>-5</sup> | 0 | 0.00E+00 |
|  | NS pCH + Kampy #1 Inoculation Site | 10<sup>-5</sup> | 3 | 1.81E+07 |
|  | NS pCH + Kampy #2 Inoculation Site | 10<sup>-5</sup> | 2 | 1.14E+07 |
|  | NS pCH + Kampy #3 Inoculation Site | 10<sup>-5</sup> | 0 | 0.00E+00 |
|  | SS pML + Kampy #1 Inoculation Site | 10<sup>-5</sup> | 61 | 2.66E+08 |
|  | SS pML + Kampy #2 Inoculation Site | 10<sup>-5</sup> | 11 | 7.00E+07 |
|  | SS pML + Kampy #3 Inoculation Site | 10<sup>-5</sup> | 0 | 0.00E+00 |
|  | NS pML + Kampy #1 Inoculation Site | 10<sup>-5</sup> | 7 | 3.97E+07 |
|  | NS pML + Kampy #2 Inoculation Site | 10<sup>-5</sup> | 4 | 1.92E+07 |
|  | NS pML + Kampy #3 Inoculation Site | 10<sup>-5</sup> | 8 | 4.57E+07 |
| Day 21 | SS Control #1 Inoculation Site | 10<sup>-5</sup> | 0 | 0.00E+00 |
|  | SS Control #2 Inoculation Site | 10<sup>-5</sup> | 0 | 0.00E+00 |
|  | SS Control #3 Inoculation Site | 10<sup>-5</sup> | 0 | 0.00E+00 |
|  | NS Control #1 Inoculation Site | 10<sup>-5</sup> | 0 | 0.00E+00 |
|  | NS Control #2 Inoculation Site | 10<sup>-5</sup> | 0 | 0.00E+00 |
|  | NS Control #3 Inoculation Site | 10<sup>-5</sup> | 0 | 0.00E+00 |
|  | SS pCH + Kampy #1 Inoculation Site | 10<sup>-5</sup> | 0 | 0.00E+00 |
|  | SS pCH + Kampy #2 Inoculation Site | 10<sup>-5</sup> | 9 | 4.61E+07 |
|  | SS pCH + Kampy #3 Inoculation Site | 10<sup>-5</sup> | 1 | 4.70E+06 |
|  | NS pCH + Kampy #1 Inoculation Site | 10<sup>-5</sup> | 0 | 0.00E+00 |
|  | NS pCH + Kampy #2 Inoculation Site | 10<sup>-5</sup> | 1 | 5.45E+06 |
|  | NS pCH + Kampy #3 Inoculation Site | 10<sup>-5</sup> | 0 | 0.00E+00 |
|  | SS pML + Kampy #1 Inoculation Site | 10<sup>-5</sup> | 4 | 2.28E+07 |
|  | SS pML + Kampy #2 Inoculation Site | 10<sup>-5</sup> | 11 | 4.84E+07 |
|  | SS pML + Kampy #3 Inoculation Site | 10<sup>-5</sup> | 4 | 2.32E+07 |
|  | NS pML + Kampy #1 Inoculation Site | 10<sup>-5</sup> | 0 | 0.00E+00 |
|  | NS pML + Kampy #2 Inoculation Site | 10<sup>-5</sup> | 2 | 1.07E+07 |
|  | NS pML + Kampy #3 Inoculation Site | 10<sup>-5</sup> | 3 | 1.57E+07 |
| Day 28 | SS Control #1 Inoculation Site | 10<sup>-4</sup> | 17 | 7.32E+06 |
|  | SS Control #2 Inoculation Site | 10<sup>-4</sup> | 0 | 0.00E+00 |
|  | SS Control #3 Inoculation Site | 10<sup>-4</sup> | 0 | 0.00E+00 |
|  | NS Control #1 Inoculation Site | 10<sup>-4</sup> | 0 | 0.00E+00 |
|  | NS Control #2 Inoculation Site | 10<sup>-4</sup> | 0 | 0.00E+00 |
|  | NS Control #3 Inoculation Site | 10<sup>-4</sup> | 0 | 0.00E+00 |
|  | SS pCH + Kampy #1 Inoculation Site | 10<sup>-4</sup> | 13 | 7.54E+06 |
|  | SS pCH + Kampy #2 Inoculation Site | 10<sup>-4</sup> | 25 | 1.17E+07 |
|  | SS pCH + Kampy #3 Inoculation Site | 10<sup>-4</sup> | 10 | 5.53E+06 |
|  | NS pCH + Kampy #1 Inoculation Site | 10<sup>-4</sup> | 0 | 0.00E+00 |
|  | NS pCH + Kampy #2 Inoculation Site | 10<sup>-4</sup> | 2 | 9.85E+05 |
|  | NS pCH + Kampy #3 Inoculation Site | 10<sup>-4</sup> | 4 | 1.91E+06 |
|  | SS pML + Kampy #1 Inoculation Site | 10<sup>-4</sup> | 44 | 2.59E+07 |
|  | SS pML + Kampy #2 Inoculation Site | 10<sup>-4</sup> | 50 | 2.90E+07 |
|  | SS pML + Kampy #3 Inoculation Site | 10<sup>-4</sup> | 2 | 8.69E+05 |
|  | NS pML + Kampy #1 Inoculation Site | 10<sup>-4</sup> | 1 | 5.58E+05 |
|  | NS pML + Kampy #2 Inoculation Site | 10<sup>-4</sup> | 21 | 1.33E+07 |
|  | NS pML + Kampy #3 Inoculation Site | 10<sup>-4</sup> | 1 | 5.94E+05 |
| Day 49 | SS Control #1 Inoculation Site | 10<sup>-4</sup> | 24 | 1.10E+07 |
|  | SS Control #2 Inoculation Site | 10<sup>-4</sup> | 1 | 6.09E+05 |
|  | SS Control #3 Inoculation Site | 10<sup>-4</sup> | 5 | 2.49E+06 |
|  | NS Control #1 Inoculation Site | 10<sup>-4</sup> | 0 | 0.00E+00 |
|  | NS Control #2 Inoculation Site | 10<sup>-4</sup> | 0 | 0.00E+00 |
|  | NS Control #3 Inoculation Site | 10<sup>-4</sup> | 0 | 0.00E+00 |
|  | SS pCH + Kamy #1 Inoculation Site | 10<sup>-4</sup> | 5 | 2.49E+06 |
|  | SS pCH + Kamy #2 Inoculation Site | 10<sup>-4</sup> | 20 | 9.87E+06 |
|  | SS pCH + Kamy #3 Inoculation Site | 10<sup>-4</sup> | 5 | 2.42E+06 |
|  | NS pCH + Kamy #1 Inoculation Site | 10<sup>-4</sup> | 0 | 0.00E+00 |
|  | NS pCH + Kamy #2 Inoculation Site | 10<sup>-4</sup> | 0 | 0.00E+00 |
|  | NS pCH + Kamy #3 Inoculation Site | 10<sup>-4</sup> | 4 | 2.62E+06 |
|  | SS pML + Kamy #1 Inoculation Site | 10<sup>-4</sup> | 22 | 1.11E+07 |
|  | SS pML + Kamy #2 Inoculation Site | 10<sup>-4</sup> | 72 | 4.48E+07 |
|  | SS pML + Kamy #3 Inoculation Site | 10<sup>-4</sup> | 9 | 5.05E+06 |
|  | NS pML + Kamy #1 Inoculation Site | 10<sup>-4</sup> | 1 | 4.97E+05 |
|  | NS pML + Kamy #2 Inoculation Site | 10<sup>-4</sup> | 1 | 5.31E+05 |
|  | NS pML + Kamy #3 Inoculation Site | 10<sup>-4</sup> | 2 | 1.22E+06 |

**Table S2:**
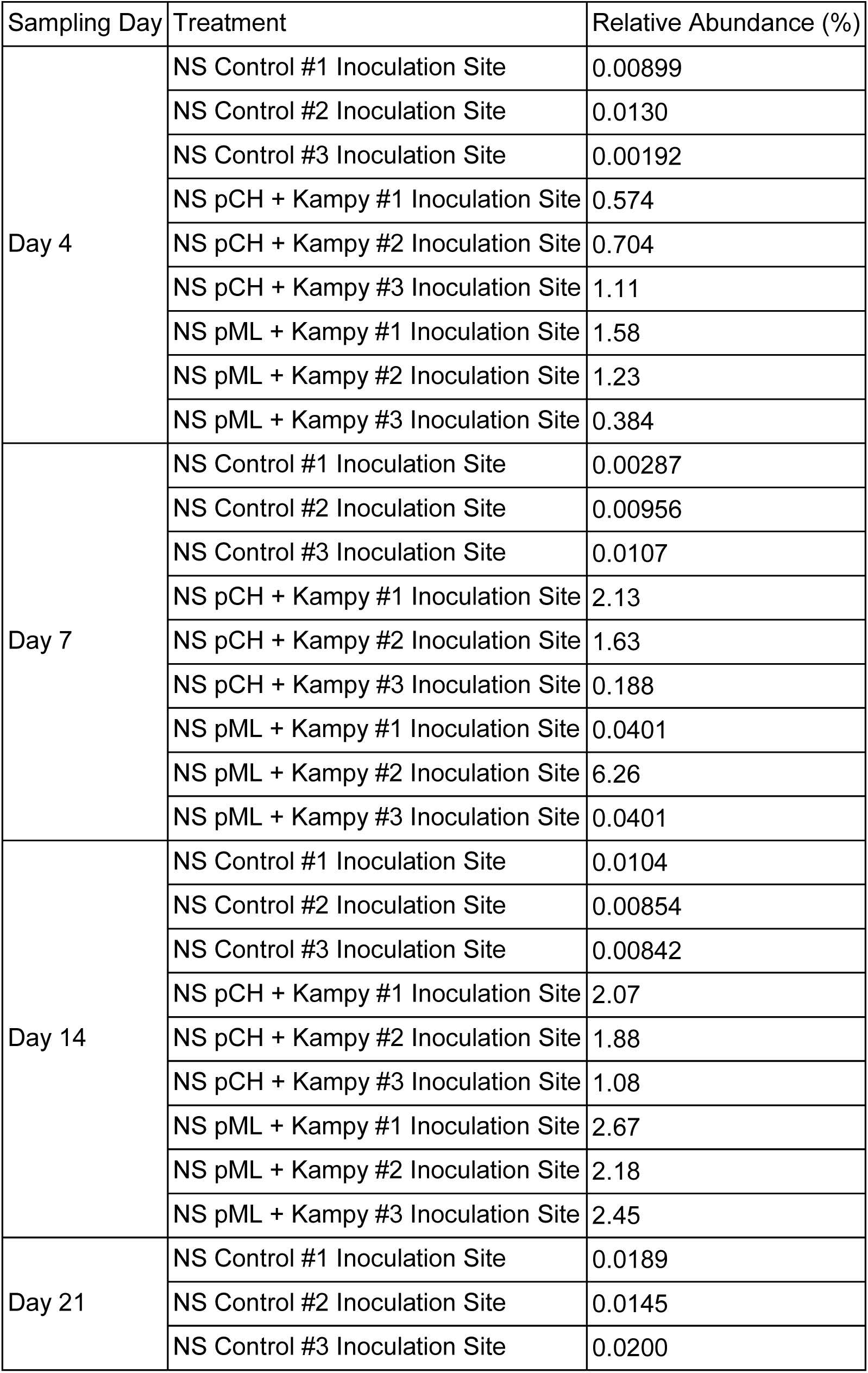
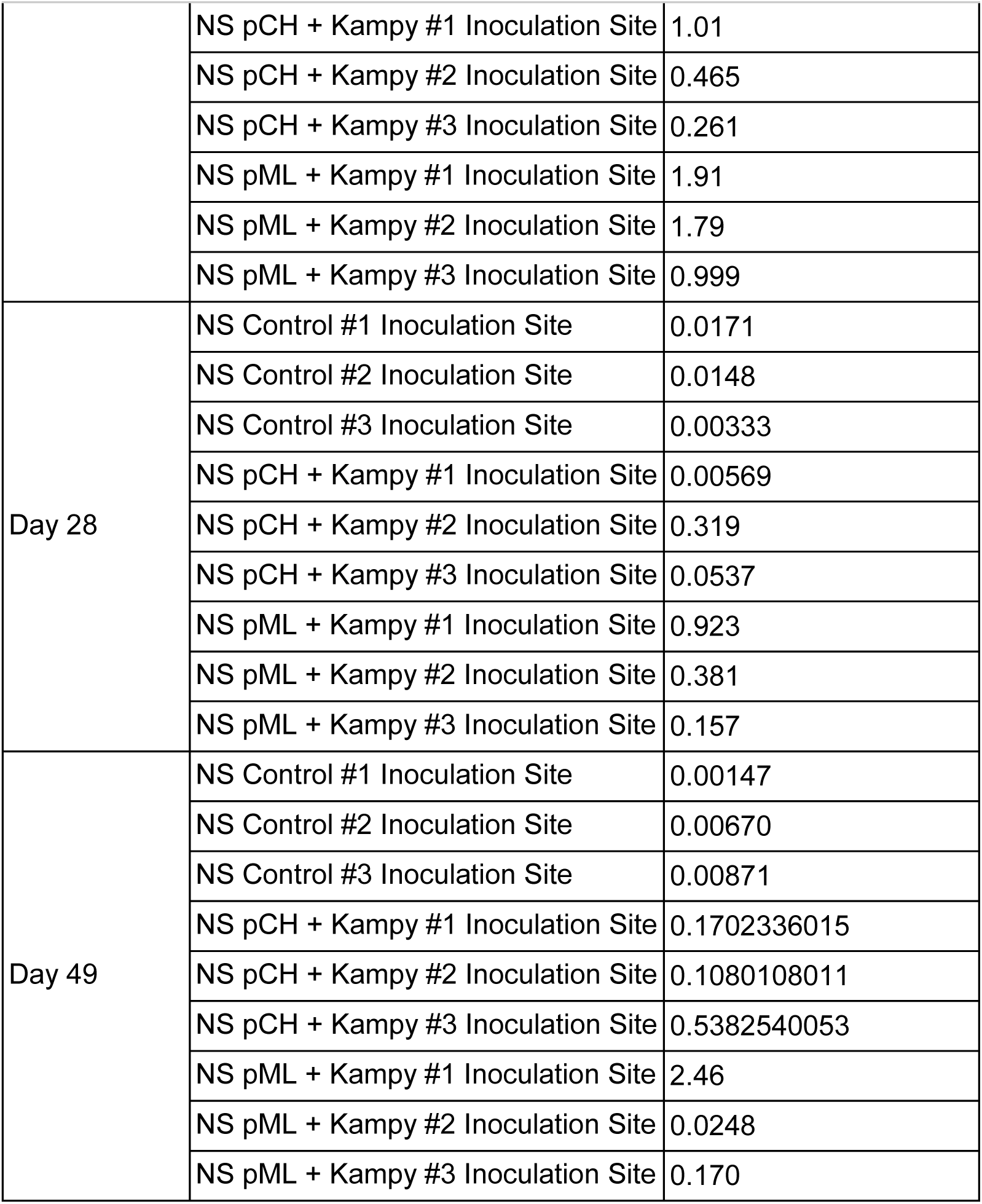
*M. smegmatis* 16s relative abundance at the inoculation site.

| Sampling Day | Treatment | Relative Abundance (%) |
| --- | --- | --- |
| Day 4 | NS Control #1 Inoculation Site | 0.00899 |
|  | NS Control #2 Inoculation Site | 0.0130 |
|  | NS Control #3 Inoculation Site | 0.00192 |
|  | NS pCH + Kamy #1 Inoculation Site | 0.574 |
|  | NS pCH + Kamy #2 Inoculation Site | 0.704 |
|  | NS pCH + Kamy #3 Inoculation Site | 1.11 |
|  | NS pML + Kamy #1 Inoculation Site | 1.58 |
|  | NS pML + Kamy #2 Inoculation Site | 1.23 |
|  | NS pML + Kamy #3 Inoculation Site | 0.384 |
| Day 7 | NS Control #1 Inoculation Site | 0.00287 |
|  | NS Control #2 Inoculation Site | 0.00956 |
|  | NS Control #3 Inoculation Site | 0.0107 |
|  | NS pCH + Kamy #1 Inoculation Site | 2.13 |
|  | NS pCH + Kamy #2 Inoculation Site | 1.63 |
|  | NS pCH + Kamy #3 Inoculation Site | 0.188 |
|  | NS pML + Kamy #1 Inoculation Site | 0.0401 |
|  | NS pML + Kamy #2 Inoculation Site | 6.26 |
|  | NS pML + Kamy #3 Inoculation Site | 0.0401 |
| Day 14 | NS Control #1 Inoculation Site | 0.0104 |
|  | NS Control #2 Inoculation Site | 0.00854 |
|  | NS Control #3 Inoculation Site | 0.00842 |
|  | NS pCH + Kamy #1 Inoculation Site | 2.07 |
|  | NS pCH + Kamy #2 Inoculation Site | 1.88 |
|  | NS pCH + Kamy #3 Inoculation Site | 1.08 |
|  | NS pML + Kamy #1 Inoculation Site | 2.67 |
|  | NS pML + Kamy #2 Inoculation Site | 2.18 |
|  | NS pML + Kamy #3 Inoculation Site | 2.45 |
| Day 21 | NS Control #1 Inoculation Site | 0.0189 |
|  | NS Control #2 Inoculation Site | 0.0145 |
|  | NS Control #3 Inoculation Site | 0.0200 |
|  | NS pCH + Kamy #1 Inoculation Site | 1.01 |
|  | NS pCH + Kamy #2 Inoculation Site | 0.465 |
|  | NS pCH + Kamy #3 Inoculation Site | 0.261 |
|  | NS pML + Kamy #1 Inoculation Site | 1.91 |
|  | NS pML + Kamy #2 Inoculation Site | 1.79 |
|  | NS pML + Kamy #3 Inoculation Site | 0.999 |
| Day 28 | NS Control #1 Inoculation Site | 0.0171 |
|  | NS Control #2 Inoculation Site | 0.0148 |
|  | NS Control #3 Inoculation Site | 0.00333 |
|  | NS pCH + Kamy #1 Inoculation Site | 0.00569 |
|  | NS pCH + Kamy #2 Inoculation Site | 0.319 |
|  | NS pCH + Kamy #3 Inoculation Site | 0.0537 |
|  | NS pML + Kamy #1 Inoculation Site | 0.923 |
|  | NS pML + Kamy #2 Inoculation Site | 0.381 |
|  | NS pML + Kamy #3 Inoculation Site | 0.157 |
| Day 49 | NS Control #1 Inoculation Site | 0.00147 |
|  | NS Control #2 Inoculation Site | 0.00670 |
|  | NS Control #3 Inoculation Site | 0.00871 |
|  | NS pCH + Kamy #1 Inoculation Site | 0.1702336015 |
|  | NS pCH + Kamy #2 Inoculation Site | 0.1080108011 |
|  | NS pCH + Kamy #3 Inoculation Site | 0.5382540053 |
|  | NS pML + Kamy #1 Inoculation Site | 2.46 |
|  | NS pML + Kamy #2 Inoculation Site | 0.0248 |
|  | NS pML + Kamy #3 Inoculation Site | 0.170 |

**Table S3:**
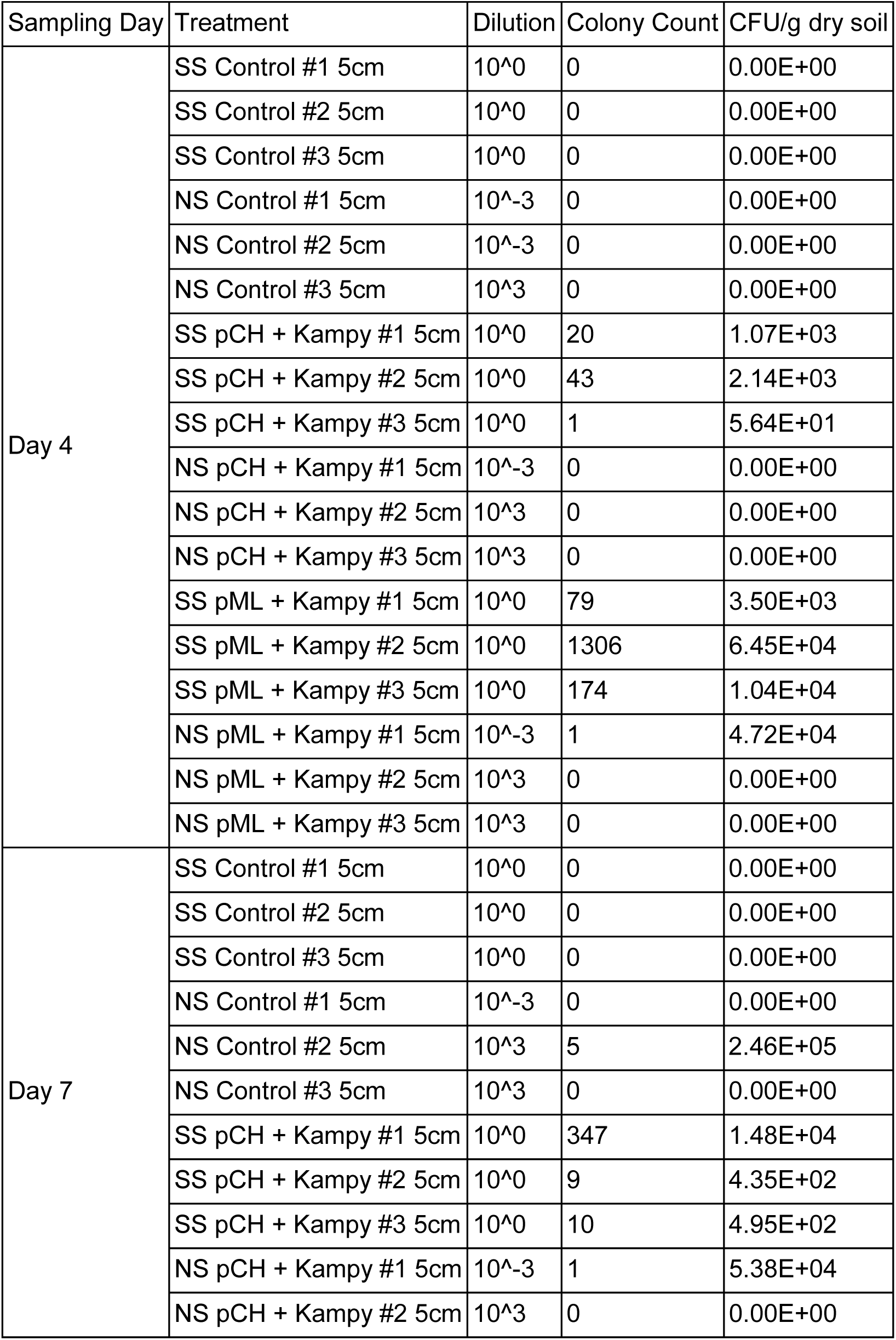
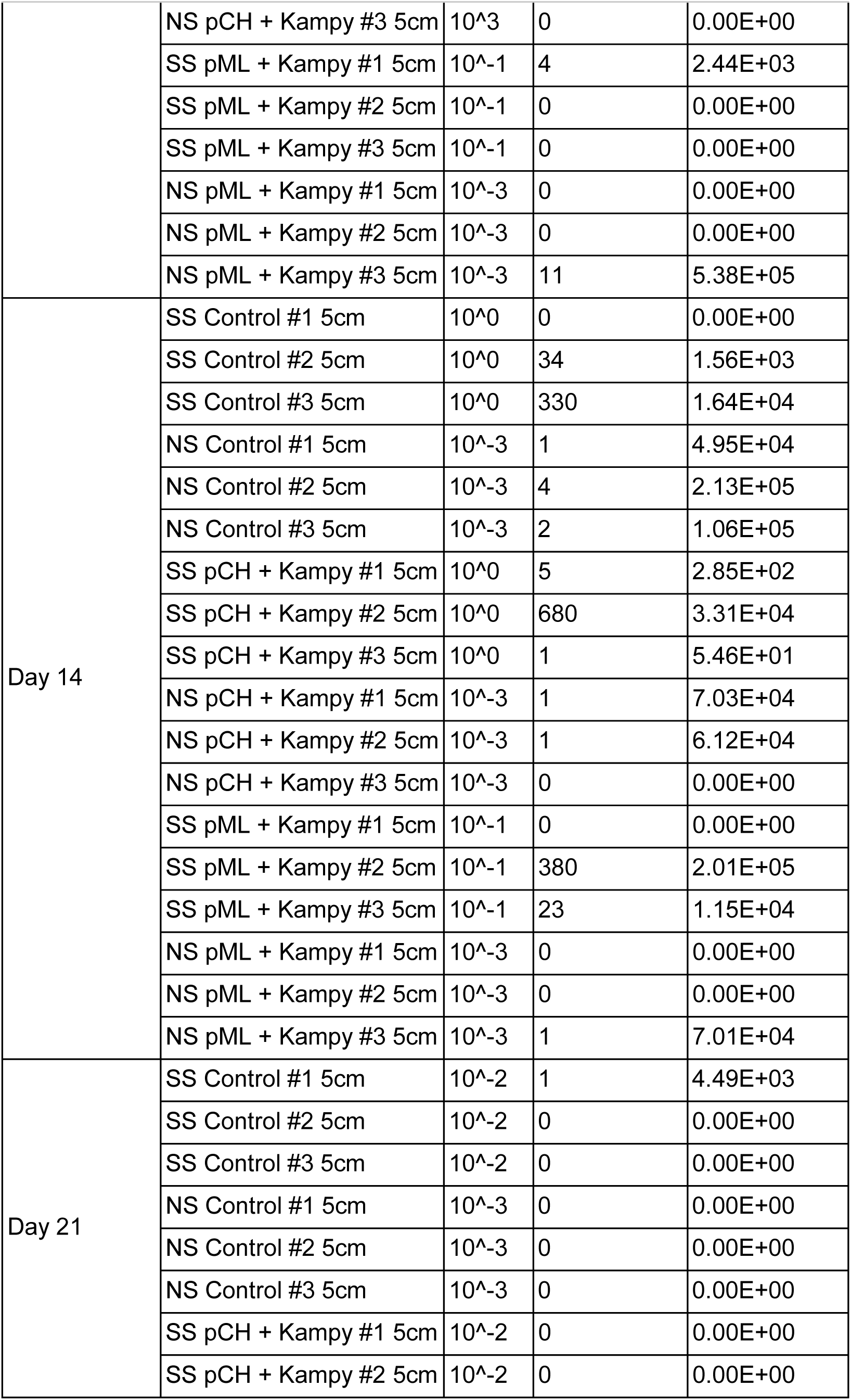
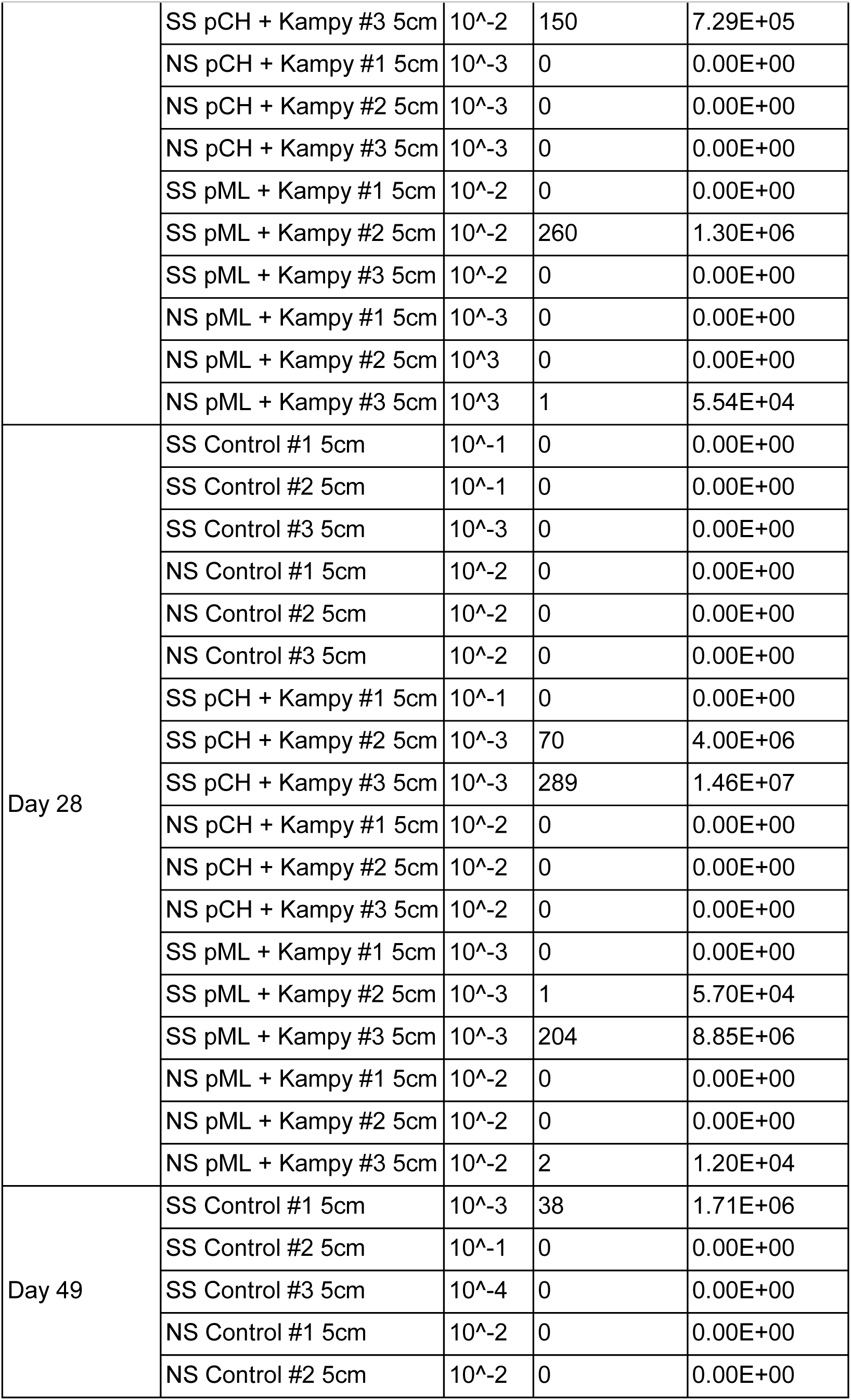
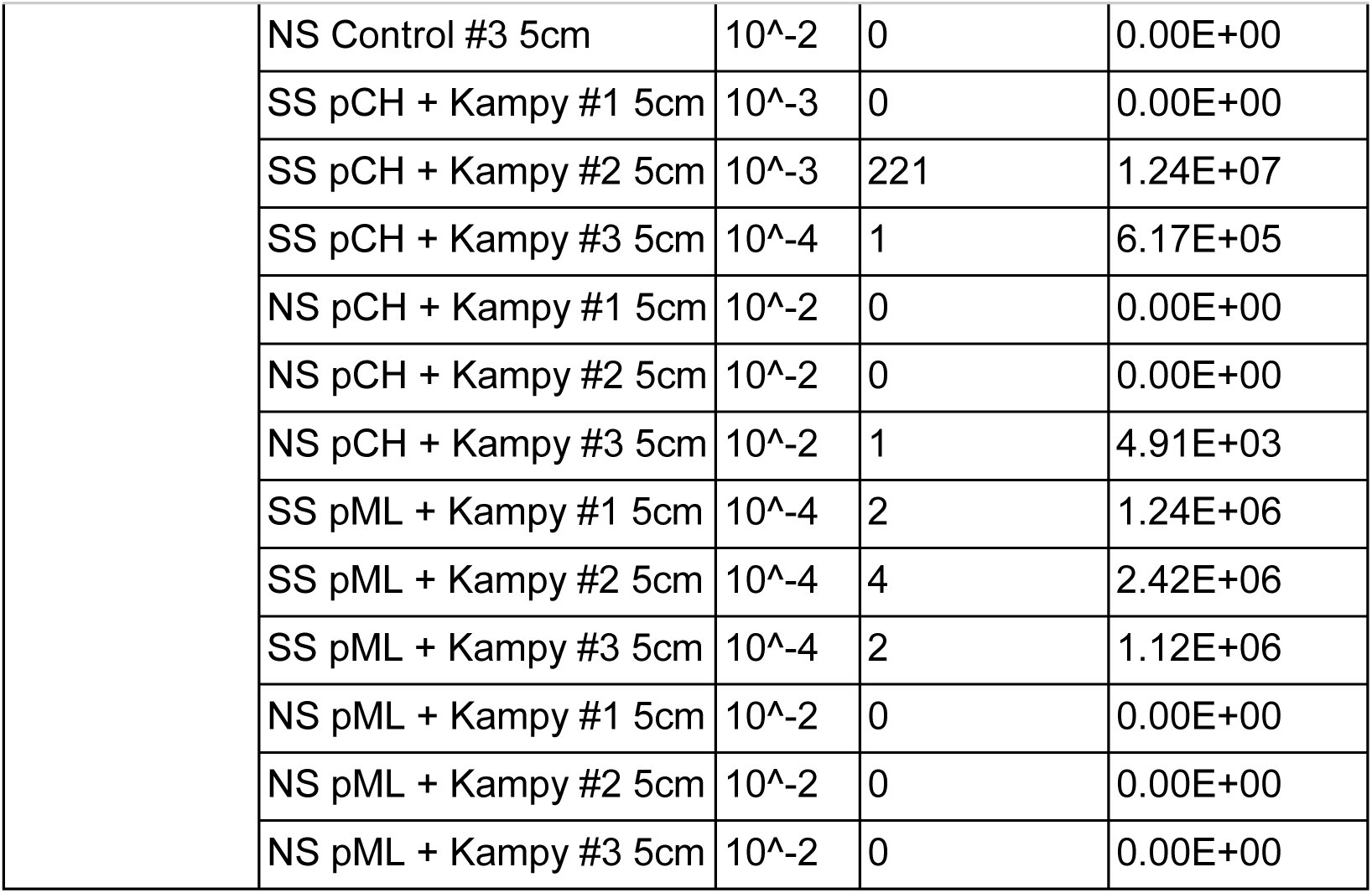
*M. smegmatis* colony counts and calculated CFU/g of dry soil at 5 cm away from the inoculation site.

| Sampling Day | Treatment | Dilution | Colony Count | CFU/g dry soil |
| --- | --- | --- | --- | --- |
| Day 4 | SS Control #1 5cm | 10<sup>0</sup> | 0 | 0.00E+00 |
|  | SS Control #2 5cm | 10<sup>0</sup> | 0 | 0.00E+00 |
|  | SS Control #3 5cm | 10<sup>0</sup> | 0 | 0.00E+00 |
|  | NS Control #1 5cm | 10<sup>-3</sup> | 0 | 0.00E+00 |
|  | NS Control #2 5cm | 10<sup>-3</sup> | 0 | 0.00E+00 |
|  | NS Control #3 5cm | 10<sup>3</sup> | 0 | 0.00E+00 |
|  | SS pCH + Kampy #1 5cm | 10<sup>0</sup> | 20 | 1.07E+03 |
|  | SS pCH + Kampy #2 5cm | 10<sup>0</sup> | 43 | 2.14E+03 |
|  | SS pCH + Kampy #3 5cm | 10<sup>0</sup> | 1 | 5.64E+01 |
|  | NS pCH + Kampy #1 5cm | 10<sup>-3</sup> | 0 | 0.00E+00 |
|  | NS pCH + Kampy #2 5cm | 10<sup>3</sup> | 0 | 0.00E+00 |
|  | NS pCH + Kampy #3 5cm | 10<sup>3</sup> | 0 | 0.00E+00 |
|  | SS pML + Kampy #1 5cm | 10<sup>0</sup> | 79 | 3.50E+03 |
|  | SS pML + Kampy #2 5cm | 10<sup>0</sup> | 1306 | 6.45E+04 |
|  | SS pML + Kampy #3 5cm | 10<sup>0</sup> | 174 | 1.04E+04 |
|  | NS pML + Kampy #1 5cm | 10<sup>-3</sup> | 1 | 4.72E+04 |
|  | NS pML + Kampy #2 5cm | 10<sup>3</sup> | 0 | 0.00E+00 |
|  | NS pML + Kampy #3 5cm | 10<sup>3</sup> | 0 | 0.00E+00 |
| Day 7 | SS Control #1 5cm | 10<sup>0</sup> | 0 | 0.00E+00 |
|  | SS Control #2 5cm | 10<sup>0</sup> | 0 | 0.00E+00 |
|  | SS Control #3 5cm | 10<sup>0</sup> | 0 | 0.00E+00 |
|  | NS Control #1 5cm | 10<sup>-3</sup> | 0 | 0.00E+00 |
|  | NS Control #2 5cm | 10<sup>3</sup> | 5 | 2.46E+05 |
|  | NS Control #3 5cm | 10<sup>3</sup> | 0 | 0.00E+00 |
|  | SS pCH + Kampy #1 5cm | 10<sup>0</sup> | 347 | 1.48E+04 |
|  | SS pCH + Kampy #2 5cm | 10<sup>0</sup> | 9 | 4.35E+02 |
|  | SS pCH + Kampy #3 5cm | 10<sup>0</sup> | 10 | 4.95E+02 |
|  | NS pCH + Kampy #1 5cm | 10<sup>-3</sup> | 1 | 5.38E+04 |
|  | NS pCH + Kampy #2 5cm | 10<sup>3</sup> | 0 | 0.00E+00 |
|  | NS pCH + Kampy #3 5cm | 10<sup>3</sup> | 0 | 0.00E+00 |
|  | SS pML + Kampy #1 5cm | 10<sup>-1</sup> | 4 | 2.44E+03 |
|  | SS pML + Kampy #2 5cm | 10<sup>-1</sup> | 0 | 0.00E+00 |
|  | SS pML + Kampy #3 5cm | 10<sup>-1</sup> | 0 | 0.00E+00 |
|  | NS pML + Kampy #1 5cm | 10<sup>-3</sup> | 0 | 0.00E+00 |
|  | NS pML + Kampy #2 5cm | 10<sup>-3</sup> | 0 | 0.00E+00 |
|  | NS pML + Kampy #3 5cm | 10<sup>-3</sup> | 11 | 5.38E+05 |
| Day 14 | SS Control #1 5cm | 10<sup>0</sup> | 0 | 0.00E+00 |
|  | SS Control #2 5cm | 10<sup>0</sup> | 34 | 1.56E+03 |
|  | SS Control #3 5cm | 10<sup>0</sup> | 330 | 1.64E+04 |
|  | NS Control #1 5cm | 10<sup>-3</sup> | 1 | 4.95E+04 |
|  | NS Control #2 5cm | 10<sup>-3</sup> | 4 | 2.13E+05 |
|  | NS Control #3 5cm | 10<sup>-3</sup> | 2 | 1.06E+05 |
|  | SS pCH + Kampy #1 5cm | 10<sup>0</sup> | 5 | 2.85E+02 |
|  | SS pCH + Kampy #2 5cm | 10<sup>0</sup> | 680 | 3.31E+04 |
|  | SS pCH + Kampy #3 5cm | 10<sup>0</sup> | 1 | 5.46E+01 |
|  | NS pCH + Kampy #1 5cm | 10<sup>-3</sup> | 1 | 7.03E+04 |
|  | NS pCH + Kampy #2 5cm | 10<sup>-3</sup> | 1 | 6.12E+04 |
|  | NS pCH + Kampy #3 5cm | 10<sup>-3</sup> | 0 | 0.00E+00 |
|  | SS pML + Kampy #1 5cm | 10<sup>-1</sup> | 0 | 0.00E+00 |
|  | SS pML + Kampy #2 5cm | 10<sup>-1</sup> | 380 | 2.01E+05 |
|  | SS pML + Kampy #3 5cm | 10<sup>-1</sup> | 23 | 1.15E+04 |
|  | NS pML + Kampy #1 5cm | 10<sup>-3</sup> | 0 | 0.00E+00 |
|  | NS pML + Kampy #2 5cm | 10<sup>-3</sup> | 0 | 0.00E+00 |
|  | NS pML + Kampy #3 5cm | 10<sup>-3</sup> | 1 | 7.01E+04 |
| Day 21 | SS Control #1 5cm | 10<sup>-2</sup> | 1 | 4.49E+03 |
|  | SS Control #2 5cm | 10<sup>-2</sup> | 0 | 0.00E+00 |
|  | SS Control #3 5cm | 10<sup>-2</sup> | 0 | 0.00E+00 |
|  | NS Control #1 5cm | 10<sup>-3</sup> | 0 | 0.00E+00 |
|  | NS Control #2 5cm | 10<sup>-3</sup> | 0 | 0.00E+00 |
|  | NS Control #3 5cm | 10<sup>-3</sup> | 0 | 0.00E+00 |
|  | SS pCH + Kampy #1 5cm | 10<sup>-2</sup> | 0 | 0.00E+00 |
|  | SS pCH + Kampy #2 5cm | 10<sup>-2</sup> | 0 | 0.00E+00 |
|  | SS pCH + Kampy #3 5cm | 10<sup>-2</sup> | 150 | 7.29E+05 |
|  | NS pCH + Kampy #1 5cm | 10<sup>-3</sup> | 0 | 0.00E+00 |
|  | NS pCH + Kampy #2 5cm | 10<sup>-3</sup> | 0 | 0.00E+00 |
|  | NS pCH + Kampy #3 5cm | 10<sup>-3</sup> | 0 | 0.00E+00 |
|  | SS pML + Kampy #1 5cm | 10<sup>-2</sup> | 0 | 0.00E+00 |
|  | SS pML + Kampy #2 5cm | 10<sup>-2</sup> | 260 | 1.30E+06 |
|  | SS pML + Kampy #3 5cm | 10<sup>-2</sup> | 0 | 0.00E+00 |
|  | NS pML + Kampy #1 5cm | 10<sup>-3</sup> | 0 | 0.00E+00 |
|  | NS pML + Kampy #2 5cm | 10<sup>-3</sup> | 0 | 0.00E+00 |
|  | NS pML + Kampy #3 5cm | 10<sup>-3</sup> | 1 | 5.54E+04 |
| Day 28 | SS Control #1 5cm | 10<sup>-1</sup> | 0 | 0.00E+00 |
|  | SS Control #2 5cm | 10<sup>-1</sup> | 0 | 0.00E+00 |
|  | SS Control #3 5cm | 10<sup>-3</sup> | 0 | 0.00E+00 |
|  | NS Control #1 5cm | 10<sup>-2</sup> | 0 | 0.00E+00 |
|  | NS Control #2 5cm | 10<sup>-2</sup> | 0 | 0.00E+00 |
|  | NS Control #3 5cm | 10<sup>-2</sup> | 0 | 0.00E+00 |
|  | SS pCH + Kampy #1 5cm | 10<sup>-1</sup> | 0 | 0.00E+00 |
|  | SS pCH + Kampy #2 5cm | 10<sup>-3</sup> | 70 | 4.00E+06 |
|  | SS pCH + Kampy #3 5cm | 10<sup>-3</sup> | 289 | 1.46E+07 |
|  | NS pCH + Kampy #1 5cm | 10<sup>-2</sup> | 0 | 0.00E+00 |
|  | NS pCH + Kampy #2 5cm | 10<sup>-2</sup> | 0 | 0.00E+00 |
|  | NS pCH + Kampy #3 5cm | 10<sup>-2</sup> | 0 | 0.00E+00 |
|  | SS pML + Kampy #1 5cm | 10<sup>-3</sup> | 0 | 0.00E+00 |
|  | SS pML + Kampy #2 5cm | 10<sup>-3</sup> | 1 | 5.70E+04 |
|  | SS pML + Kampy #3 5cm | 10<sup>-3</sup> | 204 | 8.85E+06 |
|  | NS pML + Kampy #1 5cm | 10<sup>-2</sup> | 0 | 0.00E+00 |
|  | NS pML + Kampy #2 5cm | 10<sup>-2</sup> | 0 | 0.00E+00 |
|  | NS pML + Kampy #3 5cm | 10<sup>-2</sup> | 2 | 1.20E+04 |
| Day 49 | SS Control #1 5cm | 10<sup>-3</sup> | 38 | 1.71E+06 |
|  | SS Control #2 5cm | 10<sup>-1</sup> | 0 | 0.00E+00 |
|  | SS Control #3 5cm | 10<sup>-4</sup> | 0 | 0.00E+00 |
|  | NS Control #1 5cm | 10<sup>-2</sup> | 0 | 0.00E+00 |
|  | NS Control #2 5cm | 10<sup>-2</sup> | 0 | 0.00E+00 |
|  | NS Control #3 5cm | 10<sup>-2</sup> | 0 | 0.00E+00 |
|  | SS pCH + Kamy #1 5cm | 10<sup>-3</sup> | 0 | 0.00E+00 |
|  | SS pCH + Kamy #2 5cm | 10<sup>-3</sup> | 221 | 1.24E+07 |
|  | SS pCH + Kamy #3 5cm | 10<sup>-4</sup> | 1 | 6.17E+05 |
|  | NS pCH + Kamy #1 5cm | 10<sup>-2</sup> | 0 | 0.00E+00 |
|  | NS pCH + Kamy #2 5cm | 10<sup>-2</sup> | 0 | 0.00E+00 |
|  | NS pCH + Kamy #3 5cm | 10<sup>-2</sup> | 1 | 4.91E+03 |
|  | SS pML + Kamy #1 5cm | 10<sup>-4</sup> | 2 | 1.24E+06 |
|  | SS pML + Kamy #2 5cm | 10<sup>-4</sup> | 4 | 2.42E+06 |
|  | SS pML + Kamy #3 5cm | 10<sup>-4</sup> | 2 | 1.12E+06 |
|  | NS pML + Kamy #1 5cm | 10<sup>-2</sup> | 0 | 0.00E+00 |
|  | NS pML + Kamy #2 5cm | 10<sup>-2</sup> | 0 | 0.00E+00 |
|  | NS pML + Kamy #3 5cm | 10<sup>-2</sup> | 0 | 0.00E+00 |

**Table S4:**
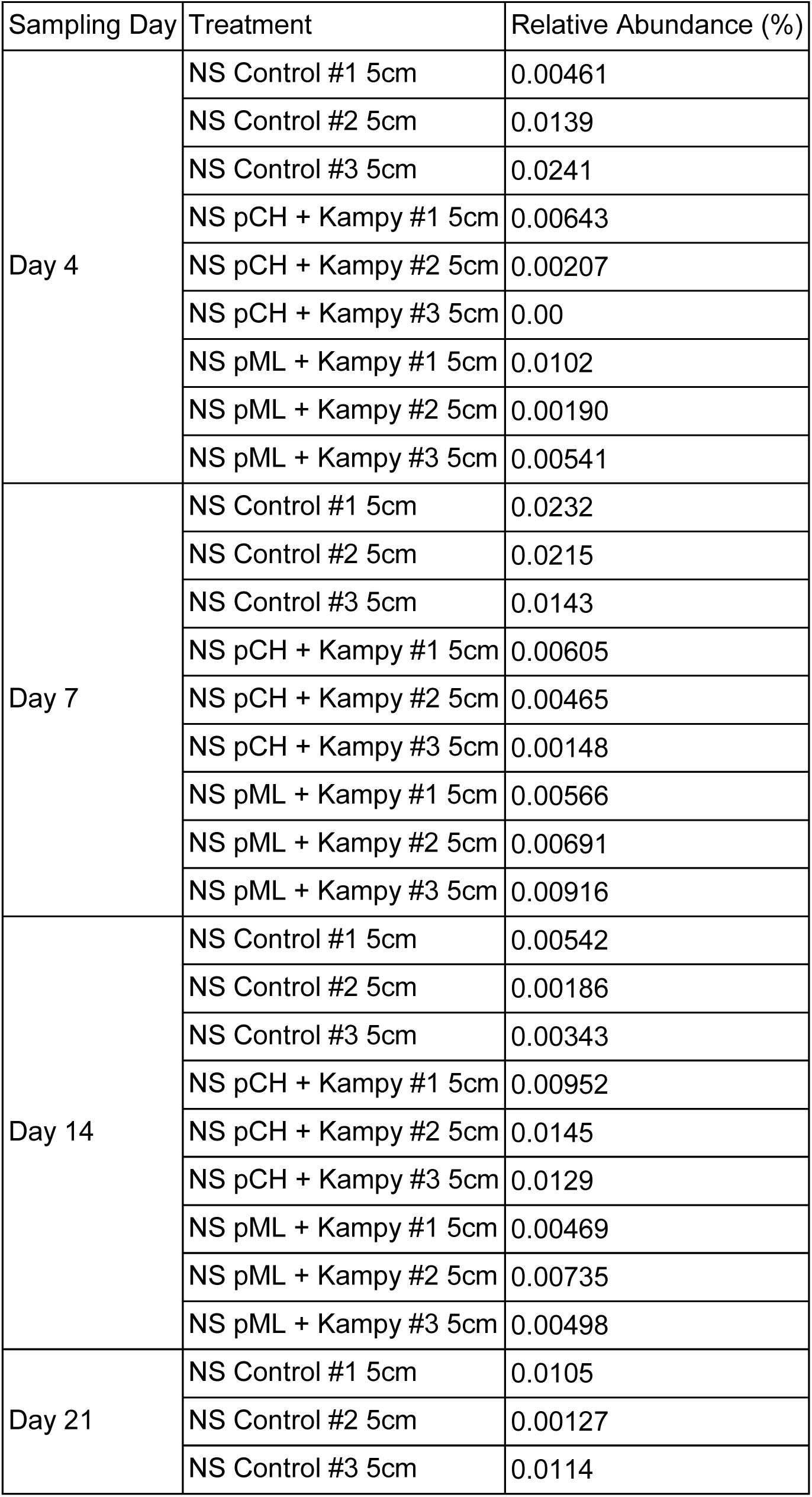
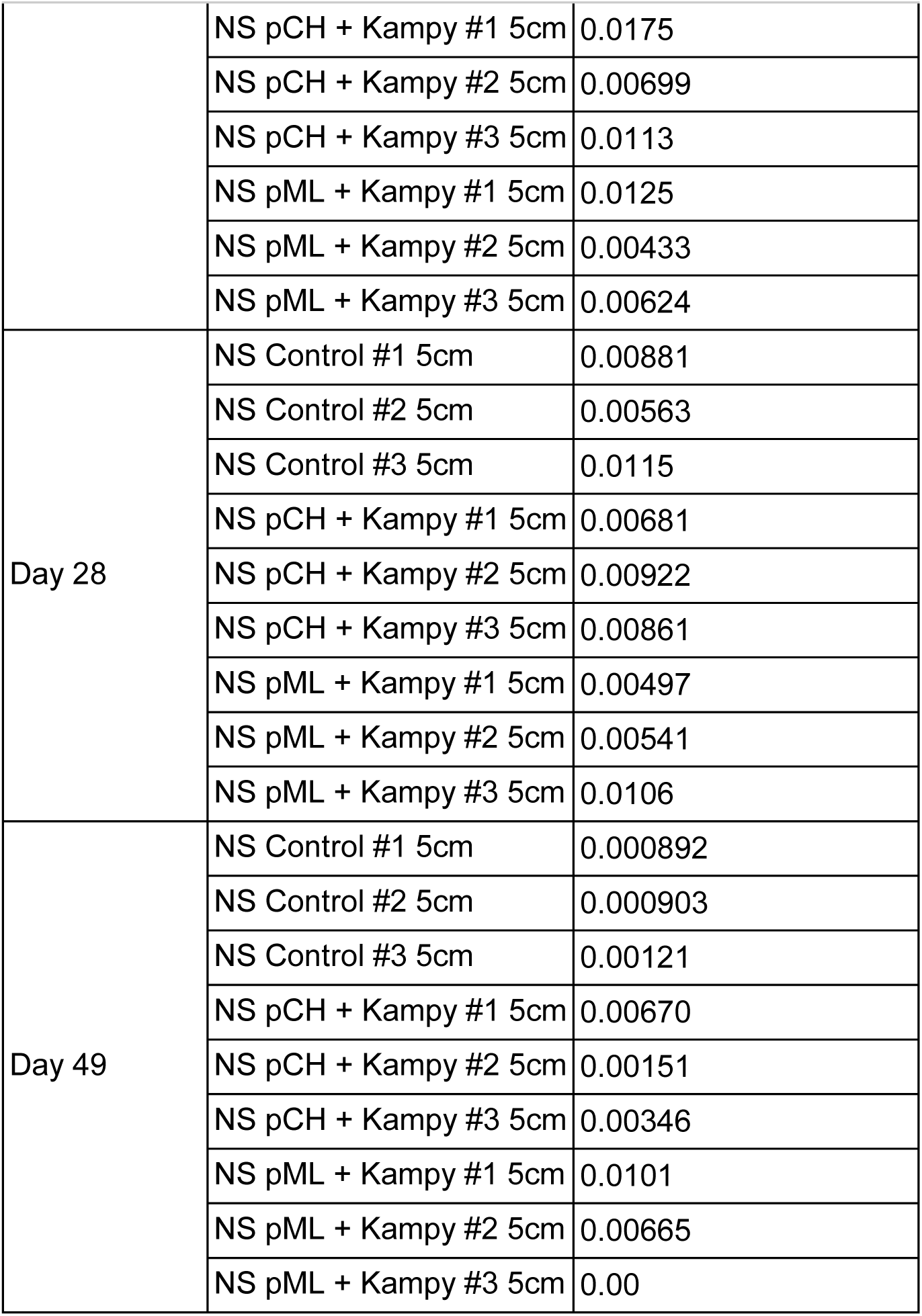
*M. smegmatis* 16s relative abundance at 5 cm away from the inoculation site.

| Sampling Day | Treatment | Relative Abundance (%) |
| --- | --- | --- |
| Day 4 | NS Control #1 5cm | 0.00461 |
|  | NS Control #2 5cm | 0.0139 |
|  | NS Control #3 5cm | 0.0241 |
|  | NS pCH + Kamy #1 5cm | 0.00643 |
|  | NS pCH + Kamy #2 5cm | 0.00207 |
|  | NS pCH + Kamy #3 5cm | 0.00 |
|  | NS pML + Kamy #1 5cm | 0.0102 |
|  | NS pML + Kamy #2 5cm | 0.00190 |
|  | NS pML + Kamy #3 5cm | 0.00541 |
| Day 7 | NS Control #1 5cm | 0.0232 |
|  | NS Control #2 5cm | 0.0215 |
|  | NS Control #3 5cm | 0.0143 |
|  | NS pCH + Kamy #1 5cm | 0.00605 |
|  | NS pCH + Kamy #2 5cm | 0.00465 |
|  | NS pCH + Kamy #3 5cm | 0.00148 |
|  | NS pML + Kamy #1 5cm | 0.00566 |
|  | NS pML + Kamy #2 5cm | 0.00691 |
|  | NS pML + Kamy #3 5cm | 0.00916 |
| Day 14 | NS Control #1 5cm | 0.00542 |
|  | NS Control #2 5cm | 0.00186 |
|  | NS Control #3 5cm | 0.00343 |
|  | NS pCH + Kamy #1 5cm | 0.00952 |
|  | NS pCH + Kamy #2 5cm | 0.0145 |
|  | NS pCH + Kamy #3 5cm | 0.0129 |
|  | NS pML + Kamy #1 5cm | 0.00469 |
|  | NS pML + Kamy #2 5cm | 0.00735 |
|  | NS pML + Kamy #3 5cm | 0.00498 |
| Day 21 | NS Control #1 5cm | 0.0105 |
|  | NS Control #2 5cm | 0.00127 |
|  | NS Control #3 5cm | 0.0114 |
|  | NS pCH + Kampy #1 5cm | 0.0175 |
|  | NS pCH + Kampy #2 5cm | 0.00699 |
|  | NS pCH + Kampy #3 5cm | 0.0113 |
|  | NS pML + Kampy #1 5cm | 0.0125 |
|  | NS pML + Kampy #2 5cm | 0.00433 |
|  | NS pML + Kampy #3 5cm | 0.00624 |
| Day 28 | NS Control #1 5cm | 0.00881 |
|  | NS Control #2 5cm | 0.00563 |
|  | NS Control #3 5cm | 0.0115 |
|  | NS pCH + Kampy #1 5cm | 0.00681 |
|  | NS pCH + Kampy #2 5cm | 0.00922 |
|  | NS pCH + Kampy #3 5cm | 0.00861 |
|  | NS pML + Kampy #1 5cm | 0.00497 |
|  | NS pML + Kampy #2 5cm | 0.00541 |
|  | NS pML + Kampy #3 5cm | 0.0106 |
| Day 49 | NS Control #1 5cm | 0.000892 |
|  | NS Control #2 5cm | 0.000903 |
|  | NS Control #3 5cm | 0.00121 |
|  | NS pCH + Kampy #1 5cm | 0.00670 |
|  | NS pCH + Kampy #2 5cm | 0.00151 |
|  | NS pCH + Kampy #3 5cm | 0.00346 |
|  | NS pML + Kampy #1 5cm | 0.0101 |
|  | NS pML + Kampy #2 5cm | 0.00665 |
|  | NS pML + Kampy #3 5cm | 0.00 |

**Table S5:**
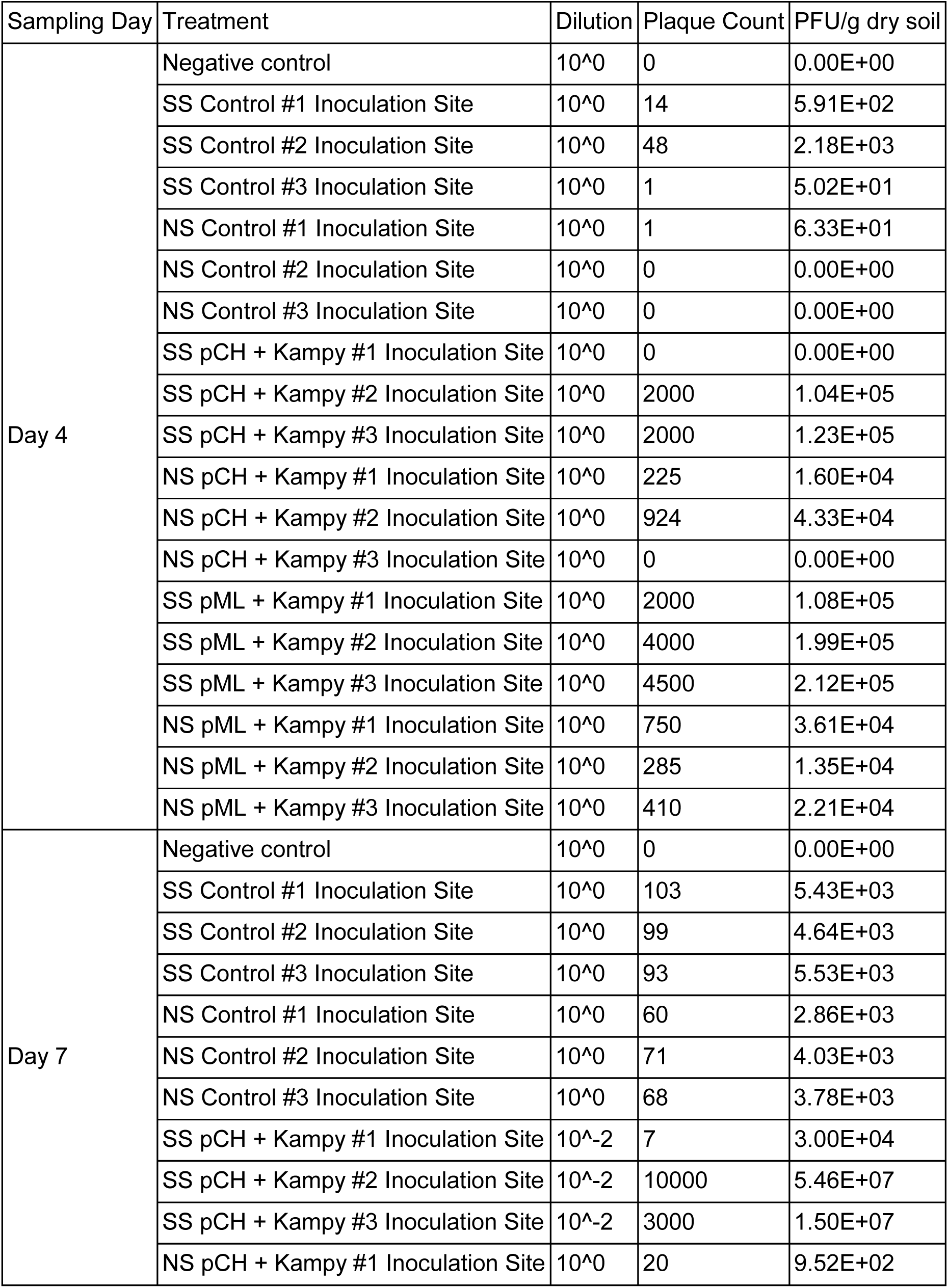
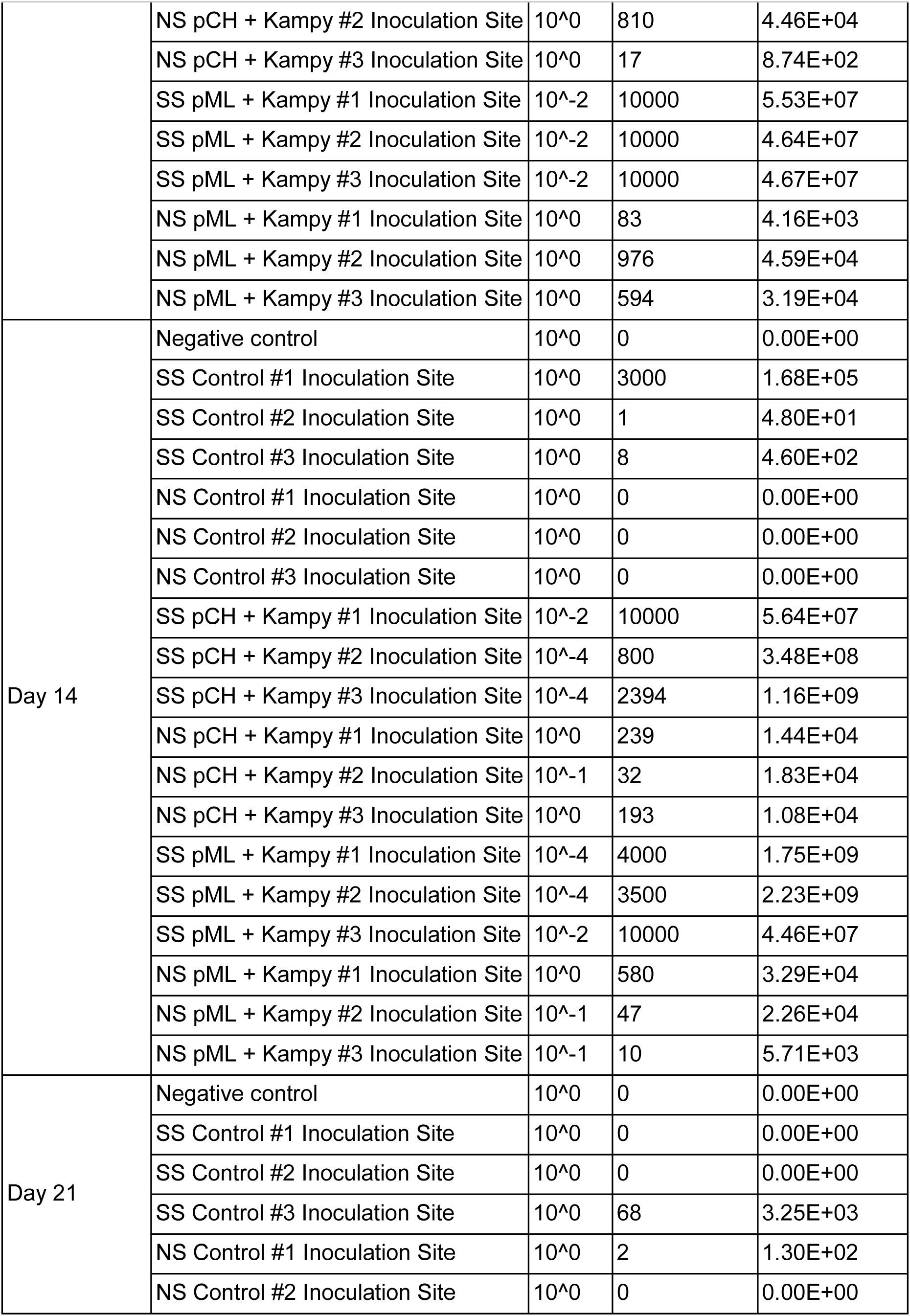
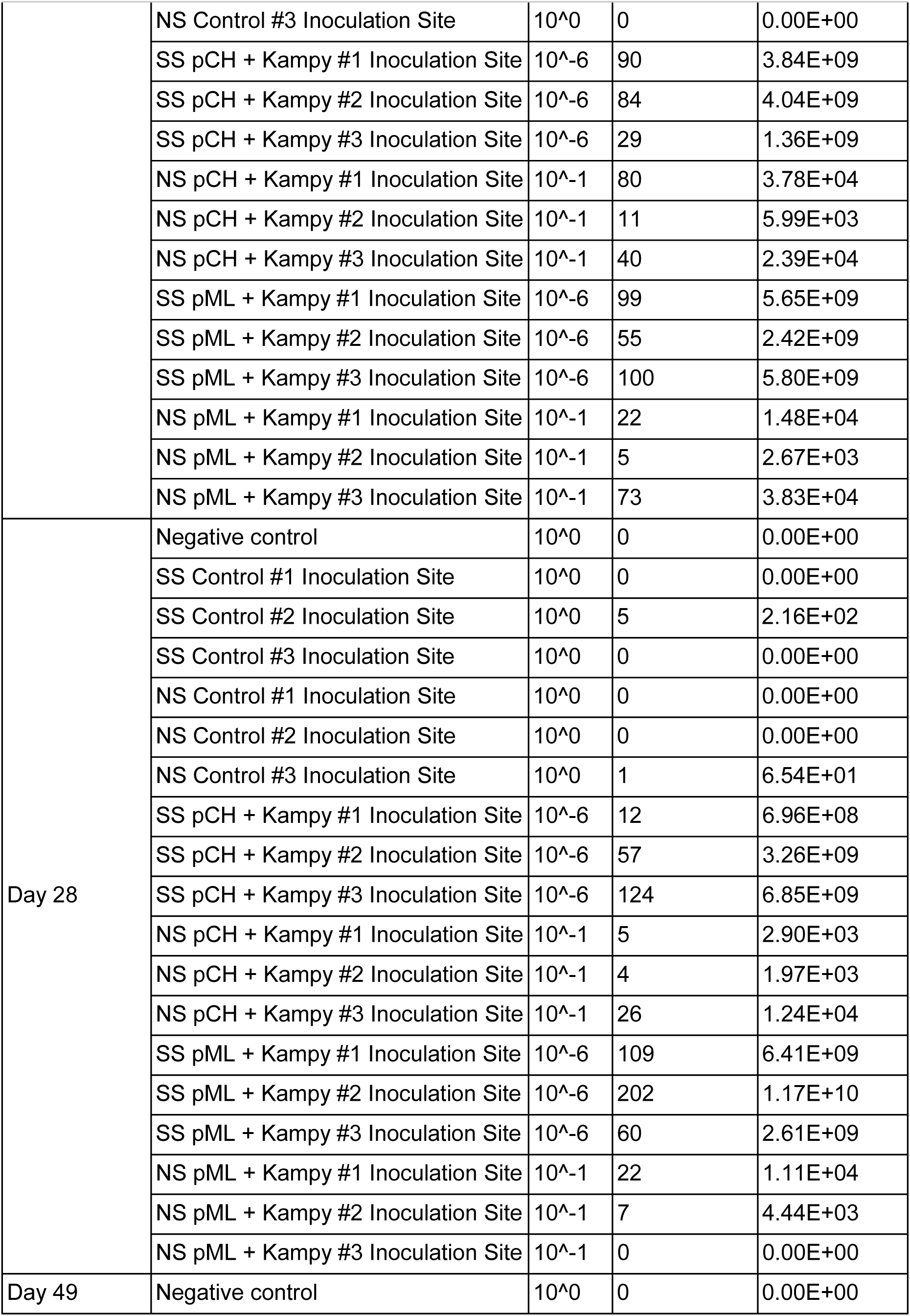
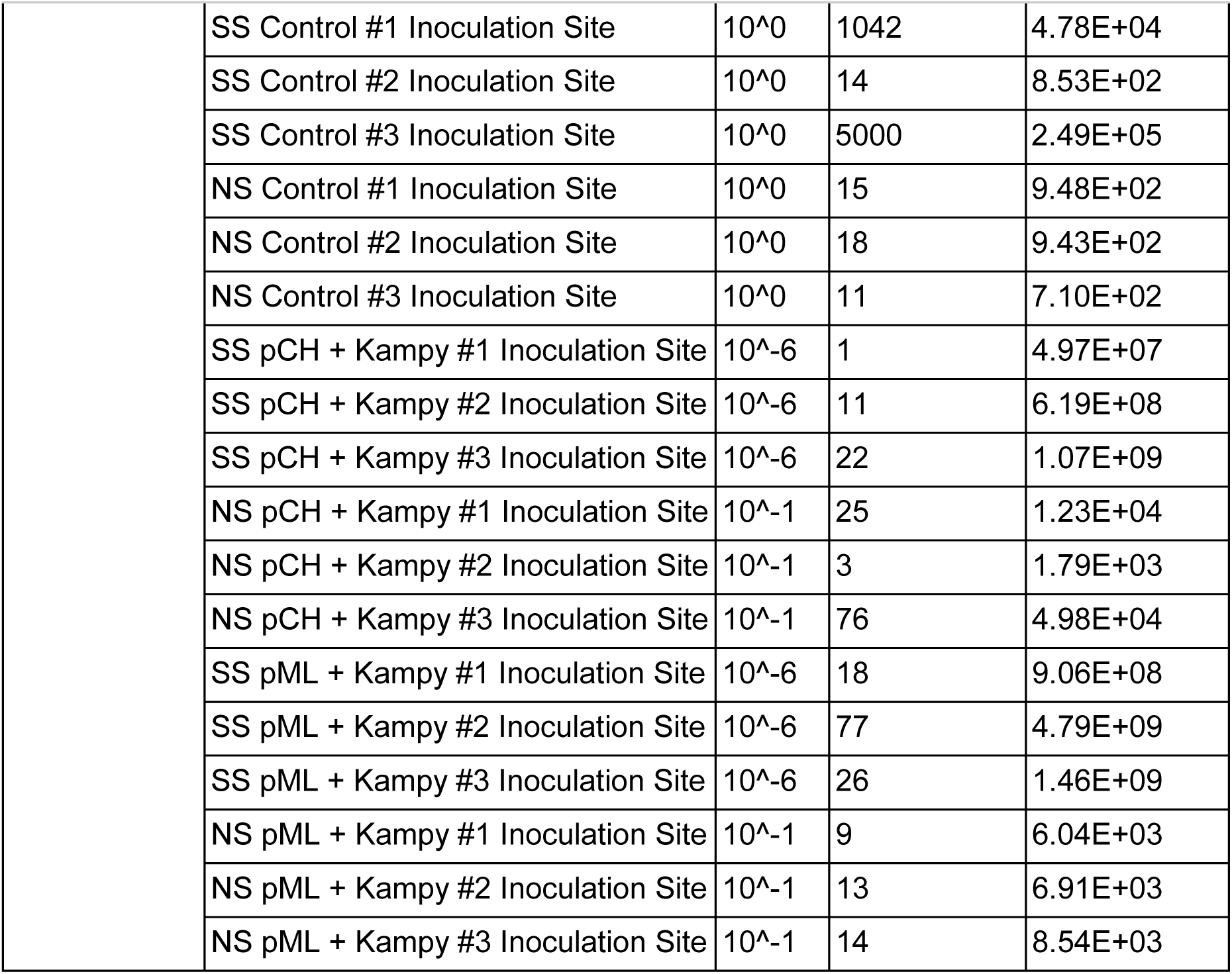
Kampy plaque counts and calculated PFU/g of dry soil at the inoculation site.

| Sampling Day | Treatment | Dilution | Plaque Count | PFU/g dry soil |
| --- | --- | --- | --- | --- |
| Day 4 | Negative control | 10<sup>0</sup> | 0 | 0.00E+00 |
|  | SS Control #1 Inoculation Site | 10<sup>0</sup> | 14 | 5.91E+02 |
|  | SS Control #2 Inoculation Site | 10<sup>0</sup> | 48 | 2.18E+03 |
|  | SS Control #3 Inoculation Site | 10<sup>0</sup> | 1 | 5.02E+01 |
|  | NS Control #1 Inoculation Site | 10<sup>0</sup> | 1 | 6.33E+01 |
|  | NS Control #2 Inoculation Site | 10<sup>0</sup> | 0 | 0.00E+00 |
|  | NS Control #3 Inoculation Site | 10<sup>0</sup> | 0 | 0.00E+00 |
|  | SS pCH + Kamy #1 Inoculation Site | 10<sup>0</sup> | 0 | 0.00E+00 |
|  | SS pCH + Kamy #2 Inoculation Site | 10<sup>0</sup> | 2000 | 1.04E+05 |
|  | SS pCH + Kamy #3 Inoculation Site | 10<sup>0</sup> | 2000 | 1.23E+05 |
|  | NS pCH + Kamy #1 Inoculation Site | 10<sup>0</sup> | 225 | 1.60E+04 |
|  | NS pCH + Kamy #2 Inoculation Site | 10<sup>0</sup> | 924 | 4.33E+04 |
|  | NS pCH + Kamy #3 Inoculation Site | 10<sup>0</sup> | 0 | 0.00E+00 |
|  | SS pML + Kamy #1 Inoculation Site | 10<sup>0</sup> | 2000 | 1.08E+05 |
|  | SS pML + Kamy #2 Inoculation Site | 10<sup>0</sup> | 4000 | 1.99E+05 |
|  | SS pML + Kamy #3 Inoculation Site | 10<sup>0</sup> | 4500 | 2.12E+05 |
|  | NS pML + Kamy #1 Inoculation Site | 10<sup>0</sup> | 750 | 3.61E+04 |
|  | NS pML + Kamy #2 Inoculation Site | 10<sup>0</sup> | 285 | 1.35E+04 |
|  | NS pML + Kamy #3 Inoculation Site | 10<sup>0</sup> | 410 | 2.21E+04 |
| Day 7 | Negative control | 10<sup>0</sup> | 0 | 0.00E+00 |
|  | SS Control #1 Inoculation Site | 10<sup>0</sup> | 103 | 5.43E+03 |
|  | SS Control #2 Inoculation Site | 10<sup>0</sup> | 99 | 4.64E+03 |
|  | SS Control #3 Inoculation Site | 10<sup>0</sup> | 93 | 5.53E+03 |
|  | NS Control #1 Inoculation Site | 10<sup>0</sup> | 60 | 2.86E+03 |
|  | NS Control #2 Inoculation Site | 10<sup>0</sup> | 71 | 4.03E+03 |
|  | NS Control #3 Inoculation Site | 10<sup>0</sup> | 68 | 3.78E+03 |
|  | SS pCH + Kamy #1 Inoculation Site | 10<sup>-2</sup> | 7 | 3.00E+04 |
|  | SS pCH + Kamy #2 Inoculation Site | 10<sup>-2</sup> | 10000 | 5.46E+07 |
|  | SS pCH + Kamy #3 Inoculation Site | 10<sup>-2</sup> | 3000 | 1.50E+07 |
|  | NS pCH + Kamy #1 Inoculation Site | 10<sup>0</sup> | 20 | 9.52E+02 |
|  | NS pCH + Kamy #2 Inoculation Site | 10<sup>0</sup> | 810 | 4.46E+04 |
|  | NS pCH + Kamy #3 Inoculation Site | 10<sup>0</sup> | 17 | 8.74E+02 |
|  | SS pML + Kamy #1 Inoculation Site | 10<sup>-2</sup> | 10000 | 5.53E+07 |
|  | SS pML + Kamy #2 Inoculation Site | 10<sup>-2</sup> | 10000 | 4.64E+07 |
|  | SS pML + Kamy #3 Inoculation Site | 10<sup>-2</sup> | 10000 | 4.67E+07 |
|  | NS pML + Kamy #1 Inoculation Site | 10<sup>0</sup> | 83 | 4.16E+03 |
|  | NS pML + Kamy #2 Inoculation Site | 10<sup>0</sup> | 976 | 4.59E+04 |
|  | NS pML + Kamy #3 Inoculation Site | 10<sup>0</sup> | 594 | 3.19E+04 |
| Day 14 | Negative control | 10<sup>0</sup> | 0 | 0.00E+00 |
|  | SS Control #1 Inoculation Site | 10<sup>0</sup> | 3000 | 1.68E+05 |
|  | SS Control #2 Inoculation Site | 10<sup>0</sup> | 1 | 4.80E+01 |
|  | SS Control #3 Inoculation Site | 10<sup>0</sup> | 8 | 4.60E+02 |
|  | NS Control #1 Inoculation Site | 10<sup>0</sup> | 0 | 0.00E+00 |
|  | NS Control #2 Inoculation Site | 10<sup>0</sup> | 0 | 0.00E+00 |
|  | NS Control #3 Inoculation Site | 10<sup>0</sup> | 0 | 0.00E+00 |
|  | SS pCH + Kamy #1 Inoculation Site | 10<sup>-2</sup> | 10000 | 5.64E+07 |
|  | SS pCH + Kamy #2 Inoculation Site | 10<sup>-4</sup> | 800 | 3.48E+08 |
|  | SS pCH + Kamy #3 Inoculation Site | 10<sup>-4</sup> | 2394 | 1.16E+09 |
|  | NS pCH + Kamy #1 Inoculation Site | 10<sup>0</sup> | 239 | 1.44E+04 |
|  | NS pCH + Kamy #2 Inoculation Site | 10<sup>-1</sup> | 32 | 1.83E+04 |
|  | NS pCH + Kamy #3 Inoculation Site | 10<sup>0</sup> | 193 | 1.08E+04 |
|  | SS pML + Kamy #1 Inoculation Site | 10<sup>-4</sup> | 4000 | 1.75E+09 |
|  | SS pML + Kamy #2 Inoculation Site | 10<sup>-4</sup> | 3500 | 2.23E+09 |
|  | SS pML + Kamy #3 Inoculation Site | 10<sup>-2</sup> | 10000 | 4.46E+07 |
|  | NS pML + Kamy #1 Inoculation Site | 10<sup>0</sup> | 580 | 3.29E+04 |
|  | NS pML + Kamy #2 Inoculation Site | 10<sup>-1</sup> | 47 | 2.26E+04 |
|  | NS pML + Kamy #3 Inoculation Site | 10<sup>-1</sup> | 10 | 5.71E+03 |
| Day 21 | Negative control | 10<sup>0</sup> | 0 | 0.00E+00 |
|  | SS Control #1 Inoculation Site | 10<sup>0</sup> | 0 | 0.00E+00 |
|  | SS Control #2 Inoculation Site | 10<sup>0</sup> | 0 | 0.00E+00 |
|  | SS Control #3 Inoculation Site | 10<sup>0</sup> | 68 | 3.25E+03 |
|  | NS Control #1 Inoculation Site | 10<sup>0</sup> | 2 | 1.30E+02 |
|  | NS Control #2 Inoculation Site | 10<sup>0</sup> | 0 | 0.00E+00 |
|  | NS Control #3 Inoculation Site | 10<sup>0</sup> | 0 | 0.00E+00 |
|  | SS pCH + Kamy #1 Inoculation Site | 10<sup>-6</sup> | 90 | 3.84E+09 |
|  | SS pCH + Kamy #2 Inoculation Site | 10<sup>-6</sup> | 84 | 4.04E+09 |
|  | SS pCH + Kamy #3 Inoculation Site | 10<sup>-6</sup> | 29 | 1.36E+09 |
|  | NS pCH + Kamy #1 Inoculation Site | 10<sup>-1</sup> | 80 | 3.78E+04 |
|  | NS pCH + Kamy #2 Inoculation Site | 10<sup>-1</sup> | 11 | 5.99E+03 |
|  | NS pCH + Kamy #3 Inoculation Site | 10<sup>-1</sup> | 40 | 2.39E+04 |
|  | SS pML + Kamy #1 Inoculation Site | 10<sup>-6</sup> | 99 | 5.65E+09 |
|  | SS pML + Kamy #2 Inoculation Site | 10<sup>-6</sup> | 55 | 2.42E+09 |
|  | SS pML + Kamy #3 Inoculation Site | 10<sup>-6</sup> | 100 | 5.80E+09 |
|  | NS pML + Kamy #1 Inoculation Site | 10<sup>-1</sup> | 22 | 1.48E+04 |
|  | NS pML + Kamy #2 Inoculation Site | 10<sup>-1</sup> | 5 | 2.67E+03 |
|  | NS pML + Kamy #3 Inoculation Site | 10<sup>-1</sup> | 73 | 3.83E+04 |
| Day 28 | Negative control | 10<sup>0</sup> | 0 | 0.00E+00 |
|  | SS Control #1 Inoculation Site | 10<sup>0</sup> | 0 | 0.00E+00 |
|  | SS Control #2 Inoculation Site | 10<sup>0</sup> | 5 | 2.16E+02 |
|  | SS Control #3 Inoculation Site | 10<sup>0</sup> | 0 | 0.00E+00 |
|  | NS Control #1 Inoculation Site | 10<sup>0</sup> | 0 | 0.00E+00 |
|  | NS Control #2 Inoculation Site | 10<sup>0</sup> | 0 | 0.00E+00 |
|  | NS Control #3 Inoculation Site | 10<sup>0</sup> | 1 | 6.54E+01 |
|  | SS pCH + Kamy #1 Inoculation Site | 10<sup>-6</sup> | 12 | 6.96E+08 |
|  | SS pCH + Kamy #2 Inoculation Site | 10<sup>-6</sup> | 57 | 3.26E+09 |
|  | SS pCH + Kamy #3 Inoculation Site | 10<sup>-6</sup> | 124 | 6.85E+09 |
|  | NS pCH + Kamy #1 Inoculation Site | 10<sup>-1</sup> | 5 | 2.90E+03 |
|  | NS pCH + Kamy #2 Inoculation Site | 10<sup>-1</sup> | 4 | 1.97E+03 |
|  | NS pCH + Kamy #3 Inoculation Site | 10<sup>-1</sup> | 26 | 1.24E+04 |
|  | SS pML + Kamy #1 Inoculation Site | 10<sup>-6</sup> | 109 | 6.41E+09 |
|  | SS pML + Kamy #2 Inoculation Site | 10<sup>-6</sup> | 202 | 1.17E+10 |
|  | SS pML + Kamy #3 Inoculation Site | 10<sup>-6</sup> | 60 | 2.61E+09 |
|  | NS pML + Kamy #1 Inoculation Site | 10<sup>-1</sup> | 22 | 1.11E+04 |
|  | NS pML + Kamy #2 Inoculation Site | 10<sup>-1</sup> | 7 | 4.44E+03 |
|  | NS pML + Kamy #3 Inoculation Site | 10<sup>-1</sup> | 0 | 0.00E+00 |
| Day 49 | Negative control | 10<sup>0</sup> | 0 | 0.00E+00 |

|  |  |  |  |
| --- | --- | --- | --- |
| SS Control #1 Inoculation Site | 10<sup>0</sup> | 1042 | 4.78E+04 |
| SS Control #2 Inoculation Site | 10<sup>0</sup> | 14 | 8.53E+02 |
| SS Control #3 Inoculation Site | 10<sup>0</sup> | 5000 | 2.49E+05 |
| NS Control #1 Inoculation Site | 10<sup>0</sup> | 15 | 9.48E+02 |
| NS Control #2 Inoculation Site | 10<sup>0</sup> | 18 | 9.43E+02 |
| NS Control #3 Inoculation Site | 10<sup>0</sup> | 11 | 7.10E+02 |
| SS pCH + Kamy #1 Inoculation Site | 10<sup>-6</sup> | 1 | 4.97E+07 |
| SS pCH + Kamy #2 Inoculation Site | 10<sup>-6</sup> | 11 | 6.19E+08 |
| SS pCH + Kamy #3 Inoculation Site | 10<sup>-6</sup> | 22 | 1.07E+09 |
| NS pCH + Kamy #1 Inoculation Site | 10<sup>-1</sup> | 25 | 1.23E+04 |
| NS pCH + Kamy #2 Inoculation Site | 10<sup>-1</sup> | 3 | 1.79E+03 |
| NS pCH + Kamy #3 Inoculation Site | 10<sup>-1</sup> | 76 | 4.98E+04 |
| SS pML + Kamy #1 Inoculation Site | 10<sup>-6</sup> | 18 | 9.06E+08 |
| SS pML + Kamy #2 Inoculation Site | 10<sup>-6</sup> | 77 | 4.79E+09 |
| SS pML + Kamy #3 Inoculation Site | 10<sup>-6</sup> | 26 | 1.46E+09 |
| NS pML + Kamy #1 Inoculation Site | 10<sup>-1</sup> | 9 | 6.04E+03 |
| NS pML + Kamy #2 Inoculation Site | 10<sup>-1</sup> | 13 | 6.91E+03 |
| NS pML + Kamy #3 Inoculation Site | 10<sup>-1</sup> | 14 | 8.54E+03 |

**Table S6:** Kampy metagenomic relative abundance at the inoculation site.

| Sampling Day | Treatment | Relative Abundance (%) |
| --- | --- | --- |
| 7 | NS pCH + Kamy #1 Inoculation Site | 0.0000198 |
|  | NS pCH + Kamy #2 Inoculation Site | 0.000252 |
|  | NS pCH + Kamy #3 Inoculation Site | 0.0000369 |
|  | NS pML + Kamy #1 Inoculation Site | 0.00 |
|  | NS pML + Kamy #2 Inoculation Site | 0.000157 |
|  | NS pML + Kamy #3 Inoculation Site | 0.0000697 |
| 49 | NS pCH + Kamy #1 Inoculation Site | 0.000161 |
|  | NS pCH + Kamy #2 Inoculation Site | 0.0000132 |
|  | NS pCH + Kamy #3 Inoculation Site | 0.0000835 |
|  | NS pML + Kamy #1 Inoculation Site | 0.00000309 |
|  | NS pML + Kamy #2 Inoculation Site | 0.0000148 |
|  | NS pML + Kamy #3 Inoculation Site | 0.000105 |

**Table S7:**
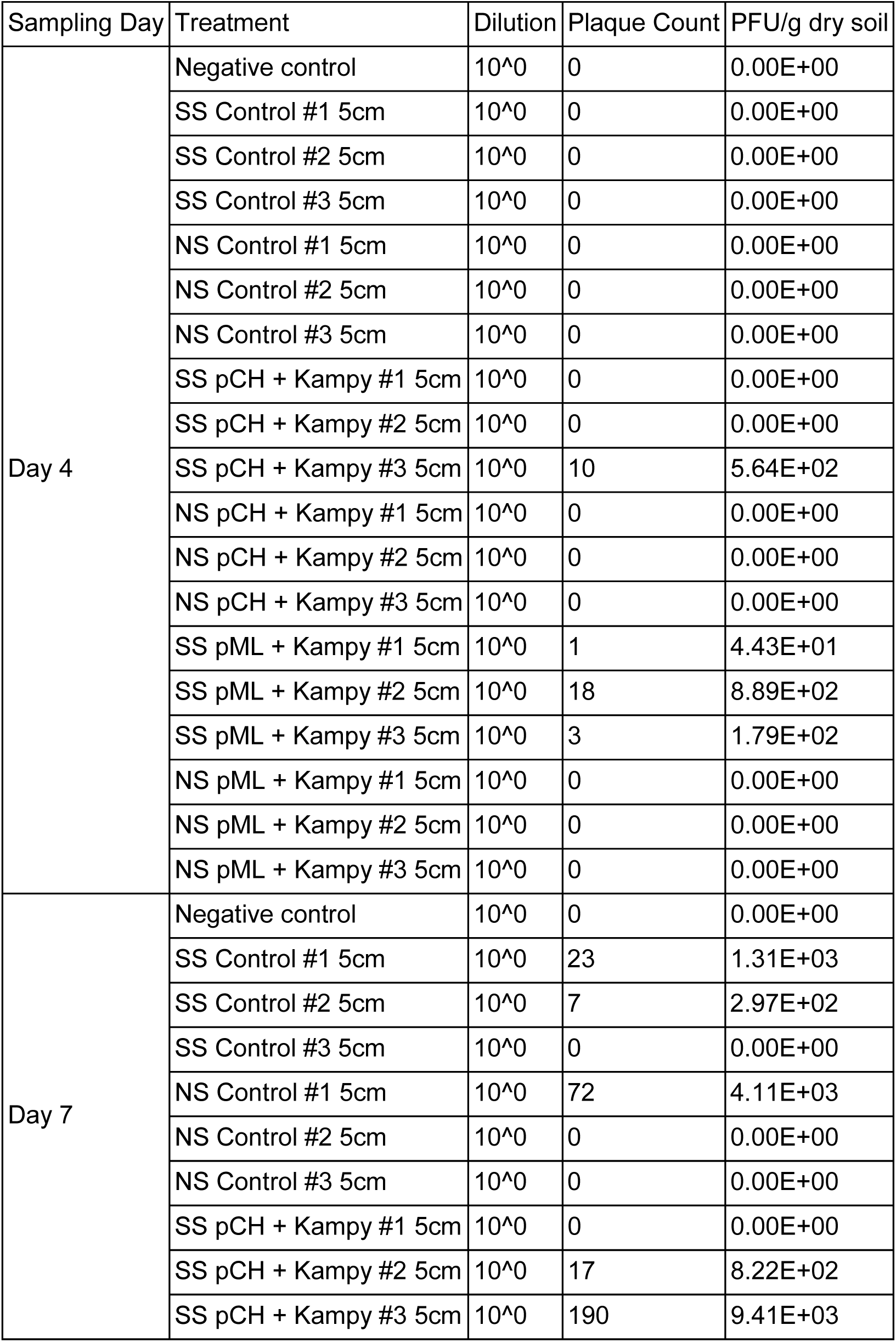
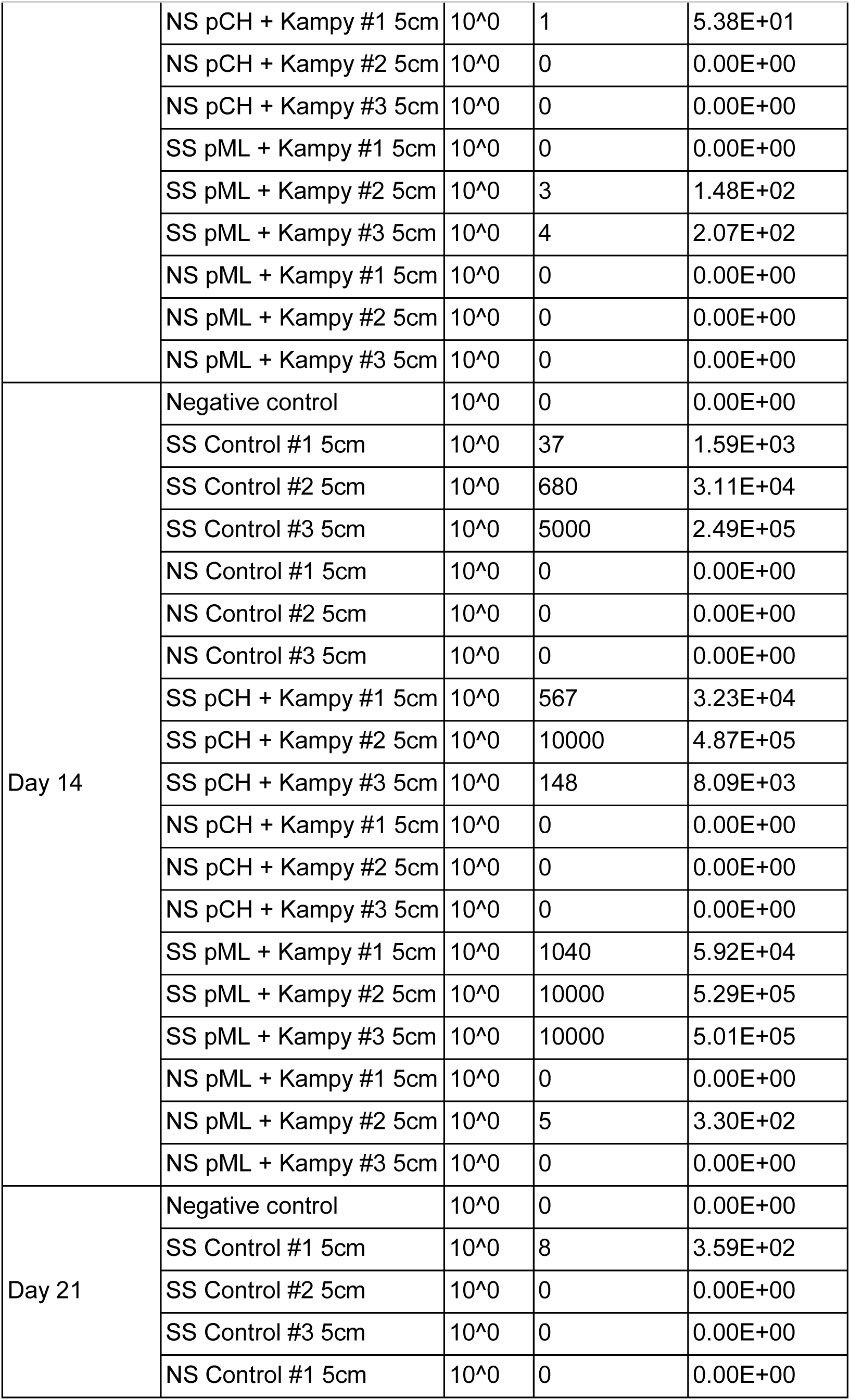
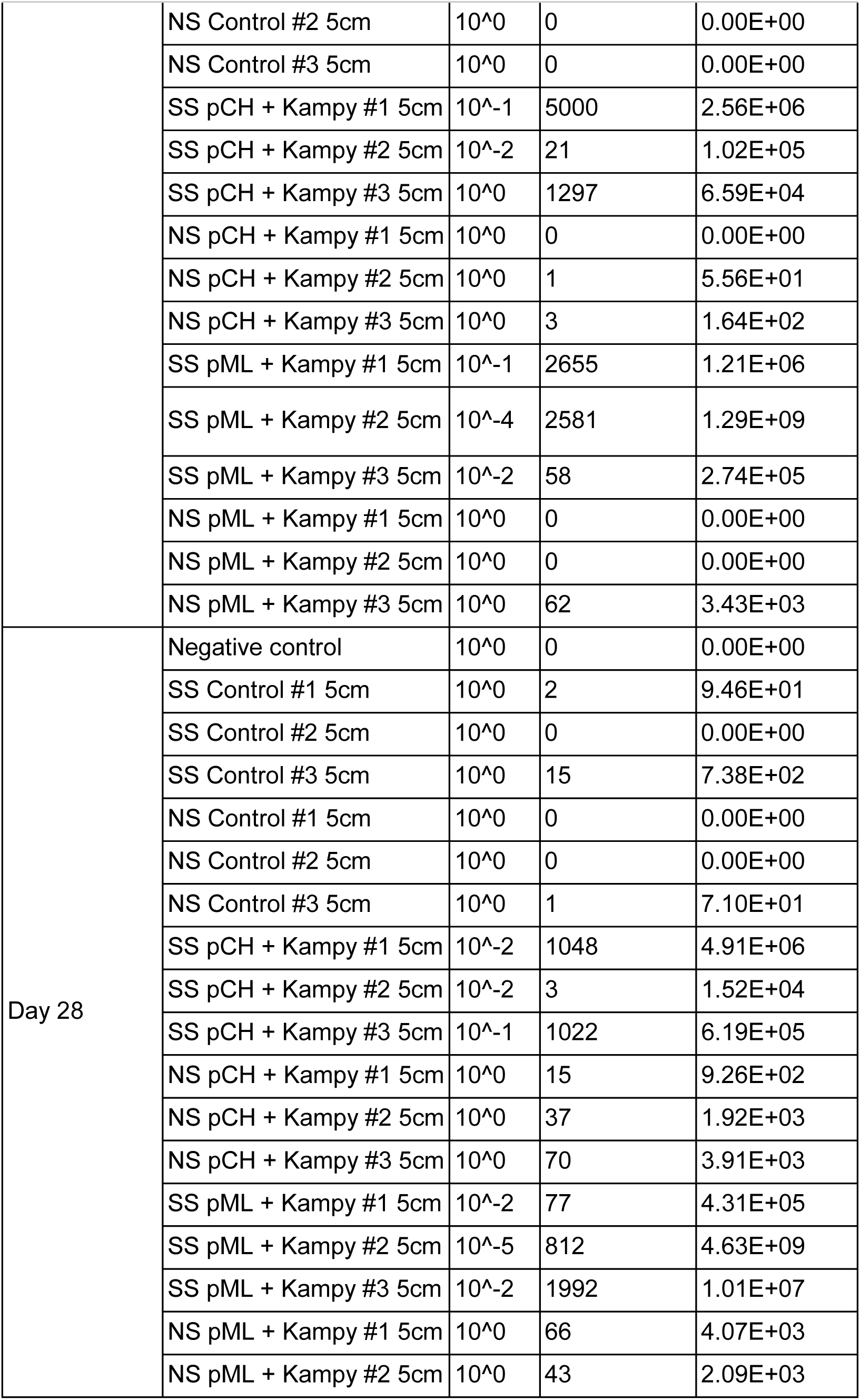
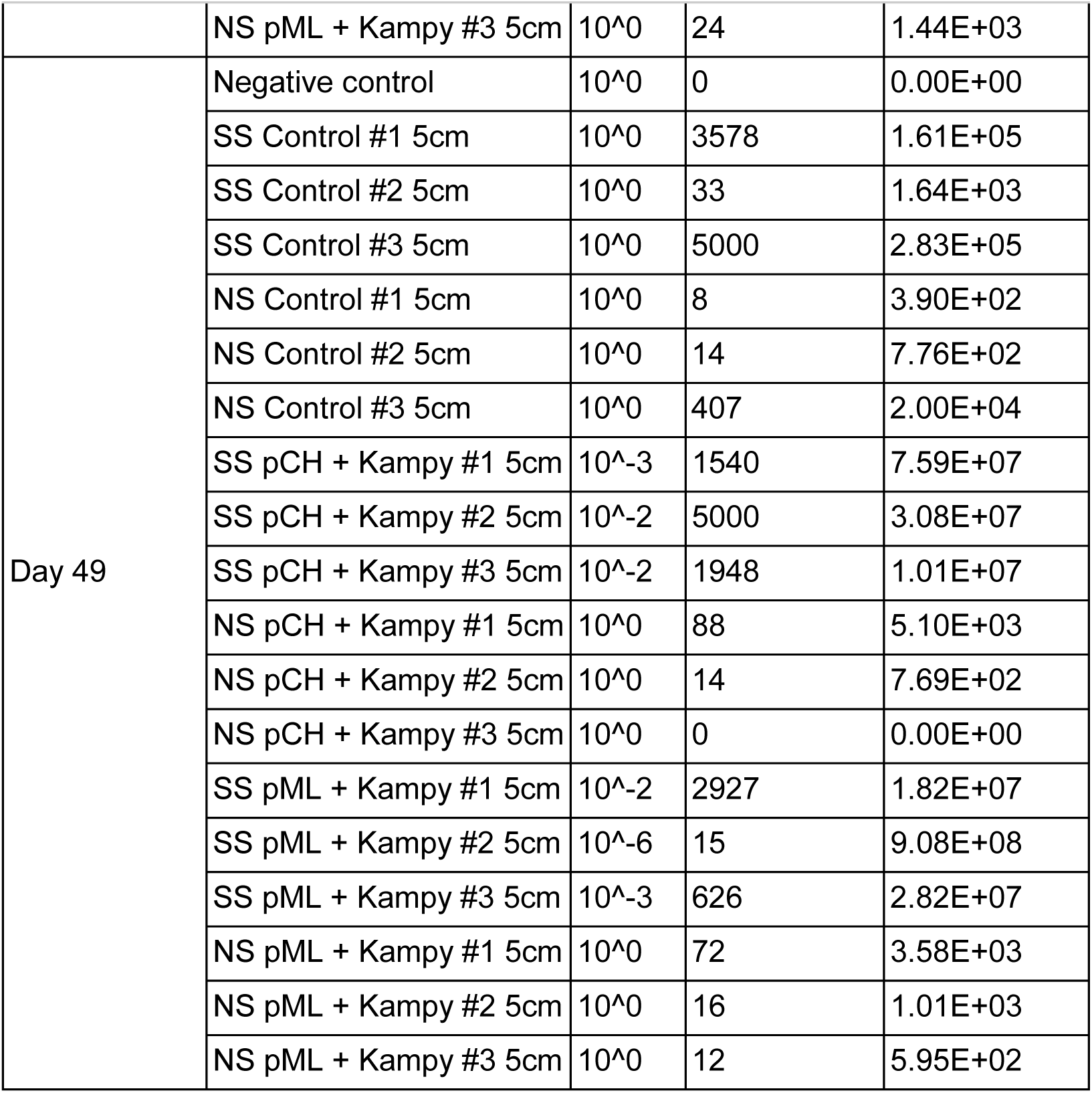
Kampy plaque counts and calculated PFU/g of dry soil at 5 cm away from the inoculation site.

| Sampling Day | Treatment | Dilution | Plaque Count | PFU/g dry soil |
| --- | --- | --- | --- | --- |
| Day 4 | Negative control | 10<sup>0</sup> | 0 | 0.00E+00 |
|  | SS Control #1 5cm | 10<sup>0</sup> | 0 | 0.00E+00 |
|  | SS Control #2 5cm | 10<sup>0</sup> | 0 | 0.00E+00 |
|  | SS Control #3 5cm | 10<sup>0</sup> | 0 | 0.00E+00 |
|  | NS Control #1 5cm | 10<sup>0</sup> | 0 | 0.00E+00 |
|  | NS Control #2 5cm | 10<sup>0</sup> | 0 | 0.00E+00 |
|  | NS Control #3 5cm | 10<sup>0</sup> | 0 | 0.00E+00 |
|  | SS pCH + Kampy #1 5cm | 10<sup>0</sup> | 0 | 0.00E+00 |
|  | SS pCH + Kampy #2 5cm | 10<sup>0</sup> | 0 | 0.00E+00 |
|  | SS pCH + Kampy #3 5cm | 10<sup>0</sup> | 10 | 5.64E+02 |
|  | NS pCH + Kampy #1 5cm | 10<sup>0</sup> | 0 | 0.00E+00 |
|  | NS pCH + Kampy #2 5cm | 10<sup>0</sup> | 0 | 0.00E+00 |
|  | NS pCH + Kampy #3 5cm | 10<sup>0</sup> | 0 | 0.00E+00 |
|  | SS pML + Kampy #1 5cm | 10<sup>0</sup> | 1 | 4.43E+01 |
|  | SS pML + Kampy #2 5cm | 10<sup>0</sup> | 18 | 8.89E+02 |
|  | SS pML + Kampy #3 5cm | 10<sup>0</sup> | 3 | 1.79E+02 |
|  | NS pML + Kampy #1 5cm | 10<sup>0</sup> | 0 | 0.00E+00 |
|  | NS pML + Kampy #2 5cm | 10<sup>0</sup> | 0 | 0.00E+00 |
|  | NS pML + Kampy #3 5cm | 10<sup>0</sup> | 0 | 0.00E+00 |
| Day 7 | Negative control | 10<sup>0</sup> | 0 | 0.00E+00 |
|  | SS Control #1 5cm | 10<sup>0</sup> | 23 | 1.31E+03 |
|  | SS Control #2 5cm | 10<sup>0</sup> | 7 | 2.97E+02 |
|  | SS Control #3 5cm | 10<sup>0</sup> | 0 | 0.00E+00 |
|  | NS Control #1 5cm | 10<sup>0</sup> | 72 | 4.11E+03 |
|  | NS Control #2 5cm | 10<sup>0</sup> | 0 | 0.00E+00 |
|  | NS Control #3 5cm | 10<sup>0</sup> | 0 | 0.00E+00 |
|  | SS pCH + Kampy #1 5cm | 10<sup>0</sup> | 0 | 0.00E+00 |
|  | SS pCH + Kampy #2 5cm | 10<sup>0</sup> | 17 | 8.22E+02 |
|  | SS pCH + Kampy #3 5cm | 10<sup>0</sup> | 190 | 9.41E+03 |
|  | NS pCH + Kampy #1 5cm | 10<sup>0</sup> | 1 | 5.38E+01 |
|  | NS pCH + Kampy #2 5cm | 10<sup>0</sup> | 0 | 0.00E+00 |
|  | NS pCH + Kampy #3 5cm | 10<sup>0</sup> | 0 | 0.00E+00 |
|  | SS pML + Kampy #1 5cm | 10<sup>0</sup> | 0 | 0.00E+00 |
|  | SS pML + Kampy #2 5cm | 10<sup>0</sup> | 3 | 1.48E+02 |
|  | SS pML + Kampy #3 5cm | 10<sup>0</sup> | 4 | 2.07E+02 |
|  | NS pML + Kampy #1 5cm | 10<sup>0</sup> | 0 | 0.00E+00 |
|  | NS pML + Kampy #2 5cm | 10<sup>0</sup> | 0 | 0.00E+00 |
|  | NS pML + Kampy #3 5cm | 10<sup>0</sup> | 0 | 0.00E+00 |
| Day 14 | Negative control | 10<sup>0</sup> | 0 | 0.00E+00 |
|  | SS Control #1 5cm | 10<sup>0</sup> | 37 | 1.59E+03 |
|  | SS Control #2 5cm | 10<sup>0</sup> | 680 | 3.11E+04 |
|  | SS Control #3 5cm | 10<sup>0</sup> | 5000 | 2.49E+05 |
|  | NS Control #1 5cm | 10<sup>0</sup> | 0 | 0.00E+00 |
|  | NS Control #2 5cm | 10<sup>0</sup> | 0 | 0.00E+00 |
|  | NS Control #3 5cm | 10<sup>0</sup> | 0 | 0.00E+00 |
|  | SS pCH + Kampy #1 5cm | 10<sup>0</sup> | 567 | 3.23E+04 |
|  | SS pCH + Kampy #2 5cm | 10<sup>0</sup> | 10000 | 4.87E+05 |
|  | SS pCH + Kampy #3 5cm | 10<sup>0</sup> | 148 | 8.09E+03 |
|  | NS pCH + Kampy #1 5cm | 10<sup>0</sup> | 0 | 0.00E+00 |
|  | NS pCH + Kampy #2 5cm | 10<sup>0</sup> | 0 | 0.00E+00 |
|  | NS pCH + Kampy #3 5cm | 10<sup>0</sup> | 0 | 0.00E+00 |
|  | SS pML + Kampy #1 5cm | 10<sup>0</sup> | 1040 | 5.92E+04 |
|  | SS pML + Kampy #2 5cm | 10<sup>0</sup> | 10000 | 5.29E+05 |
|  | SS pML + Kampy #3 5cm | 10<sup>0</sup> | 10000 | 5.01E+05 |
|  | NS pML + Kampy #1 5cm | 10<sup>0</sup> | 0 | 0.00E+00 |
|  | NS pML + Kampy #2 5cm | 10<sup>0</sup> | 5 | 3.30E+02 |
|  | NS pML + Kampy #3 5cm | 10<sup>0</sup> | 0 | 0.00E+00 |
| Day 21 | Negative control | 10<sup>0</sup> | 0 | 0.00E+00 |
|  | SS Control #1 5cm | 10<sup>0</sup> | 8 | 3.59E+02 |
|  | SS Control #2 5cm | 10<sup>0</sup> | 0 | 0.00E+00 |
|  | SS Control #3 5cm | 10<sup>0</sup> | 0 | 0.00E+00 |
|  | NS Control #1 5cm | 10<sup>0</sup> | 0 | 0.00E+00 |
|  | NS Control #2 5cm | 10<sup>0</sup> | 0 | 0.00E+00 |
|  | NS Control #3 5cm | 10<sup>0</sup> | 0 | 0.00E+00 |
|  | SS pCH + Kampy #1 5cm | 10<sup>-1</sup> | 5000 | 2.56E+06 |
|  | SS pCH + Kampy #2 5cm | 10<sup>-2</sup> | 21 | 1.02E+05 |
|  | SS pCH + Kampy #3 5cm | 10<sup>0</sup> | 1297 | 6.59E+04 |
|  | NS pCH + Kampy #1 5cm | 10<sup>0</sup> | 0 | 0.00E+00 |
|  | NS pCH + Kampy #2 5cm | 10<sup>0</sup> | 1 | 5.56E+01 |
|  | NS pCH + Kampy #3 5cm | 10<sup>0</sup> | 3 | 1.64E+02 |
|  | SS pML + Kampy #1 5cm | 10<sup>-1</sup> | 2655 | 1.21E+06 |
|  | SS pML + Kampy #2 5cm | 10<sup>-4</sup> | 2581 | 1.29E+09 |
|  | SS pML + Kampy #3 5cm | 10<sup>-2</sup> | 58 | 2.74E+05 |
|  | NS pML + Kampy #1 5cm | 10<sup>0</sup> | 0 | 0.00E+00 |
|  | NS pML + Kampy #2 5cm | 10<sup>0</sup> | 0 | 0.00E+00 |
|  | NS pML + Kampy #3 5cm | 10<sup>0</sup> | 62 | 3.43E+03 |
| Day 28 | Negative control | 10<sup>0</sup> | 0 | 0.00E+00 |
|  | SS Control #1 5cm | 10<sup>0</sup> | 2 | 9.46E+01 |
|  | SS Control #2 5cm | 10<sup>0</sup> | 0 | 0.00E+00 |
|  | SS Control #3 5cm | 10<sup>0</sup> | 15 | 7.38E+02 |
|  | NS Control #1 5cm | 10<sup>0</sup> | 0 | 0.00E+00 |
|  | NS Control #2 5cm | 10<sup>0</sup> | 0 | 0.00E+00 |
|  | NS Control #3 5cm | 10<sup>0</sup> | 1 | 7.10E+01 |
|  | SS pCH + Kampy #1 5cm | 10<sup>-2</sup> | 1048 | 4.91E+06 |
|  | SS pCH + Kampy #2 5cm | 10<sup>-2</sup> | 3 | 1.52E+04 |
|  | SS pCH + Kampy #3 5cm | 10<sup>-1</sup> | 1022 | 6.19E+05 |
|  | NS pCH + Kampy #1 5cm | 10<sup>0</sup> | 15 | 9.26E+02 |
|  | NS pCH + Kampy #2 5cm | 10<sup>0</sup> | 37 | 1.92E+03 |
|  | NS pCH + Kampy #3 5cm | 10<sup>0</sup> | 70 | 3.91E+03 |
|  | SS pML + Kampy #1 5cm | 10<sup>-2</sup> | 77 | 4.31E+05 |
|  | SS pML + Kampy #2 5cm | 10<sup>-5</sup> | 812 | 4.63E+09 |
|  | SS pML + Kampy #3 5cm | 10<sup>-2</sup> | 1992 | 1.01E+07 |
|  | NS pML + Kampy #1 5cm | 10<sup>0</sup> | 66 | 4.07E+03 |
|  | NS pML + Kampy #2 5cm | 10<sup>0</sup> | 43 | 2.09E+03 |
|  | NS pML + Kampy #3 5cm | 10<sup>0</sup> | 24 | 1.44E+03 |
| Day 49 | Negative control | 10<sup>0</sup> | 0 | 0.00E+00 |
|  | SS Control #1 5cm | 10<sup>0</sup> | 3578 | 1.61E+05 |
|  | SS Control #2 5cm | 10<sup>0</sup> | 33 | 1.64E+03 |
|  | SS Control #3 5cm | 10<sup>0</sup> | 5000 | 2.83E+05 |
|  | NS Control #1 5cm | 10<sup>0</sup> | 8 | 3.90E+02 |
|  | NS Control #2 5cm | 10<sup>0</sup> | 14 | 7.76E+02 |
|  | NS Control #3 5cm | 10<sup>0</sup> | 407 | 2.00E+04 |
|  | SS pCH + Kampy #1 5cm | 10<sup>-3</sup> | 1540 | 7.59E+07 |
|  | SS pCH + Kampy #2 5cm | 10<sup>-2</sup> | 5000 | 3.08E+07 |
|  | SS pCH + Kampy #3 5cm | 10<sup>-2</sup> | 1948 | 1.01E+07 |
|  | NS pCH + Kampy #1 5cm | 10<sup>0</sup> | 88 | 5.10E+03 |
|  | NS pCH + Kampy #2 5cm | 10<sup>0</sup> | 14 | 7.69E+02 |
|  | NS pCH + Kampy #3 5cm | 10<sup>0</sup> | 0 | 0.00E+00 |
|  | SS pML + Kampy #1 5cm | 10<sup>-2</sup> | 2927 | 1.82E+07 |
|  | SS pML + Kampy #2 5cm | 10<sup>-6</sup> | 15 | 9.08E+08 |
|  | SS pML + Kampy #3 5cm | 10<sup>-3</sup> | 626 | 2.82E+07 |
|  | NS pML + Kampy #1 5cm | 10<sup>0</sup> | 72 | 3.58E+03 |
|  | NS pML + Kampy #2 5cm | 10<sup>0</sup> | 16 | 1.01E+03 |
|  | NS pML + Kampy #3 5cm | 10<sup>0</sup> | 12 | 5.95E+02 |

**Table S8:** Permutation *P*-values for statistically significant comparisons.

|  |  |  |  |  |  |  |  |  |
| --- | --- | --- | --- | --- | --- | --- | --- | --- |
| *M. smegmatis* colony count absolute abundance at inoculation site in only non-sterilized soil microcosms - *Ms*-pC3 vs. *Ms*-pML(int) ($P = 0.025$) | | | | | | | | |
| 0.07 | 0.86 | 0.56 | 0.0034 | 0.78 | 0.29 | 0.26 | 0.54 | 0.88 |
| *M. smegmatis* 16S relative abundance at inoculation site - *Ms*-pC3 vs. *Ms*-pML(int) ($P = 0.04$) | | | | | | | | |
| 0.23 | 0.77 | 0.08 | 0.41 | 0.96 | 0.14 | 0.86 | 0.59 | 0.34 |
| Kampy PFU/g dry soil at inoculation site - Sterile vs. Non-Sterile Soil ($P < 0.001$) | | | | | | | | |
| 0.09 | 0.09 | 0.09 | 0.57 | 0.57 | 0.57 | 0.09 | 0.09 | 0.09 |

**Table S9:** Percentage of pink *Ms*-pC3 colonies.

|  | #1 | #2 | #3 | Average |
| --- | --- | --- | --- | --- |
| Day 4, Sterilized Soil | 86% | 60% | N/A | 73% |
| Day 4, Non-Sterilized Soil | 100% | 75% | 100% | 92% |
| Day 7, Sterilized Soil | N/A | 70% | N/A | 70% |
| Day 7, Non-Sterilized Soil | 40% | 53% | N/A | 47% |
| Day 14, Sterilized Soil | 80% | 63% | N/A | 71% |
| Day 14, Non-Sterilized Soil | 67% | 88% | N/A | 77% |
| Day 49, Sterilized Soil | 8% | 73% | 0% | 27% |
| Day 49, Non-Sterilized Soil | N/A | N/A | 50% | 50% |

**Fig. S1:**
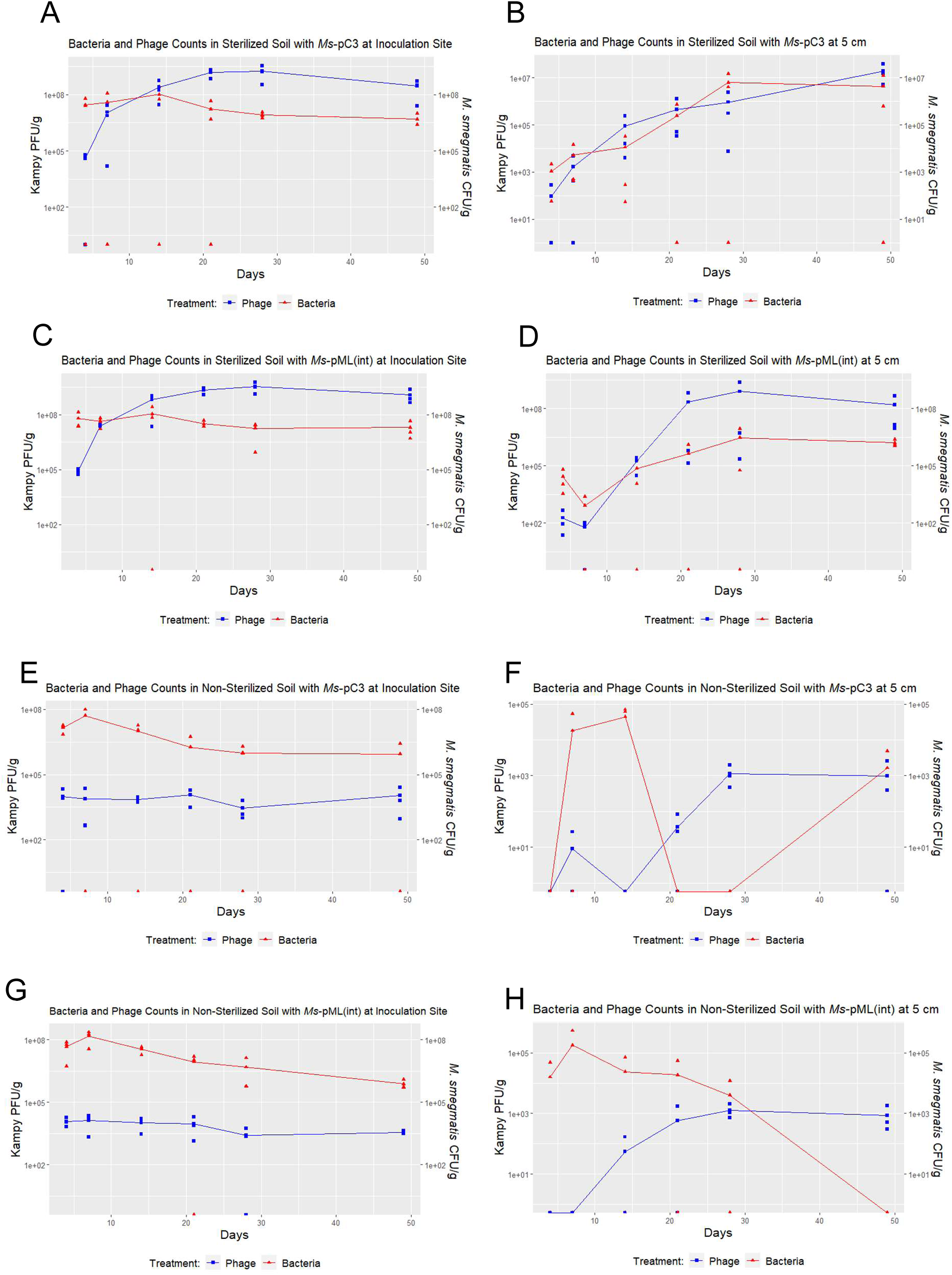
Graphs comparing bacteria and phage abundance within each microcosm.

**Fig. S2:**
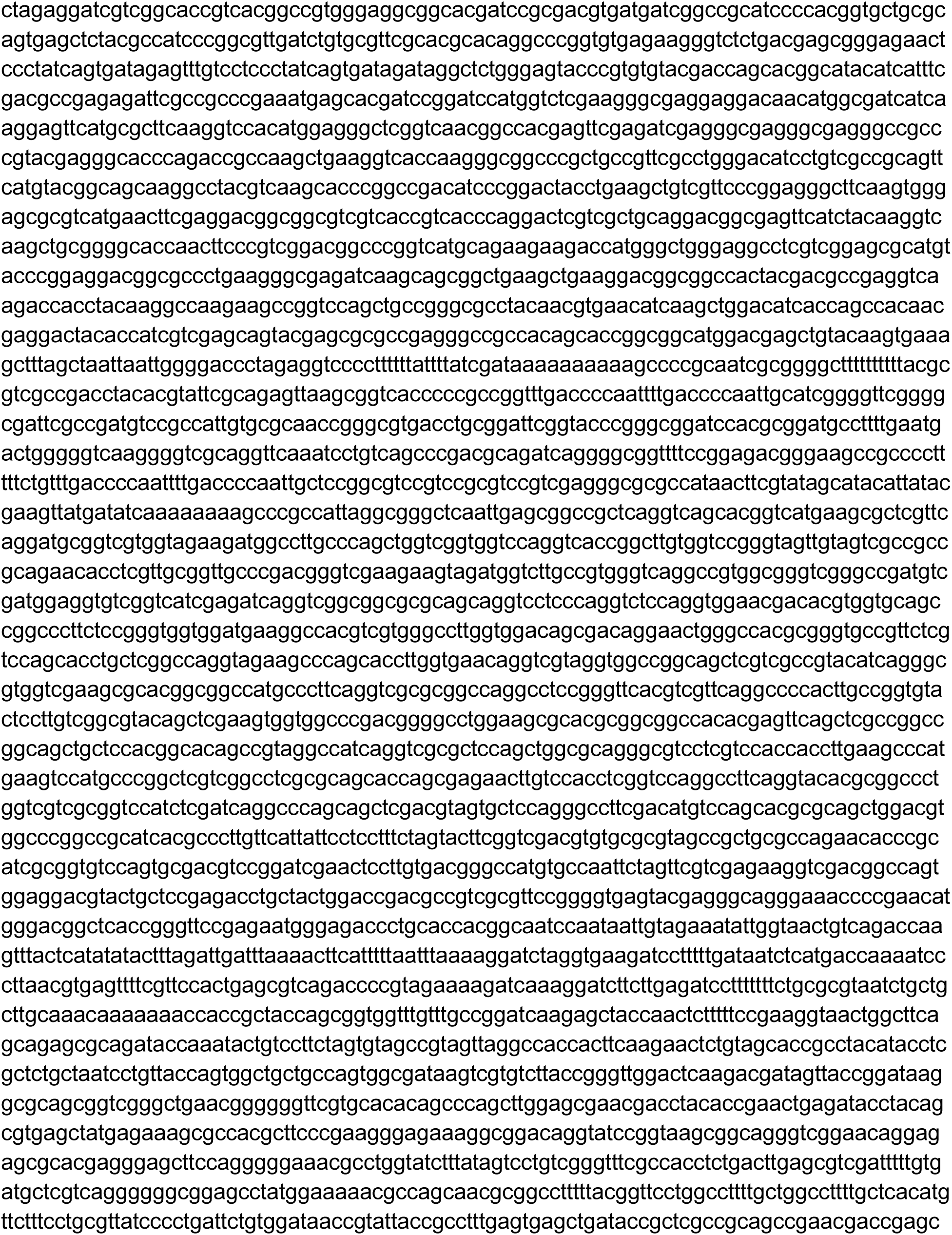
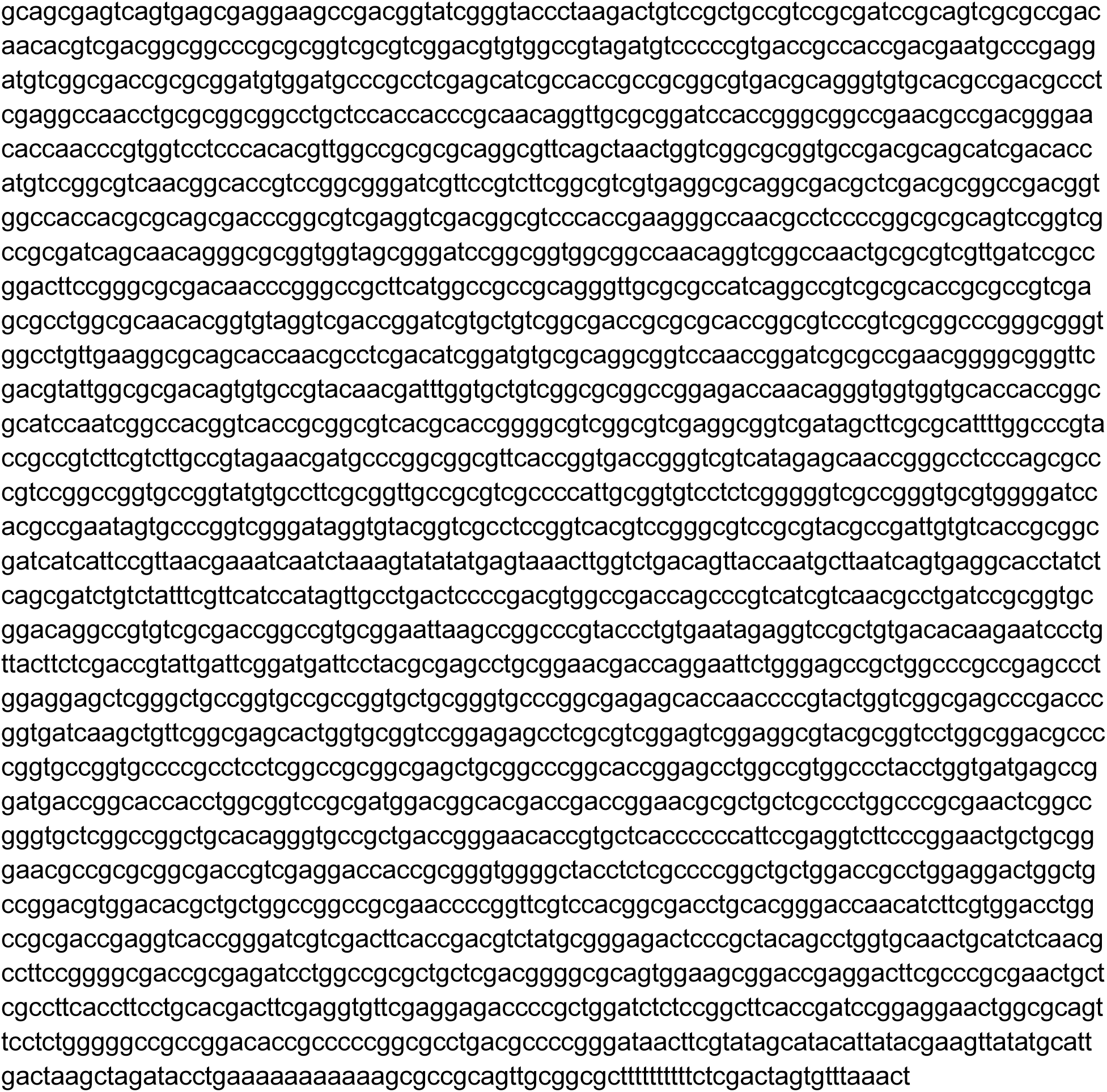
pMLcherry Plasmid Sequence

